# A PLM-Based Method for Predicting Protein Ion Channel Modulators for Drug Discovery and Safety Evaluation

**DOI:** 10.1101/2025.11.13.688159

**Authors:** Anand Singh Rathore, Saloni Jain, Naman Kumar Mehta, Gajendra P. S. Raghava

## Abstract

Ion channels are central to regulating neuronal communication, cardiac rhythm, and muscle contraction. Their modulation can induce therapeutic benefits but may also lead to adverse or toxic effects. This study presents IonNTXpred, a protein language model (PLM)-based method for predicting protein ion channel modulators, including channel-specific (Na□, K□, Ca²□, and others) and moonlighting proteins capable of modulating multiple ion channels. We train, test, and evaluate our models on the largest dataset of non-redundant ion channel–modulating proteins, where no two proteins have more than 40% sequence identity. Composition analysis revealed that residues Cys, Gly, and Trp are highly prevalent, whereas Ala, Glu, Leu, Gln, and Val are scarce in ion channel–modulating proteins, with Cys identified as a key discriminative residue. We explored both alignment-based (BLAST, MERCI) and alignment-free (machine learning, deep learning, and PLM-based) approaches. Among these, our evolutionary information–based PLM (ESM2-t33) achieved the best performance, with an AUROC of 0.97 across ion channels, which further improved to 0.98 when integrated with BLAST output. The proposed method outperformed existing approaches on independent datasets. We used IonNTXpred to screen FDA-approved and organismal proteins to identify candidates with ion channel–modulating potential, supporting drug repurposing, discovery of new therapeutic proteins, and safety assessment of existing biologics. We implemented these models in a user-friendly web server, IonNTXpred, which facilitates the design and discovery of ion channel–modulating protein-based drugs and supports the biosafety evaluation of therapeutic proteins through neurotoxin screening (https://webs.iiitd.edu.in/raghava/ionntxpred/).

**Highlights:**

- Prediction, design, and large-scale screening of ion channel–modulating proteins.
- Identifies neurotoxins to support safety evaluation of therapeutic proteins.
- Evolutionary information–based protein language model ESM2-t33 for prediction.
- Predicts moonlighting proteins capable of modulating multiple ion channels.
- Provides standalone software, a web server, and a PyPI package for user accessibility.

## 1 Introduction

Ion channels are essential membrane proteins that regulate the flow of ions across cell membranes, playing a critical role in various physiological processes. These proteins are fundamental to functions such as neuronal signaling, muscle contraction, cardiac activity, hormone secretion, immune responses, and epithelial transport of nutrients and electrolytes. Many ion channels function through a gating mechanism, which acts as a physical barrier to ion movement and controls cellular excitability (Ashcroft, 2006). Consequently, numerous therapeutic agents have been developed to modulate ion channel activity, leading to treatments for a wide range of conditions. Several therapeutic agents have been developed to modulate ion channel activity, leading to treatments for conditions such as epilepsy, chronic pain, diabetes, hypertension, cardiac arrhythmias, anxiety disorders, and autoimmune diseases. Ion channel modulators, including sodium (Na□), potassium (K□), and calcium (Ca²□) channel-targeting compounds, have been successfully used in clinical applications (Armijo et al., 2005; Nattel, 1999; Waszkielewicz et al., 2013). Dysregulation of ion channels has also been implicated in cancer progression, further expanding their relevance as potential oncology targets(Shi et al., 2025).

Animal toxins deploy a range of sophisticated mechanisms to regulate ion channel function, reflecting their evolutionary adaptation for prey immobilization and defense (Kalia et al., 2015). Pore-blocking toxins, such as ω-conotoxins and charybdotoxin, physically occlude the channel pore, thereby directly preventing ion passage across the membrane. In contrast, gating modifier toxins interact with voltage-sensing domains of channels, like δ-atracotoxin and hanatoxin, altering gating kinetics to stabilize channels in open or closed states, or disrupt the inactivation process, which can induce hyperexcitability in excitable cells (Lau, King, & Mobli, 2016). Allosteric modulators, exemplified by sea anemone toxins, bind at discrete non-pore sites, leading to conformational shifts that either facilitate or hinder channel opening. Ligand-competitive toxins, such as α-bungarotoxin, mimic or block neurotransmitter binding at ligand-gated channels, resulting in functional antagonism and neurological effects, including paralysis (Huang et al., 2013). Additionally, some toxins exert their influence by altering the membrane channel interface, subtly modulating gating behavior via changes in lipid-protein interactions a mechanism exploited by various spider toxins (Xu et al., 2019). Collectively, these diverse strategies underscore animal venom as a rich source of molecular probes for dissecting ion channel pharmacology and physiology. Figure 1 illustrates the principal mechanisms by which animal venom-derived ion channel toxins exert their modulatory effects, highlighting key toxin classes and representative examples within the context of cellular membrane physiology.

**Figure 1.**
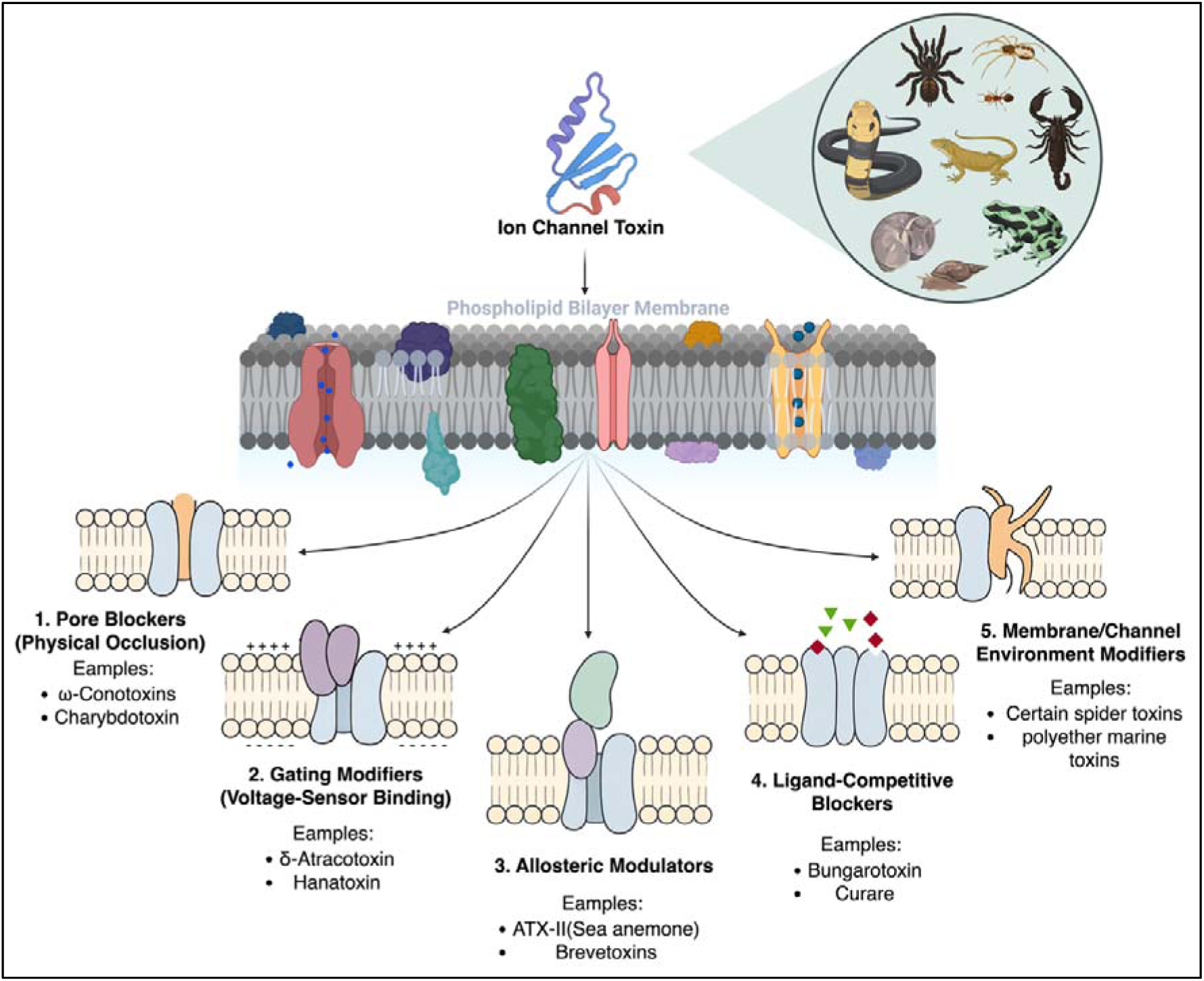
Schematic representation of major mechanisms by which ion channel toxins modulate channel function.

Traditionally, the identification and characterization of ion channel modulators have relied on various wet lab techniques. The manual patch-clamp technique, considered the gold standard, involves using a glass pipette to form a tight seal with the cell membrane, allowing for the measurement of ion channel currents (*Assay Guidance Manual*, 2004). Despite its precision, this method is labor-intensive and has low throughput. Fluorescence-based assays utilize voltage-sensitive dyes to detect changes in membrane potential or ion concentrations, offering higher throughput but providing indirect measurements of ion channel activity. Ion flux assays measure the movement of specific ions, such as using thallium (TlLJ) flux to assess potassium channel activity; however, they often lack the temporal resolution to capture rapid ion channel kinetics (Yu, Li, Wang, & Wang, 2016). Automated patch-clamp systems have been developed to address the limitations of manual techniques, increasing throughput and reducing variability, yet they remain costly and may not be accessible to all laboratories. While these traditional methods have significantly contributed to our understanding of ion channels, they present challenges in terms of scalability, cost, and efficiency (Zhu et al., 2022). While these methods have been invaluable, the exponential growth of protein sequence databases necessitates the development of high-throughput computational frameworks to accelerate the discovery process. Machine learning algorithms, in particular, offer a promising avenue for screening and identifying potential modulators from sequence data alone.

Over the past two decades, several machine learning-based tools have been developed for this purpose. However, many of these tools have notable limitations. Early work, such as NTXPred (2007) (Saha & Raghava, 2007b), pioneered the classification of neurotoxins and ion channel inhibitors using SVM-based models. Several subsequent predictors focused specifically on conotoxins, the venom peptides well known for targeting ion channels: the RBF Network (2013) (Yuan et al., 2013), iCTX-Type (2014) (Ding et al., 2014), and AVC-SVM (2017) (Xianfang, Junmei, Xiaolei, & Yue, 2017) were all designed to distinguish Na□, K□, and Ca²□ channel toxins within this family. In parallel, a few efforts aimed to develop general predictors that span multiple ion channel classes; examples include PrIMP (2022) (Lee et al., 2022), STACKION (2023) (Ali, Ahmed, Bui, & Chen, 2023), and the model proposed by Juan & Ji (2018), all of which attempt to capture shared molecular determinants of Na□, K□, Ca²□ and other channel modulators. More recently, several predictors have been tailored toward single-channel specificity: PEP-PREDNa□ (2022) (Herrera-Bravo, Farías, Contreras, Herrera-Belén, & Beltrán, 2022), NALL-Pred (2024), and MetaNaBP (2024) (Shoombuatong, Homdee, Schaduangrat, & Chumnanpuen, 2024) were designed exclusively for Na□ channel inhibitors, while PPLK^+^C (2019) (Lissabet, Belén, & Farias, 2020) was dedicated to potassium channel ligands. Together, these tools reflect the evolution of the field from early conotoxin-centered predictors, through broader multi-channel frameworks, to more refined channel-specific approaches. However, the predictive power of these earlier methods was often limited by their reliance on small, outdated datasets with high internal redundancy (using 80-90% sequence identity thresholds) and simple compositional features. Detailed information on existing tools related to the reduction of ion channel-impairing or modulating proteins is provided in Table 1.

**Table 1:** Overview of computational tools for predicting different types of ion channel modulating proteins.

| Tool (Year) | Methodology | Reference |
| --- | --- | --- |
| NTXpred<br>(2007) | SVM based model Neurotoxin classification and sub-classification of ion channel inhibitors (Na<sup>+</sup>, K<sup>+</sup>, Ca<sup>2+</sup>, Cl<sup>-</sup>) using compositional features. (https://webs.iitd.edu.in/raghava/ntxpred/) | (Saha & Raghava, 2007b) |
| RBF Network<br>(2013) | Radial Basis Function Neural Network model for Ion channel-targeted conotoxins. (http://cobi.uestc.edu.cn/people/hlin/data/conotoxin/) | (Yuan et al., 2013) |
| iCTX-Type<br>(2014) | Pseudo-amino acid composition, dipeptide occurrence based ML model for Ion channel-targeted conotoxins. (http://lingroup.cn/server/iCTX-Type) | (Ding et al., 2014) |
| AVC-SVM.<br>(2017) | Adaptive variable component SVM model for Ion channel-targeted conotoxins. | (Xianfang et al., 2017) |
| Juan & Ji<br>(2018) | Feature selection, pseudo-amino acid composition based model for Ion channel inhibitors. | (Mei, Fu, & Zhao, 2018) |
| PPLK<sup>+</sup>C<br>(2019) | Machine learning Model for Peptide ligands for potassium channels. (https://github.com/bio-coding/PPLKC-) | (Lissabet et al., 2020) |
| PEP-PRED<sup>Na+</sup><br>(2022) | Random forest Model for Voltage-gated Na<sup>+</sup> channel-blocking peptides. (https://www.biochemintelli.com/peppredna/) | (Herrera-Bravo et al., 2022) |
| PrIMP (2022) | Knowledge transfer approach for Ion channel (Na<sup>+</sup>, K<sup>+</sup>, Ca<sup>2+</sup> and nAChRs) modulating peptides. (https://github.com/bzlee-bio/PrIMP) | (Lee et al., 2022) |
| STACKION<br>(2023) | Machine learning model for predicting Ion channel (Na<sup>+</sup>, K<sup>+</sup> and Ca<sup>+</sup>) modulators. | (Ali et al., 2023) |
| NALL-Pred<br>(2024) | Ensemble model for Na<sup>+</sup> ion channel inhibitors prediction. (https://github.com/MirTanveer/NaII-Pred) | (Hassan, Tayara, & Chong, 2024) |
| MetaNaBP<br>(2024) | Meta-predictor, ensemble learning model for Voltage-gated Na<sup>+</sup> channel-blocking peptides. (https://github.com/Shoombuatong/MetaNaBP) | (Shoombuatong et al., 2024) |

Our group developed NTXpred, which was built on limited dataset available in 2007 using old machine learning techniques. In the past, most studies reused this small dataset for predicting the neurotoxicity of proteins (Herrera-Bravo et al., 2022; Shoombuatong et al., 2024). Recently, our group developed an updated version of NTxPred2, which is built on a large dataset using state-of-the-art machine learning techniques. One of the limitations of the NTXpred series is that it does not allow for the prediction of ion-channel-specific neurotoxins or ion-channel-modulating proteins. To address these gaps and overcome these limitations, in the present work we have constructed the largest and most up-to-date dataset of ion channel modulators, processed at 40% redundancy using CD-HIT to ensure non-redundant and diverse representation, while also encompassing sequences from earlier datasets to maintain continuity. Building on this resource, we developed a dedicated and comprehensive computational framework that leverages state-of-the-art (SOTA) large language models (LLMs) in combination with traditional machine learning methods to predict proteins that selectively modulate sodium, potassium, and calcium ion channels, while also accounting for their potential moonlighting properties. Additionally, we benchmarked our tool against existing methods to assess its effectiveness and identify areas for improvement. Figure 2 offers a visual overview of the algorithm and the procedures conducted within this study. This study addresses the growing need for reliable computational models in ion channel research, providing a high-throughput and data-driven solution to accelerate protein and peptide-based drug discovery and ion channel modulator identification.

**Figure 2:**
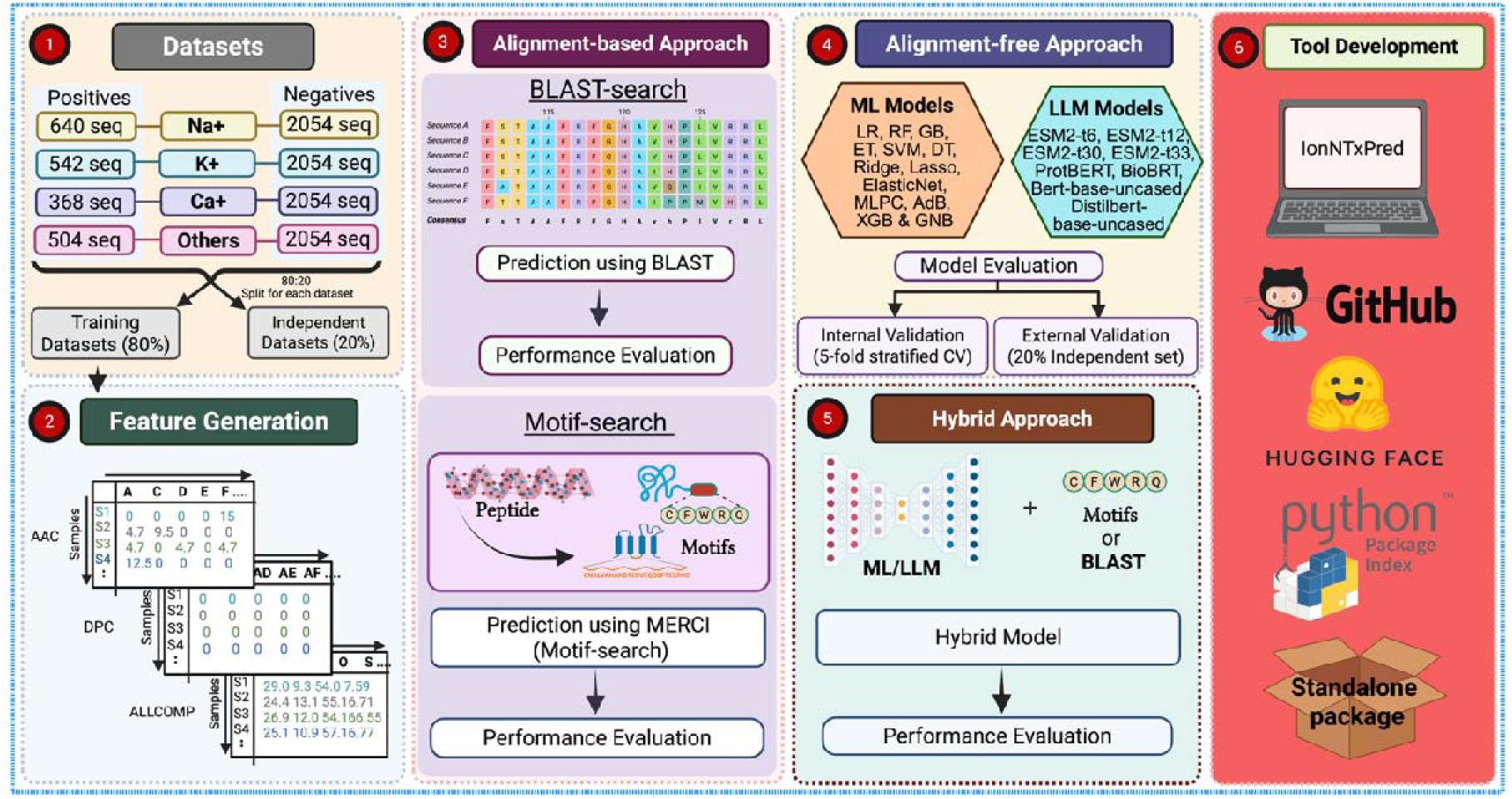
Flowchart illustrating the complete architecture of IonNTxPred.

## 2 Materials and Methods

### 2.1 Dataset

This section provides an overview of the datasets used in this study, including their compilation, preprocessing, and redundancy removal. Subsequent steps involved removing redundant sequences and identifying moonlighting ion channel toxins.

#### 2.1.1 Dataset Compilation and Preprocessing

Ion channel-impairing proteins were retrieved from the UniProtKB/Swiss-Prot Tox-Prot database (Jungo, Bougueleret, Xenarios, & Poux, 2012), using the query “KW-0872 reviewed:true.” This yielded 2,886 sequences, which were filtered to remove non-canonical amino acids, duplicates, and length outliers, resulting in 2,827 unique sequences. Based on functional annotations, these were classified into sodium (1,087), potassium (850), and calcium (593) channel modulators, with 817 sequences targeting other ion channels (e.g., chloride channels, ionotropic glutamate receptors, and acetylcholine receptors). Sequences affecting multiple channels were included in all relevant categories. This approach ensures that the models are trained on real-world examples of moonlighting toxins, enabling them to learn the shared features that confer multi-channel activity. For the negative dataset, sequences were collected from UniProt using the query “NOT toxin NOT neurotoxin reviewed: true.” The same preprocessing steps were applied, yielding an initial pool of 566,303 non-toxic (non-ion channel modulating) sequences.

#### 2.1.2 Creation of a Non-Redundant Dataset

To ensure model robustness and prevent data leakage, redundant sequences were removed. Previous studies have employed various sequence identity thresholds (80–90%) to maintain dataset integrity (Hassan et al., 2024; Lee et al., 2022; Yuan et al., 2013). While reducing redundancy limits the dataset size, it enhances generalizability. To address the class imbalance, we first applied CD-HIT (Li & Godzik, 2006) at a 90% sequence identity threshold, filtering highly similar sequences in each dataset individually. This resulted in 640 sodium (Na□), 542 potassium (K□), and 368 calcium (Ca²□) channel modulators. The remaining 504 sequences are included in other ion channel modulators.

To ensure both diversity and non-redundancy, we applied CD-HIT clustering at a 40% sequence identity threshold. This resulted in 115, 142, 102, and 110 clusters for Na□, K□, Ca²□, and other ion channel modulators, respectively. From these clusters, we constructed two non-overlapping datasets: a training set containing 80% of the sequences and an independent test set containing the remaining 20% of the sequences. Importantly, entire clusters were kept intact within either training or test sets, ensuring that no sequences from the same cluster appeared in both, thereby avoiding information leakage. The training set was further partitioned into five non-overlapping folds for cross-validation, again preserving cluster integrity and maintaining nearly equal sequence counts per fold. This cluster-based partitioning strategy provides a more realistic evaluation and has been widely adopted in earlier computational biology studies (Bendtsen, Jensen, Blom, Von Heijne, & Brunak, 2004; Gahlot, Choudhury, Bajiya, Kumar, & Raghava, 2024; Kaur, Arora, & Raghava, 2020; N. Sharma et al., 2021).

For the negative dataset, an ample number of non-modulating sequences allowed us to apply the same 40% identity threshold, selecting 2,054 representative sequences with a length distribution comparable to the positive dataset. This dataset was similarly split into training and independent sets, with training data fractionated into five folds and merged with positive training data.

#### 2.1.3 Moonlighting ion channel toxins

During dataset curation, we observed that a subset of peptides modulate more than one ion channel subtype (e.g., sodium and potassium, or calcium and sodium simultaneously). This functional promiscuity, often termed moonlighting activity, is biologically important: it reflects the evolutionary pressure on venom peptides to act as broad-spectrum modulators, enhancing prey immobilization and defense. A well-known example is Kurtoxin from the scorpion *Parabuthus transvaalicus*, which not only modifies the gating of voltage-gated sodium channels but also potently inhibits T-type calcium channels (Cav3.1/Cav3.2). Such moonlighting toxins complicate computational modeling, as the same sequence may appear in multiple functional categories, blurring the boundaries between channel classes. Rather than treating this as noise, we explicitly retained these sequences, since they provide a realistic representation of ion channel modulators in nature and highlight the need for predictive models that can accommodate multi-target activity.

### 2.2 Compositional and Positional Analysis

To identify key features distinguishing ion channel-impairing proteins, we conducted a detailed compositional and positional analysis across all four datasets. The study involved multiple statistical and sequence-based approaches to characterize amino acid composition and positional preferences. First, we analyzed amino acid composition within each class, calculating mean, median, and standard deviation of residue frequencies for both ion channel-modulating and non-modulating sequences. To determine statistically significant differences, we performed independent t-tests (*scipy.stats*), with false discovery rate (FDR) correction applied using the Benjamini-Hochberg procedure (*statsmodels.stats.multitest*), minimizing type I errors. To explore sequence-specific preferences, we utilized the Two Sample Logo (TSL) method, identifying residues that were significantly enriched or depleted at specific positions within protein sequences. This approach provided insights into residue distributions associated with neurotoxicity, highlighting potential sequence motifs contributing to ion channel modulation.

### 2.3 Alignment-based Approaches

#### 2.3.1 BLAST Search

A number of existing tools have primarily been tailored for identifying similarities or homologies between protein sequences. For sequence similarity evaluation and classification of ion channel-impairing proteins, we used an alignment-based method with Basic Local Alignment Search Tool (BLAST) (Altschul, Gish, Miller, Myers, & Lipman, 1990), release 2.9.0. We used the BLASTp version, which is specifically crafted to perform alignments on proteins sequences. This study was performed over four datasets: Na□, K□, Ca²□, and other ion channels. For each dataset, a BLAST database was prepared utilizing training datasets. Query sequences from either the training set or the independent test set were aligned against this repository to ascertain their classification based on sequence similarity. To investigate the effect of alignment stringency, we considered BLAST performance over a series of e-value thresholds from 10^2^ to 10^-12^. Upon training set assessment, we used the first non-self-alignment match to prevent bias from full matches to evaluate performance on the training dataset. For independent sets, classification was made according to the best alignment match: if the query sequence aligned most closely with ion channel modulating proteins, it was labeled as a modulator, and similarly for non-modulating sequences. This alignment-based strategy provided a consistent and interpretable method for identifying ion channel-modulating proteins based on known sequence homology and was systematically applied across all four datasets.

#### 2.3.2 Motif Search

Motifs are short, re-occurring amino acid patterns and are frequently predictive of the molecular interactions leading to toxicity. In motif-based search, we focus on identifying specific segments or patterns within proteins that are linked to channel modulator properties (Bailey, 2008). Detection of motifs allows not only for better model performance but also for mechanistic interpretation of proteins function. To extract functionally informative and class-specific motifs, we utilized the Motif-EmeRging and Classes-Identification (MERCI) software (Vens, Rosso, & Danchin, 2011), which detects conserved sub-sequences exclusive to a particular class. MERCI was executed with its standard Perl-based script using default parameters, with the gap setting fixed to zero (-g 0) to focus on contiguous, ungapped motifs. The number of motifs to extract (-fp) was changed through values (5, 10, 15, 20) to consider the impact of motif number on subsequent analysis. Each functional category was processed independently to identify channel-specific patterns. Furthermore, a classification scheme based on the Koolman-Röhm classification was explored to categorize motifs according to the physicochemical properties of constituent amino acids, including polarity, charge, hydrophobicity, aromaticity, and aliphatic characteristics. The motif analysis was conducted independently on four separate sets of ion channel-targeting proteins.

### 2.4 Feature Extraction

To generate a comprehensive feature set, we employed the *Pfeature* standalone tool [PMID: 36251780], which provides an extensive range of sequence-based descriptors. A total of 9,190 features were extracted for each protein, categorized into 18 distinct feature types. These include Amino Acid Composition (AAC), Dipeptide Composition (DPC), Atom Type Composition, Bond Type Composition, and a variety of physicochemical properties (PCP), among others. A detailed breakdown of all extracted features, along with their respective vector lengths, is provided in Supplementary Table S1.

### 2.5 Alignment-Free Approaches

#### 2.5.1 Machine Learning Models

Machine learning techniques play a pivotal role in differentiating ion channel-modulating proteins from non-modulating counterparts. In this study, we implemented and evaluated a diverse set of classification models using *scikit-learn* (sklearn), a widely adopted machine learning library. The classifiers employed include Logistic Regression (LR), Ridge, Lasso, ElasticNet, Random Forest (RF), ExtraTrees (ET), Gaussian Naïve Bayes (GNB), Decision Trees (DT), Multi-Layer Perceptron (MLP), XGBoost (XGB), AdaBoost (AdB), and Support Vector Classifier (SVC) with multiple kernel functions (linear, radial basis function (RBF), polynomial, and sigmoid). To enhance predictive performance, each model underwent systematic hyperparameter tuning, ensuring optimal classification accuracy and generalizability across datasets.

#### 2.5.2 Large Language Models

Large Language Models (LLMs) have transformed protein sequence analysis using transformer architectures to capture sequence patterns and structural properties. We utilized ESM2-t33, ESM2-t30, ESM2-t12, ESM2-t6, and BERT-based models like BioBERT, Prot-BERT, bert-base-uncased, and distilbert-base-uncased. ESM models use autoregressive transformers with self-attention, where ESM2-t33 (33 layers, ∼15B parameters) is the largest, and ESM2-t6 (6 layers, ∼8M parameters) is the most efficient (Lin et al., 2022, 2023). BERT-based models, using bidirectional transformers, process sequences simultaneously. BioBERT and PROT-BERT specialize in biomedical and protein data, while bert-base-uncased and distilbert-base-uncased (a compressed version) maintain general applicability. These models feature 12 transformer layers, 768 hidden units, and 12 attention heads. Hyperparameters were fine-tuned using *AdamW* optimization, with learning rates between 1e-4 and 5e-5, dropout (0.1–0.3), and linear decay schedules. The loss function was cross-entropy loss. Byte-pair encoding (BPE) for ESM models and *WordPiece* tokenization for BERT models ensured efficient representation. Gradient accumulation was applied to manage memory constraints in large models. Model selection prioritized AUC, computational feasibility, and inference speed.

### 2.6 Hybrid Approach

In this study, we developed a hybrid approach integrating motif-based analysis using MERCI with ML models and fine-tuned language models (PLMs) for the classification of channel-modulating proteins. To enhance predictive accuracy, we incorporated similarity-based methods such as BLAST and MERCI, which categorize classification sequences based on known patterns. A weighted scoring system was implemented, assigning +0.5 for positive predictions (channel modulating), -0.5 for negative predictions (non-modulating), and 0 for sequences with no matches. This weighting mechanism provides a quantitative measure of confidence, improving the reliability of the classification system.

### 2.7 *n*-Fold Cross-validation and Performance Metrics

To ensure a rigorous and unbiased evaluation of model performance, a systematic data partitioning strategy was employed. The dataset was randomly divided into two subsets: 80% for training and 20% for independent testing. The independent test set was strictly withheld from training, model selection, and hyperparameter tuning to provide an objective assessment of the final model’s performance. Within the training set, stratified five-fold cross-validation was implemented to prevent data leakage and ensure robust performance estimation. Stratification preserved the original class distribution in each fold, making it particularly beneficial for imbalanced datasets. This method provides a more reliable performance estimate than standard k-fold cross-validation. Model performance was assessed using threshold-independent and threshold-dependent metrics. The area under the receiver operating characteristic curve (AUC-ROC) served as a threshold-independent metric, while accuracy, sensitivity, specificity, and the Matthews correlation coefficient (MCC) were used as threshold-dependent measures. These metrics have been widely validated in prior studies (Rathore, Choudhury, Arora, Tijare, & Raghava, 2024; Saha & Raghava, 2007b) and are crucial for evaluating the effectiveness of classification models in distinguishing channel-modulating proteins from non-modulating ones.

## 3 Results

This section summarizes the compositional and positional analyses of ion channel modulators and evaluates predictive performance using both alignment-based (BLAST, MERCI) and alignment-free (machine learning, PLM) approaches, highlighting key trends and comparative outcomes.

### 3.1 Analysis

Analysis encompasses detailed compositional and positional analyses, examining amino acid frequencies, residue preferences, and the discriminative power of key residues such as cysteine. It further includes logistic regression modeling and probability-based assessments to quantify residue contributions, providing foundational insights into sequence determinants of ion channel modulation.

#### 3.1.1 Compositional Analysis

The amino acid profiles of channel-modulating and non-modulating proteins were analyzed, and a comparative assessment of their average amino acid compositions was conducted. Figure 3 illustrates the average AAC of sodium, potassium, calcium, and other channel toxins compared to non-toxic proteins. Significant compositional differences are evident across several residues. In particular, cysteine (C), a polar and uncharged amino acid, is markedly enriched in ion channel-modulating proteins. Additionally, toxin classes exhibit higher proportions of residues such as glycine (G), lysine (K), tyrosine (Y), and tryptophan (W), while residues like isoleucine (I), glutamic acid (E), phenylalanine (F), and glutamine (Q) are less abundant. These patterns suggest that specific amino acids may be critical for conferring toxicity or determining channel specificity. To further elucidate these differences, we conducted a comparative analysis of AAC between positive (toxic) and negative (non-toxic) groups using Welch’s t-test, as detailed in Supplementary Figure S1. Statistical analysis revealed that the compositional differences for all amino acids, except asparagine (N), proline (P), and threonine (T), were significant at a corrected p-value threshold of 0.001, underscoring the distinct structural and functional characteristics of ion channel toxins.

**Figure 3:**
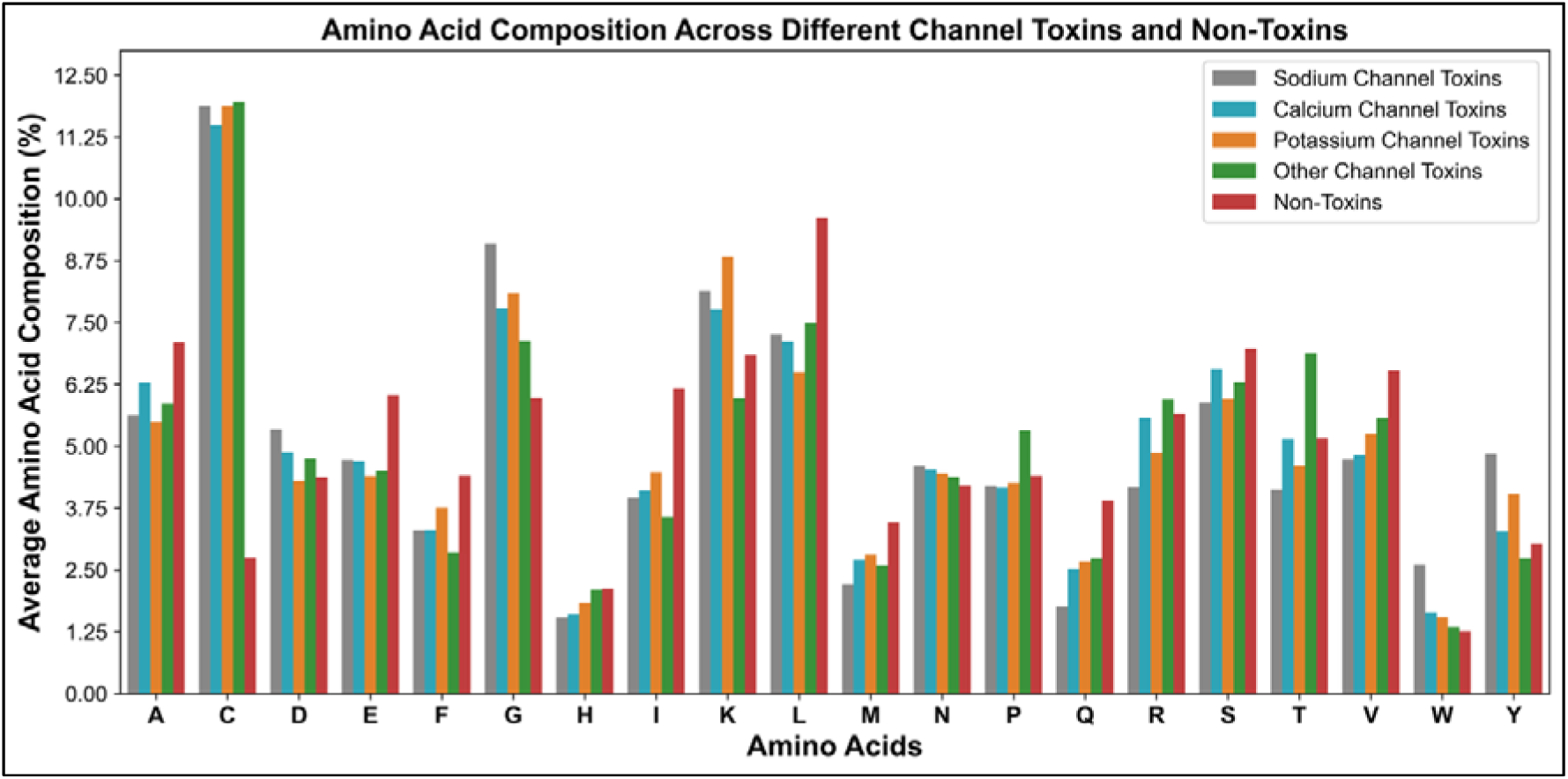
Average amino acid composition of ion-channel modulating and non-toxin sequences.

#### 3.1.2 Positional Analysis

We employed the Two Sample Logo (TSL) method to compare the positional amino acid preferences between ion channel-modulating and non-modulating proteins, analyzing both overall trends and individual ion channel types. Figure 4A illustrates the first 10 N-terminal residues, while Figure 4B depicts the last 10 C-terminal residues. In general, channel-modulating proteins exhibit a pronounced enrichment of residues such as lysine (K), leucine (L), valine (V), threonine (T), and cysteine (C) at the N-terminus, with cysteine becoming particularly dominant at the C-terminus. In contrast, non-toxic proteins tend to feature methionine at the N-terminus and leucine at the C-terminus. Analysis of individual ion channel modulators (Supplementary Figure S2) further reveals distinct patterns: sodium channel modulators prefer glycine (G), K, L, isoleucine (I), methionine (M), serine (S), and C at the N-terminal, with C, along with tryptophan (W) and K, frequently occurring at the C-terminal; potassium channel modulators show a preference for G, K, L, C, and phenylalanine (F) at the N-terminal, with C, proline (P), and K in the C-terminal region; calcium channel modulators favor G, K, L, T, C, V, I, and alanine (A) at the N-terminal, while the C-terminal is dominated by C, with occasional enrichment of G, arginine (R), and K; and other channel modulators are enriched for L, K, C, T, and V at the N-terminal, with C predominating the C-terminal along with T and R in select positions. Overall, the consistent predominance of cysteine at the C-terminus across all channel-modulating proteins suggests its critical role in conferring structural stability and functional specificity.

**Figure 4:**
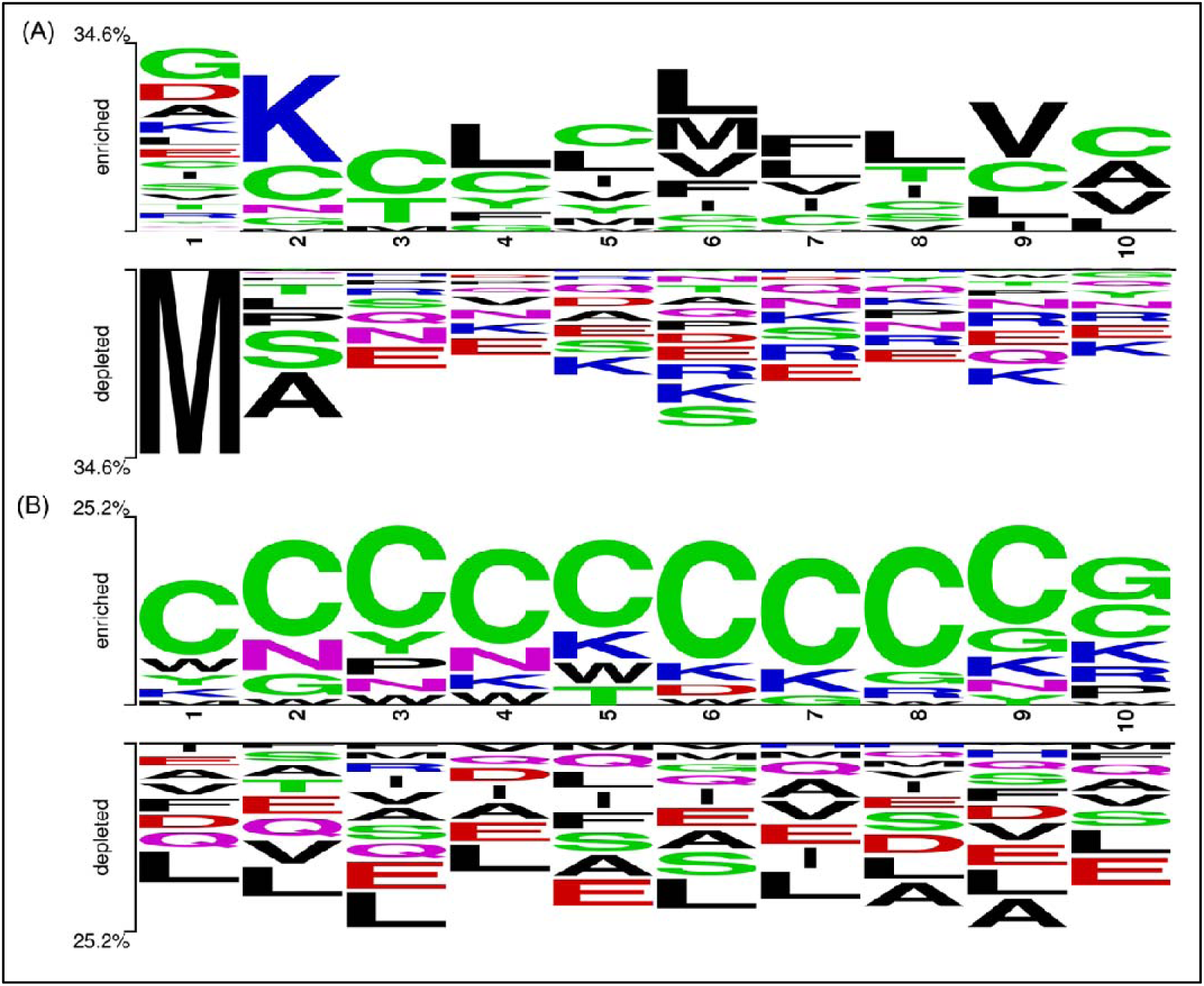
**Residue preferences at different positions in ion channel modulators and non-modulator sequences, as revealed by the Two Sample Logo (TSL) method. (A) Shows the first 10 residues at the N-terminus, and (B) shows the last 10 residues at the C-terminus.**

#### 3.1.3 Amino Acid Composition-based Prediction

To assess the discriminative power of individual amino acids in differentiating ion channel-modulating from non-modulating sequences, we systematically evaluated AAC across the four datasets (NaLJ, KLJ, Ca²LJ, and Other). Among all residues, cysteine (AAC_C) consistently emerged as the most powerful predictor (Table 2). Across datasets, cysteine composition was markedly higher in modulating proteins compared to non-modulating ones, reflecting its strong association with ion channel activity. This trend is likely explained by its structural role in forming disulfide bridges that stabilize cysteine-rich scaffolds, a hallmark of many ion channel toxins.

**Table 2.** Predictive performance (threshold, ACC, AUC) of individual amino acid compositions for classifying ion channel modulating proteins across Na. D**, K**D**, Ca²**D**, and Other datasets.**

| Amino Acid | Na<sup>+</sup> |  |  | K<sup>+</sup> |  |  | Ca<sup>2+</sup> |  |  | Other |  |  |
| --- | --- | --- | --- | --- | --- | --- | --- | --- | --- | --- | --- | --- |
|  | Best Threshold | ACC | AUC | Best Threshold | ACC | AUC | Best Threshold | ACC | AUC | Best Threshold | ACC | AUC |
| A | 6.364 | 0.537 | 0.596 | 6.298 | 0.531 | 0.597 | 6.692 | 0.486 | 0.529 | 6.484 | 0.520 | 0.580 |
| C | 7.312 | 0.889 | 0.934 | 7.312 | 0.879 | 0.932 | 7.124 | 0.869 | 0.910 | 7.354 | 0.877 | 0.932 |
| D | 4.856 | 0.589 | 0.599 | 4.337 | 0.464 | 0.505 | 4.627 | 0.566 | 0.552 | 4.559 | 0.558 | 0.554 |
| E | 5.380 | 0.531 | 0.587 | 5.218 | 0.548 | 0.616 | 5.370 | 0.522 | 0.595 | 5.269 | 0.536 | 0.614 |
| F | 3.852 | 0.524 | 0.603 | 4.081 | 0.478 | 0.545 | 3.854 | 0.504 | 0.591 | 3.632 | 0.546 | 0.644 |
| G | 7.535 | 0.698 | 0.737 | 7.034 | 0.648 | 0.669 | 6.884 | 0.628 | 0.652 | 6.553 | 0.593 | 0.616 |
| H | 1.834 | 0.498 | 0.585 | 1.977 | 0.455 | 0.535 | 1.865 | 0.468 | 0.567 | 2.116 | 0.435 | 0.505 |
| I | 5.069 | 0.587 | 0.664 | 5.320 | 0.547 | 0.625 | 5.139 | 0.562 | 0.654 | 4.873 | 0.600 | 0.700 |
| K | 7.493 | 0.602 | 0.604 | 7.839 | 0.620 | 0.629 | 7.306 | 0.590 | 0.564 | 6.407 | 0.491 | 0.544 |
| L | 8.438 | 0.585 | 0.650 | 8.055 | 0.619 | 0.693 | 8.366 | 0.583 | 0.654 | 8.552 | 0.566 | 0.622 |
| M | 2.839 | 0.575 | 0.661 | 3.135 | 0.532 | 0.579 | 3.079 | 0.541 | 0.595 | 3.028 | 0.550 | 0.620 |
| N | 4.403 | 0.581 | 0.556 | 4.326 | 0.572 | 0.540 | 4.368 | 0.582 | 0.550 | 4.292 | 0.561 | 0.525 |
| P | 4.297 | 0.469 | 0.499 | 4.332 | 0.456 | 0.500 | 4.280 | 0.453 | 0.497 | 4.859 | 0.604 | 0.582 |
| Q | 2.836 | 0.612 | 0.730 | 3.291 | 0.545 | 0.617 | 3.217 | 0.552 | 0.630 | 3.323 | 0.540 | 0.611 |
| R | 4.912 | 0.552 | 0.607 | 5.261 | 0.503 | 0.541 | 5.614 | 0.463 | 0.484 | 5.798 | 0.555 | 0.540 |
| S | 6.429 | 0.536 | 0.579 | 6.464 | 0.527 | 0.570 | 6.770 | 0.494 | 0.529 | 6.636 | 0.507 | 0.537 |
| T | 4.645 | 0.554 | 0.601 | 4.885 | 0.509 | 0.552 | 5.159 | 0.460 | 0.491 | 6.023 | 0.639 | 0.629 |
| V | 5.628 | 0.571 | 0.642 | 5.889 | 0.542 | 0.597 | 5.674 | 0.565 | 0.629 | 6.050 | 0.513 | 0.558 |
| W | 1.935 | 0.725 | 0.700 | 1.407 | 0.572 | 0.517 | 1.452 | 0.598 | 0.545 | 1.304 | 0.553 | 0.521 |
| Y | 3.942 | 0.666 | 0.643 | 3.537 | 0.612 | 0.599 | 3.158 | 0.549 | 0.529 | 2.887 | 0.495 | 0.531 |

(Abbreviations: ACC: Accuracy; AUC: Area Under the Receiver Operating Characteristic Curve)

In terms of predictive performance, AAC_C achieved high accuracies (0.869–0.889) and AUC values (>0.91) across all four datasets, outperforming other amino acids. Error analysis further highlighted its robustness: despite the strong separation, only a small fraction of sequences were misclassified when using AAC_C thresholds (Table 3). For instance, in the Na□ dataset (N=2694), 8.24% of sequences were false positives (non-modulating but above the threshold) and 2.86% false negatives (modulating but below the threshold). Similar proportions were observed for K□ (FP: 8.55%, FN: 3.58%), Ca²□ (FP: 9.83%, FN: 3.26%), and Other datasets (FP: 8.68%, FN: 3.56%). Importantly, very few misclassified sequences exceeded the neurotoxic mean (AAC_C > 11.8; ∼1.7–1.9% of total) or fell below the non-toxic mean (AAC_C < 2.75; ∼0.3–0.7%). This indicates that most errors occur near the decision boundary, reinforcing the discriminative reliability of cysteine.

**Table 3:** Performance of amino acid composition of cystine-based prediction across four ion channel modulator datasets.

| Dataset | Mean (Modulator) | Mean (Non-modulator) | Best Threshold | TP | TN | FP | FN | FP > 11.8 | FN < 2.75 |
| --- | --- | --- | --- | --- | --- | --- | --- | --- | --- |
| Na<sup>+</sup> | 11.88 | 2.75 | 7.312 | 563<br>(20.9%) | 1832<br>(68.0%) | 222<br>(8.24%) | 77<br>(2.86%) | 47<br>(1.74%) | 18<br>(0.67%) |
| K<sup>+</sup> | 11.88 | 2.75 | 7.312 | 449<br>(17.3%) | 1832<br>(70.6%) | 222<br>(8.55%) | 93<br>(3.58%) | 47<br>(1.81%) | 8<br>(0.31%) |
| Ca<sup>2+</sup> | 11.5 | 2.75 | 7.124 | 289<br>(11.9%) | 1816<br>(75.0%) | 238<br>(9.83%) | 79<br>(3.26%) | 47<br>(1.94%) | 14<br>(0.58%) |
| Other | 11.96 | 2.75 | 7.354 | 413<br>(16.1%) | 1832<br>(71.6%) | 222<br>(8.68%) | 91<br>(3.56%) | 47<br>(1.84%) | 11<br>(0.43%) |

(Abbreviations: TP: True Positive, TN: True Negative, FP: False Positive, FN: False Negative)

Other amino acids also provided meaningful channel-specific signals. Glycine (G), which imparts flexibility to peptide backbones, demonstrated strong discriminative power in Na□ (ACC: 0.698, AUC: 0.737) and K□ (ACC: 0.648, AUC: 0.669). Aromatic residues such as tryptophan (W) and tyrosine (Y), known for π–π stacking and hydrophobic interactions (McGaughey, Gagné, & Rappé, 1998), showed moderate yet consistent predictive contributions, with W performing best in Na□ (AUC: 0.700) and Y in Na□ and K□ datasets. Leucine (L) exihibited notable predictive strength in the K□ dataset (ACC: 0.619, AUC: 0.693), likely reflecting hydrophobic core stabilization or transmembrane interactions specific to potassium channel toxins. Alanine (A), in contrast, was weakly discriminative (AUC: 0.53–0.59), serving as a baseline for low-informative residues. Additional trends included threonine (T) and isoleucine (I), which were more relevant in the Other dataset (T AUC: 0.629; I AUC: 0.700), and glutamine (Q), which had higher predictive value in the Na□ dataset (AUC: 0.730).

By contrast, residues such as proline (P), asparagine (N), and histidine (H) exhibited poor discriminative performance across all datasets, indicating limited roles in ion channel modulation. Collectively, these results emphasize that AAC, particularly cysteine enrichment, is a defining feature of ion channel-modulating peptides. While certain residues provide channel-specific signals, cysteine’s consistent and robust predictive strength across all datasets highlights its critical role in shaping both structure and function. More detailed results are provided in Supplementary Table S2.

### 3.1.4 Probability Trends with Cysteine Content

To further investigate the role of cysteine in differentiating modulators from non-modulators, we analyzed how the probability of classification changes across varying cysteine content in the four datasets (Na□, K□, Ca²□, and Other). As shown in Figure 5, the probability of sequences being predicted as modulators increased sharply with higher cysteine composition, while the probability of non-modulators decreased correspondingly. A crossover point was consistently observed around 8–10% cysteine in the Na□, K□, and Other datasets, whereas in the Ca²□ dataset this shift occurred slightly later, around 10–12% cysteine. Beyond these thresholds, modulators became markedly more probable than non-modulators across all datasets. This trend was stable across all datasets, highlighting cysteine as a critical compositional determinant of ion channel modulation. The observed shift is consistent with the structural necessity of cysteine residues in forming disulfide bonds, which stabilize the conformations required for effective interaction with ion channels. These findings not only reinforce the discriminative power of cysteine but also emphasize the importance of residue-level compositional thresholds in predicting functional activity.

**Figure 5:**
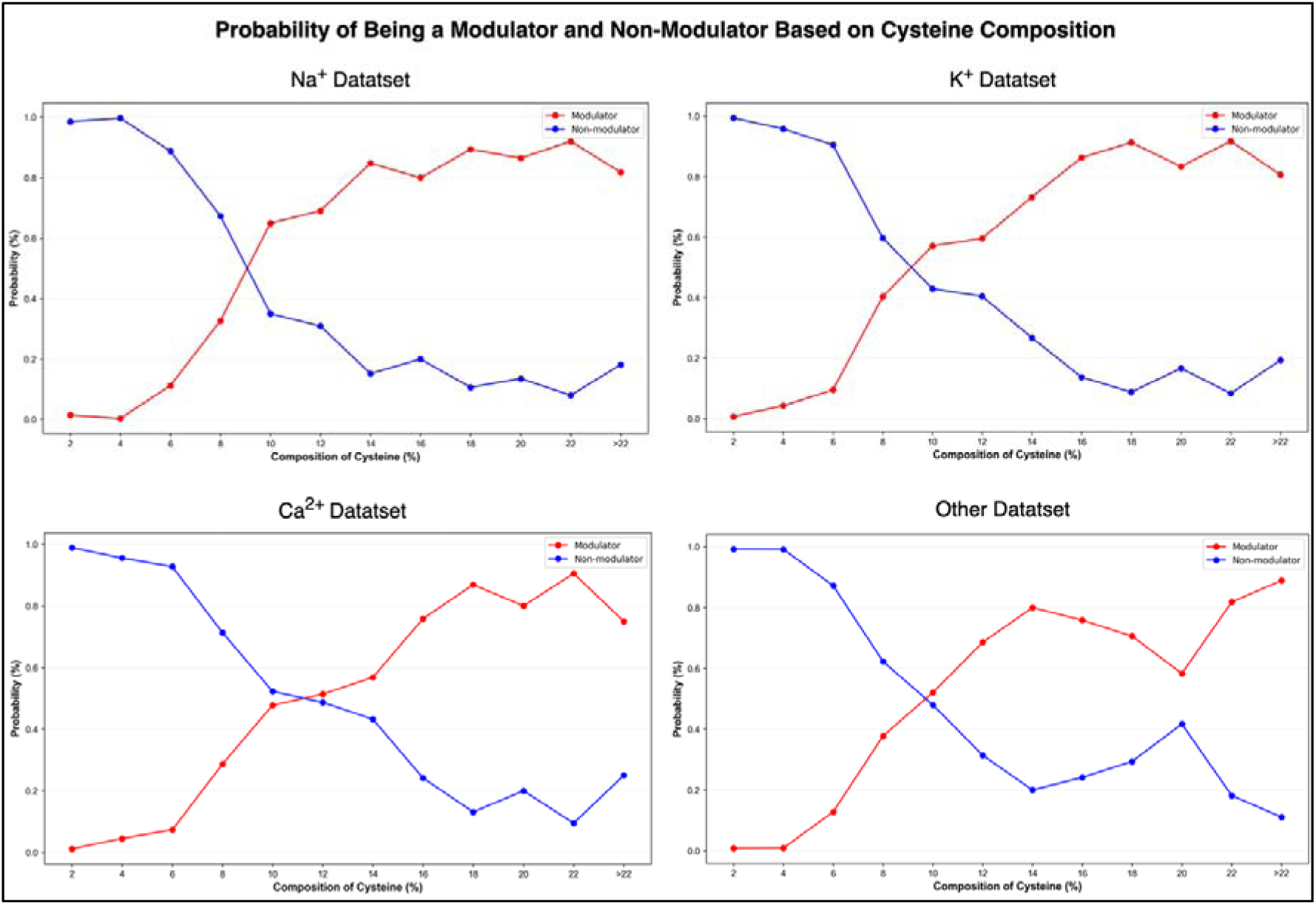
Probability trends of modulator and non-modulator classification across varying cysteine composition in Na□, K□, Ca^2^□, and Other datasets.

#### 3.1.5 Logistic Regression–Based Prediction Models

To investigate the predictive contribution of AAC features, we initially trained logistic regression models using a single feature (AAC_C, cysteine composition) across all four datasets (Na^+^, K^+^, Ca^2+^, and Others). Although single-feature models captured basic probability trends, their predictive performance was limited. Therefore, we systematically explored combinations of AAC features to enhance model accuracy. The final models, obtained through regularized logistic regression, incorporated multiple features and demonstrated improved discrimination between modulators and non-modulators.

The logistic regression model is expressed as:

where *x_i_* represents AAC features, *β_i_* their corresponding coefficients, and p (x) the probability of a sequence being an ion channel modulator.

Logistic regression models were developed for each ion channel–specific dataset using cysteine composition (AAC_C) along with additional AAC features, and their performance was evaluated through cross-validation and independent testing. For the Na+ dataset (AAC_C, AAC_Q, AAC_S), the model achieved high accuracy (91% cross-validation, 87% independent) with strong sensitivity (0.93 cross-validation, 0.90 independent) and AUC values of 0.95 and 0.94 (Table 4). In the K+ dataset (AAC_C, AAC_I, AAC_Y), performance was slightly lower, with accuracy of 88% (CV) and 84% (independent) and AUC of 0.94 (CV) and 0.89 (test), reflecting reduced specificity. The Ca2+ model (AAC_C, AAC_I, AAC_N) maintained robust performance with accuracy of 89% (cross-validation and independent) and AUC values of 0.92 and 0.90, respectively. The “Other” dataset (AAC_C, AAC_I, AAC_F) showed the strongest generalization, yielding accuracy of 89% (cross-validation) and 86% (independent) with AUC values of 0.95 and 0.91. Overall, all models achieved AUCs above 0.89 with consistently high sensitivity (>0.86) and precision (>0.93), underscoring the central importance of cysteine composition across datasets, while other residues (Q, S, I, Y, F, N) provided dataset-specific refinements.

**Table 4:** Evaluation metrics of logistic regression models for Na+, K+, Ca2+and Other datasets based on selected AAC features.

| Dataset | Features Used | Data Split | SENS | SPEC | PPV | ACC | MCC | AUC |
| --- | --- | --- | --- | --- | --- | --- | --- | --- |
| Na<sup>+</sup> | AAC_C, AAC_Q, AAC_S | Cross-validation | 0.934 | 0.826 | 0.948 | 0.910 | 0.746 | 0.950 |
|  |  | Independent | 0.895 | 0.755 | 0.934 | 0.866 | 0.615 | 0.942 |
| K<sup>+</sup> | AAC_C, AAC_I, AAC_Y | Cross-validation | 0.902 | 0.767 | 0.951 | 0.880 | 0.611 | 0.944 |
|  |  | Independent | 0.866 | 0.671 | 0.942 | 0.839 | 0.458 | 0.893 |
| Ca<sup>2+</sup> | AAC_C, AAC_I, AAC_N | Cross-validation | 0.906 | 0.714 | 0.968 | 0.888 | 0.504 | 0.924 |
|  |  | Independent | 0.912 | 0.712 | 0.963 | 0.891 | 0.534 | 0.901 |
| Other | AAC_C, AAC_I, AAC_F | Cross-validation | 0.918 | 0.767 | 0.951 | 0.893 | 0.645 | 0.950 |
|  |  | Independent | 0.897 | 0.671 | 0.932 | 0.859 | 0.531 | 0.910 |

(Abbreviations: SENS: Sensitivity; SPEC: Specificity; PPV: Positive Predictive Value; ACC: Accuracy; MCC: Matthews Correlation Coefficient; AUC: Area Under the Receiver Operating Characteristic Curve; AAC\_C (Cysteine), AAC\_Q (Glutamine), AAC\_S (Serine), AAC\_I (Isoleucine), AAC\_Y (Tyrosine), AAC\_N (Asparagine), AAC\_F (Phenylalanine))

**Final Predictive Equations**

The fitted logistic regression formulas for each dataset are:

- Na^+^: *logit* (*p*) = -2.4367 + 0.4316 • AAC_C - 0.3740 • AAC_Q - 0.1072 • AAC_S
- K^+^: *logit* (*p*) = -4.3269 + 0.4024 • AAC_C - 0.0210 • AAC_I+ 0.1306 • AAC_Y
- Ca^2+^: *logit* (*p*) = -3.5594 + 0.3513 • AAC_C - 0.0994 • AAC_I+ 0.0490 • AAC_N
- Other: *logit* (*p*) = -2.8775 + 0.4174 • AACC - 0.1963 • AAC_I- 0.0874 • AAC_F

These equations can be directly applied to estimate the probability p (x) for new protein sequences, providing interpretable models grounded in AAC.

### 3.2 Alignment-based methods

This evaluates sequence similarity and motif-based approaches, employing BLAST to assess homology-driven classification and MERCI to identify conserved amino acid patterns of ion channel modulators.

#### 3.2.1 Performance of BLAST-search

To compare the performance of sequence similarity-based classification on novel data, BLAST searches were conducted between the training and independent datasets at different E-value cut-offs from 10^2^ to 10^-12^ for each category of channel modulators. BLAST performance, in terms of alignment reliability, is inversely correlated with the E-value, as higher E-values increase the likelihood of random matches. The BLAST search results on the independent dataset (detailed in Supplementary Table S3) reveal that the approach is highly accurate in detecting ion channel modulators, with very low error rates. However, the coverage of BLAST is very limited. Although the BLAST-based approach is highly sensitive, its overall predictive performance is limited by overall coverage.

Across all four independent datasets (Na□, K□, Ca²□, and Other), BLAST similarity search results show a clear trend based on the E-value threshold. At the most relaxed E-value of 100, BLAST achieved nearly 100% coverage in each case (Na□: 539 hits, K□: 520 hits, Ca²□: 485 hits, Other: 512 hits). However, this came with reduced accuracy, only 77.06–79.07% of modulator hits and 85.41–89.30% of non-modulator hits were correct, indicating a rise in false positives. In contrast, at the most stringent E-value of 10^-12^, coverage dropped substantially (Na□: 9.46%, K□: 8.27%, Ca²□: 5.77%, Other: 6.64%), but accuracy was excellent: 88.89–100% of modulator hits and 95.00–100% of non-modulator hits were correct. At a moderate threshold of 10^-3^, the datasets demonstrated a favourable balance between sensitivity and specificity, coverage ranged from 27.63% (Ca²□) to 33.77% (Na□), while modulator hit accuracy was between 93.67–97.56% and non-modulator accuracy exceeded 99% in most cases. These results suggest that an E-value around 10^-3^ offers the best trade-off between broad sequence identification and high classification precision.

#### 3.2.2 Motif-based Prediction

To evaluate the contribution of sequence motifs in predicting ion channel modulators, we applied MERCI-based motif analysis across four independent datasets (Na□, K□, Ca²□, and Other ions) using both default and Koolman-Rohm configurations. The top 10 motifs obtained from each configuration are presented in Table 5. Based on the motif analysis across four independent datasets, we identified the best-performing MERCI configurations in terms of highest classification accuracy. In the Na□ dataset, the Koolman-Rohm configuration (fp 5, top 50 motifs) showed the highest accuracy of 87.21%, effectively balancing true positives and minimizing false positives. For the K□ dataset, the best result was obtained using default MERCI (fp 5, top 100 motifs) with an accuracy of 86.25%, slightly outperforming Koolman-Rohm options. In the Ca²□ dataset, Koolman-Rohm configuration (fp 5, top 50 motifs) led with the best accuracy of 89.58%, demonstrating its strength in detecting relevant patterns. Finally, in the Other dataset, both default MERCI (fp 5, top 100) and Koolman-Rohm (fp 5, top 100) configurations performed comparably, with accuracies of 86.01% and 83.70%, respectively, making default MERCI slightly better in this case.

**Table 5:**
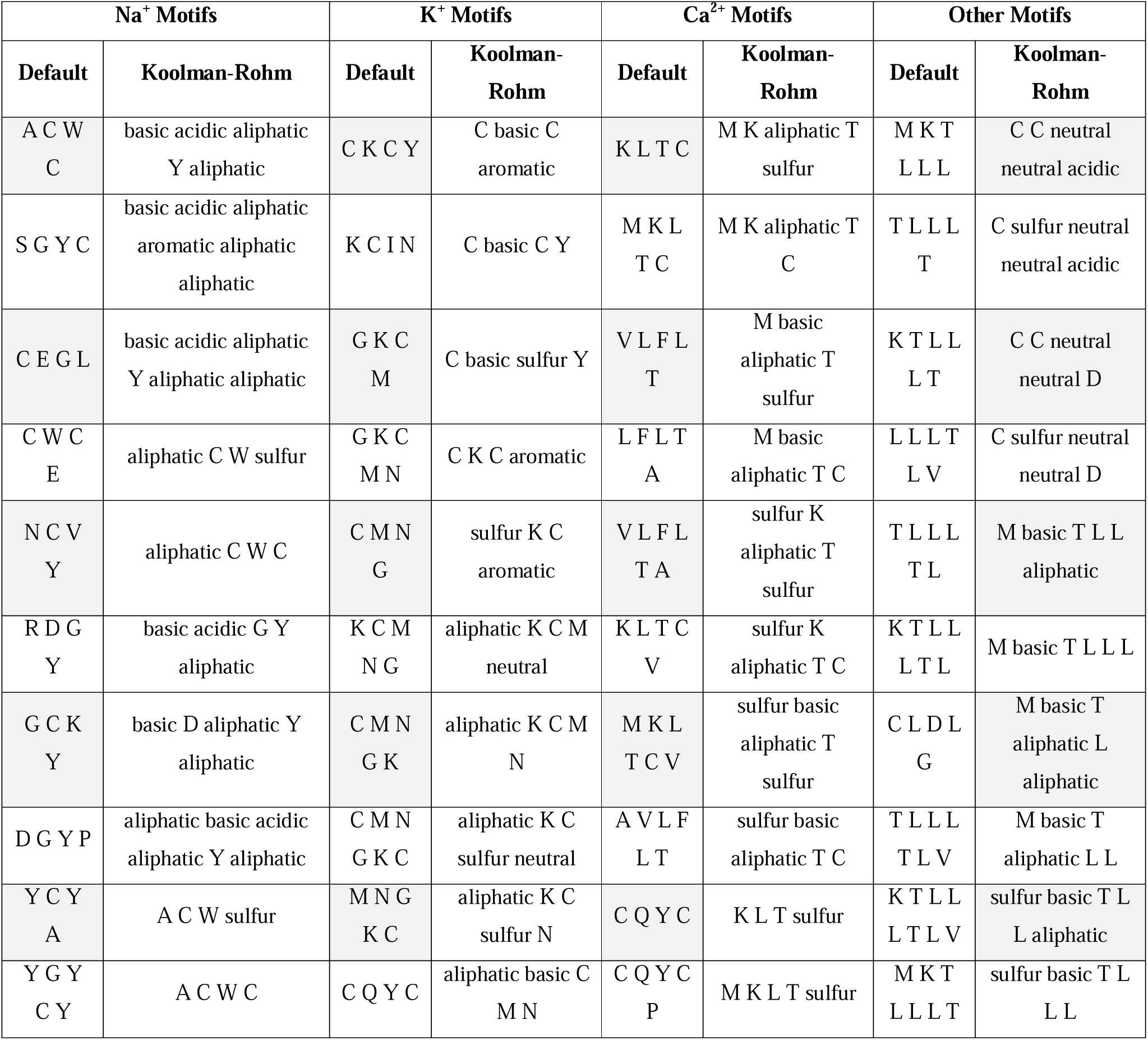
Top motifs identified by MERCI using Default and Koolman-Rohm configurations across four independent datasets (Na. D**, K**D**, Ca²**D**, and Other).**

Despite testing various motifs collected using a range of configurations on the training dataset, the majority of predictions successfully identified negatives but failed to yield hits in the independent dataset, as reflected by the low number of positive hits. This underscores that MERCI motifs may contribute only marginally to identifying true modulators in unseen data. The complete confusion matrices and configuration-specific performance metrics are available in Supplementary Table S4.

### 3.3 Alignment-free methods

This describes machine learning and protein language model (PLM) approaches that predict ion channel modulators without relying on sequence alignment, emphasizing feature-based modeling and contextual sequence representations.

#### 3.3.1 Machine Learning-based Classifiers

We evaluated multiple machine learning models (e.g., ET, RF, SVM) on four ion channel-modulating protein datasets (Na□, K□, Ca²□, and Other) using various compositional features, including AAC, DPC, their combination (AAC+DPC), and a high-dimensional ALLCOMP feature set comprising 9,190 features. Table 6 summarizes the performance of the best-performing models on different feature combinations. Overall, models trained on the ALLCOMP feature set consistently demonstrated superior performance compared to those utilizing simpler descriptors such as AAC or DPC. For example, on the Na□ independent dataset, the ALLCOMP-based model achieved an AUC of 0.962 and an MCC of 0.591, outperforming both AAC (AUC: 0.949, MCC: 0.532) and DPC (AUC: 0.952, MCC: 0.535).

**Table 6:** Performance evaluation of tree based models on the different data using compositional features.

| Dataset | Features | Classifier | Coss-validation Dataset |  |  |  |  |  | Independent Dataset |  |  |  |  |  |
| --- | --- | --- | --- | --- | --- | --- | --- | --- | --- | --- | --- | --- | --- | --- |
|  |  |  | SENS | SPEC | PPV | ACC | MCC | AUC | SENS | SPEC | PPV | ACC | MCC | AUC |
| Na+ | AAC | ET | 0.640 | 0.970 | 0.870 | 0.880 | 0.650 | 0.960 | 0.547 | 0.978 | 0.886 | 0.876 | 0.632 | 0.949 |
|  | DPC | RF | 0.460 | 0.985 | 0.910 | 0.860 | 0.580 | 0.940 | 0.375 | 0.993 | 0.941 | 0.846 | 0.535 | 0.952 |
|  | AAC+DPC | RF | 0.660 | 0.975 | 0.880 | 0.870 | 0.600 | 0.940 | 0.577 | 0.978 | 0.871 | 0.859 | 0.576 | 0.955 |
|  | ALLCOMP | ET | 0.723 | 0.994 | 0.974 | 0.933 | 0.743 | 0.944 | 0.453 | 0.99 | 0.935 | 0.863 | 0.591 | 0.962 |
| K+ | AAC | ET | 0.530 | 0.960 | 0.770 | 0.850 | 0.500 | 0.910 | 0.431 | 0.966 | 0.770 | 0.854 | 0.502 | 0.920 |
|  | DPC | ET | 0.400 | 0.990 | 0.880 | 0.850 | 0.520 | 0.930 | 0.294 | 0.983 | 0.821 | 0.838 | 0.427 | 0.911 |
|  | AAC+DPC | ET | 0.500 | 0.985 | 0.910 | 0.860 | 0.580 | 0.940 | 0.303 | 0.985 | 0.846 | 0.842 | 0.445 | 0.917 |
|  | ALLCOMP | ET | 0.660 | 0.975 | 0.880 | 0.890 | 0.700 | 0.935 | 0.486 | 0.981 | 0.869 | 0.877 | 0.590 | 0.940 |
| Ca+ | AAC | ET | 0.600 | 0.975 | 0.820 | 0.910 | 0.640 | 0.950 | 0.560 | 0.971 | 0.778 | 0.907 | 0.610 | 0.939 |
|  | DPC | ET | 0.420 | 0.980 | 0.860 | 0.880 | 0.540 | 0.920 | 0.267 | 0.990 | 0.833 | 0.878 | 0.428 | 0.919 |
|  | AAC+DPC | ET | 0.580 | 0.970 | 0.850 | 0.900 | 0.660 | 0.940 | 0.480 | 0.990 | 0.840 | 0.880 | 0.442 | 0.936 |
|  | ALLCOMP | RF | 0.590 | 0.985 | 0.910 | 0.900 | 0.680 | 0.940 | 0.373 | 0.988 | 0.848 | 0.893 | 0.519 | 0.943 |
| Other | AAC | ET | 0.560 | 0.970 | 0.810 | 0.880 | 0.580 | 0.930 | 0.505 | 0.971 | 0.797 | 0.871 | 0.544 | 0.933 |
|  | DPC | ET | 0.400 | 0.985 | 0.870 | 0.880 | 0.490 | 0.930 | 0.377 | 0.990 | 0.889 | 0.857 | 0.478 | 0.926 |
|  | AAC+DPC | ET | 0.59 | 0.975 | 0.91 | 0.91 | 0.72 | 0.945 | 0.476 | 0.978 | 0.809 | 0.859 | 0.488 | 0.937 |
|  | ALLCOMP | ET | 0.630 | 0.975 | 0.910 | 0.900 | 0.720 | 0.945 | 0.545 | 0.978 | 0.859 | 0.893 | 0.629 | 0.946 |

(Abbreviations: SENS: Sensitivity; SPEC: Specificity; PPV: Positive Predictive Value; ACC: Accuracy; MCC: Matthews Correlation Coefficient; AUC: Area Under the Receiver Operating Characteristic Curve)

Similarly, for the K□ dataset, ALLCOMP yielded the highest performance with an AUC of 0.940 and MCC of 0.590. In the case of the Ca²□ dataset, the ALLCOMP model again outperformed others with an AUC of 0.943 and MCC of 0.519. Finally, for the Other ions dataset, the ALLCOMP feature set produced the most robust results, achieving an AUC of 0.946 and MCC of 0.629. These results underscore the advantage of comprehensive feature representations in enhancing model generalizability and predictive power across diverse ion-channel modulator datasets.

However, the performance gain with ALLCOMP came at a cost. Despite a massive increase in feature dimensionality (from 20 in AAC and 400 in DPC to 9,190 in ALLCOMP), the improvement in predictive performance was only modest in many cases. For example, in the Ca²□ dataset, AAC achieved an AUC of 0.949, while ALLCOMP reached 0.962 a small gain considering the 450-fold increase in features. Moreover, sensitivity remained relatively low in some datasets, particularly for calcium and potassium, indicating a challenge in detecting true positives in these cases. In summary, while ALLCOMP-based models yield the best overall performance, the trade-off between computational complexity and performance gain should be considered carefully. Simpler models using AAC or DPC still provide competitive results and may be more practical for large-scale screening or interpretable modeling. Supplementary Table S5 provides detailed performance metrics of all the ML models on different datasets.

#### 3.3.2 Feature Importance Analysis

We employed the ET classifier, which showed superior performance across most datasets, to evaluate the discriminative power of the 9,169 features generated by the Pfeature tool. Based on this analysis, the top 10 most informative features are presented in Table 7. This convergence strongly indicates that these features capture fundamental sequence characteristics underlying ion channel modulation, independent of channel type.

**Table 7.** Top 10 discriminative features selected by ExtraTrees across all ion channel datasets, showing mean values in positive and negative datasets, fold differences, and importance scores.

| Top Features | Negative Data Mean | Na+ |  | K+ |  | Ca+ |  | Other |  |
| --- | --- | --- | --- | --- | --- | --- | --- | --- | --- |
|  |  | Positive Data Mean | Importance | Positive Data Mean | Importance | Positive Data Mean | Importance | Positive Data Mean | Importance |
| SER_C | 0.114 | 0.259 | 0.021 | 0.259 | 0.021 | 0.253 | 0.021 | 0.257 | 0.021 |
| QSO1_G_C | 1.90E-05 | 6.80E-05 | 0.0157 | 6.70E-05 | 0.0157 | 6.70E-05 | 0.0157 | 7.20E-05 | 0.0157 |
| PAAC1_C | 2.749 | 11.875 | 0.015 | 11.875 | 0.015 | 11.458 | 0.015 | 11.958 | 0.015 |
| AAC_C | 2.751 | 11.875 | 0.014 | 11.875 | 0.014 | 11.458 | 0.014 | 11.958 | 0.014 |
| SOC1_G1 | 9975.083 | 13931.258 | 0.011 | 13867.774 | 0.011 | 13133.194 | 0.011 | 12642.942 | 0.011 |
| ATC_S | 0.332 | 0.774 | 0.011 | 0.804 | 0.011 | 0.781 | 0.011 | 0.81 | 0.011 |
| PCP_SC | 0.062 | 0.111 | 0.009 | 0.114 | 0.009 | 0.112 | 0.009 | 0.114 | 0.009 |
| CTC_137 | 0.026 | 0.222 | 0.009 | 0.123 | 0.009 | 0.101 | 0.009 | 0.105 | 0.009 |
| DPC1_CC | 0.167 | 1.289 | 0.009 | 0.800 | 0.009 | 1.267 | 0.009 | 1.910 | 0.009 |
| DPC1_YC | 0.098 | 1.131 | 0.007 | 0.655 | 0.007 | 0.454 | 0.007 | 0.407 | 0.007 |

Among the top-ranked features, cysteine-related features (SER_C, QSO1_G_C, AAC_C, PAAC1_C) appeared repeatedly with high importance scores. Their enrichment in positive datasets aligns with the well-established role of cysteine residues in forming disulfide bridges that stabilize the compact structural motifs (e.g., inhibitor cystine knot) commonly found in ion channel toxins. This provides direct biological validation of the model’s feature selection.

Other high-ranking features included sequence-order descriptors (SOC1_G1, QSO1_G_C, QSO1_SC_C) and DPC features (DPC1_CC, DPC1_YC, DPC1_CK). These highlight that not only overall AAC but also residue ordering and local sequence context contribute critically to differentiating modulators from non-modulators. Interestingly, physicochemical property-based descriptors (PCP_SC, PCP_Z5, PCP_Z3) also featured prominently, suggesting that differences in charge, polarity, and hydrophobicity may help explain why certain peptides selectively interact with ion channels.

The robustness of these features across multiple datasets highlights their generalizability and suggests they may serve as core sequence signatures of ion channel modulation. Their consistent selection also enhances confidence in the stability of the ET model, minimizing concerns of overfitting to any particular dataset.

#### 3.3.3 PLM-Based Classifiers

We next evaluated the predictive capabilities of different fine-tuned PLMs (ESM2, ProtBERT, Distilbert etc.) across all four ion channel-modulating protein datasets. These models, which utilize deep contextual embeddings of protein sequences, demonstrated superior performance over traditional machine learning (ML) classifiers built on compositional features. Among the tested models, ESM2-t33 consistently delivered the best results (Table 8). On the Na□ dataset, ESM2-t33 achieved an outstanding AUC of 0.988 and MCC of 0.826 on the cross-validation dataset. It maintained high performance on the independent dataset as well (AUC: 0.982 and MCC: 0.799, AUC: 0.996). Similarly, on the K□ dataset, it attained AUC: 0.973, MCC: 0.838, and continued to perform strongly on the independent set (AUC: 0.971 and MCC: 0.812). On the Ca²□ and Other datasets as well, ESM2-t33 achieved top AUC (0.955 and 0.957, respectively) with MCC values (0.768 and 0.854, respectively) on independent set evaluations. Detailed performance metrics of the rest of the PLMs on different datasets are provided in Supplementary Table S6.

**Table 8:** Performance evaluation of PLM-based classifiers on the different datasets.

| Dataset | PLM Classifier | Coss-validation Dataset |  |  |  |  |  | Independent Dataset |  |  |  |  |  |
| --- | --- | --- | --- | --- | --- | --- | --- | --- | --- | --- | --- | --- | --- |
|  |  | SENS | SPEC | PPV | ACC | MCC | AUC | SENS | SPEC | PPV | ACC | MCC | AUC |
| Na+ | ESM2-t33 | **0.803** | **0.994** | **0.980** | **0.960** | **0.826** | **0.988** | **0.788** | **0.983** | **0.933** | **0.930** | **0.799** | **0.982** |
|  | bert-base-uncased | 0.792 | 0.966 | 0.906 | 0.799 | 0.809 | 0.941 | 0.789 | 0.917 | 0.887 | 0.694 | 0.768 | 0.934 |
|  | ProtBERT | 0.981 | 0.970 | 0.909 | 0.972 | 0.926 | 0.963 | 0.875 | 0.956 | 0.862 | 0.937 | 0.827 | 0.955 |
| K+ | ESM2-t33 | **0.852** | **0.985** | **0.932** | **0.952** | **0.838** | **0.973** | **0.829** | **0.976** | **0.896** | **0.947** | **0.812** | **0.966** |
|  | bert-base-uncased | 0.494 | 0.998 | 0.876 | 0.526 | 0.539 | 0.915 | 0.449 | 0.983 | 0.859 | 0.489 | 0.494 | 0.904 |
|  | ProtBERT | 0.873 | 0.931 | 0.727 | 0.921 | 0.747 | 0.942 | 0.840 | 0.929 | 0.685 | 0.916 | 0.709 | 0.925 |
| Ca+ | ESM2-t33 | **0.753** | **0.976** | **0.859** | **0.942** | **0.771** | **0.950** | **0.733** | **0.981** | **0.873** | **0.942** | **0.768** | **0.963** |
|  | bert-base-uncased | 0.690 | 0.993 | 0.918 | 0.606 | 0.663 | 0.912 | 0.653 | 0.937 | 0.893 | 0.590 | 0.653 | 0.900 |
|  | ProtBERT | 0.873 | 0.931 | 0.727 | 0.921 | 0.747 | 0.942 | 0.840 | 0.929 | 0.685 | 0.916 | 0.709 | 0.925 |
| Other | ESM2-t33 | **0.785** | **0.993** | **0.930** | **0.950** | **0.836** | **0.975** | **0.831** | **0.985** | **0.949** | **0.955** | **0.854** | **0.957** |
|  | bert-base-uncased | 0.631 | 0.995 | 0.817 | 0.919 | 0.682 | 0.926 | 0.624 | 0.964 | 0.808 | 0.897 | 0.650 | 0.918 |
|  | ProtBERT | 0.743 | 0.997 | 0.938 | 0.941 | 0.798 | 0.827 | 0.733 | 0.985 | 0.925 | 0.936 | 0.787 | 0.816 |

(Abbreviations: SENS: Sensitivity; SPEC: Specificity; PPV: Positive Predictive Value; ACC: Accuracy; MCC: Matthews Correlation Coefficient; AUC: Area Under the Receiver Operating Characteristic Curve; Note: The bold values indicate the best-performing models for each dataset)

These results substantially outperform those achieved by ML models using composition-based features like AAC (20), DPC (400), or even ALLCOMP (9190 features). While the ML models showed reasonable performance, especially with ALLCOMP, the performance gains from PLMs were significantly higher despite not requiring manual feature engineering. This highlights the ability of PLMs to capture intricate contextual and evolutionary information from sequence data directly. Overall, ESM2-t33 emerged as the most effective model, consistently achieving the highest performance across all datasets. Other models, including ESM2 and ProtBERT, also performed comparably well, further supporting the effectiveness of PLM-based approaches for protein classification tasks

### 3.4 Performance of Hybrid Models

ML-based models have yielded encouraging results, they often exhibit high AUC values while displaying suboptimal performance in terms of the MCC. In contrast, PLMs, particularly ESM2-t33, have demonstrated consistently superior performance, achieving high sensitivity, specificity, and AUC across all evaluated datasets. Our analysis indicates an average AUC of 0.97 and an MCC of 0.83 using ESM2-t33. To further improve the performance, we developed a hybrid model that integrates multiple strategies to enhance the precision of ion channel modulator prediction.

In this hybrid framework, we combined a composition-based ML model with BLAST- and MERCI-based motif search approaches. Specifically, each protein was first classified using BLAST at an E-value cutoff of 10^-3^. Proteins not identified by BLAST were then subjected to MERCI motif-based classification. Finally, for sequences not identified by either method (i.e., no hits), a PLM-based model was employed to predict their modulatory potential. Interestingly, combining MERCI with PLMs did not result in any performance improvement. However, the integration of BLAST with PLMs led to a noticeable enhancement in overall performance, indicating better balance and agreement between true and predicted classifications. As summarized in Table 9, the hybrid method demonstrated improved performance when all BLAST at an E-value cutoff of 10^-3^ were combined with the PLM-based predictions. The best-performing model was the ESM2-t33-based hybrid, which achieved average values of 0.93 for sensitivity, 0.98 for specificity, 0.95 for precision, 0.97 for accuracy, 0.91 for MCC, and 0.98 for AUC on the independent datasets.

**Table 9:** Performance of the hybrid method that combines the ESM2-t33 model with BLAST search on an independent dataset.

| Dataset | Classifier | Threshold | SENS | SPEC | PPV | ACC | MCC | AUC |
| --- | --- | --- | --- | --- | --- | --- | --- | --- |
| Na<sup>+</sup> | ESM2-t33 | 0.21 | 0.859 | 0.959 | 0.866 | 0.935 | 0.820 | 0.982 |
|  | ESM2-t33 + BLAST | 0.41 | 0.992 | 0.971 | 0.994 | 0.988 | 0.958 | 0.997 |
| K<sup>+</sup> | ESM2-t33 | 0.21 | 0.872 | 0.959 | 0.848 | 0.940 | 0.822 | 0.966 |
|  | ESM2-t33 + BLAST | 0.21 | 0.988 | 0.967 | 0.984 | 0.981 | 0.957 | 0.977 |
| Ca<sup>2+</sup> | ESM2-t33 | 0.35 | 0.760 | 0.980 | 0.877 | 0.946 | 0.786 | 0.963 |
|  | ESM2-t33 + BLAST | 0.35 | 0.861 | 0.985 | 0.935 | 0.961 | 0.874 | 0.982 |
| Other | ESM2-t33 | 0.30 | 0.861 | 0.985 | 0.935 | 0.961 | 0.874 | 0.957 |
|  | ESM2-t33 + BLAST | 0.30 | 0.861 | 0.977 | 0.879 | 0.958 | 0.845 | 0.970 |

(Abbreviations: SENS: Sensitivity; SPEC: Specificity; PPV: Positive Predictive Value; ACC: Accuracy; MCC: Matthews Correlation Coefficient; AUC: Area Under the Receiver Operating Characteristic Curve)

### 3.5 Cross-dataset prediction

To evaluate the robustness of the best-performing models, we carried out cross-dataset prediction experiments in which models trained on one ion channel class (Na□, K□, Ca²□, or Other) were tested on the remaining classes. This analysis provides an assessment of model transferability beyond within-dataset evaluation. The results (Table 10) show that models trained on individual channel datasets achieved consistently high AUC values when applied to other channel classes, ranging from 0.937 to 0.981. Models trained on Na□ and K□ datasets yielded the strongest cross-prediction, with AUC values above 0.96 across most test sets. Ca²□ models performed comparably, while the “Other” class showed slightly lower transferability but still maintained AUC values close to 0.95. These findings indicate that classifiers capture both channel-specific and shared molecular signals, allowing generalization across related ion channel modulators.

**Table 10.**
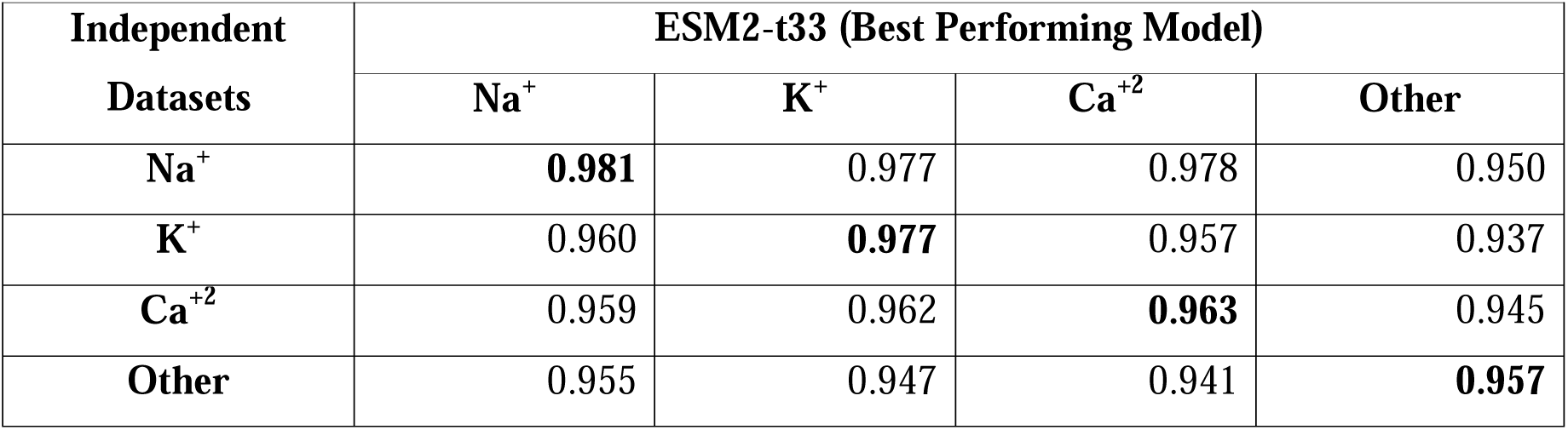
Cross-dataset prediction performance of the best-performing ESM2-t33 model evaluated using AUC.

| Independent Datasets | ESM2-t33 (Best Performing Model) |  |  |  |
| --- | --- | --- | --- | --- |
|  | Na<sup>+</sup> | K<sup>+</sup> | Ca<sup>2+</sup> | Other |
| Na<sup>+</sup> | **0.981** | 0.977 | 0.978 | 0.950 |
| K<sup>+</sup> | 0.960 | **0.977** | 0.957 | 0.937 |
| Ca<sup>2+</sup> | 0.959 | 0.962 | **0.963** | 0.945 |
| Other | 0.955 | 0.947 | 0.941 | **0.957** |

### 3.6 Unified Model Training and Evaluation

To further assess the generalizability of our approach, we trained the best-performing model (ESM2-t33) on the combined training datasets from all ion channel classes (Na□, K□, Ca²□, and Other) and evaluated its performance separately on the independent test sets of each channel class. This unified training strategy simulates a real-world application where prior knowledge from multiple ion channels can be leveraged to improve predictive accuracy. The unified model achieved high performance across all test sets, with AUC values ranging from 0.968 (Other) to 0.980 (Na□) and MCC values between 0.815 (Ca²□) and 0.833 (Other) (Table 11). Notably, the model retained strong predictive power for each individual channel type, confirming that shared sequence and compositional features can be exploited without compromising class-specific accuracy.

**Table 11:**
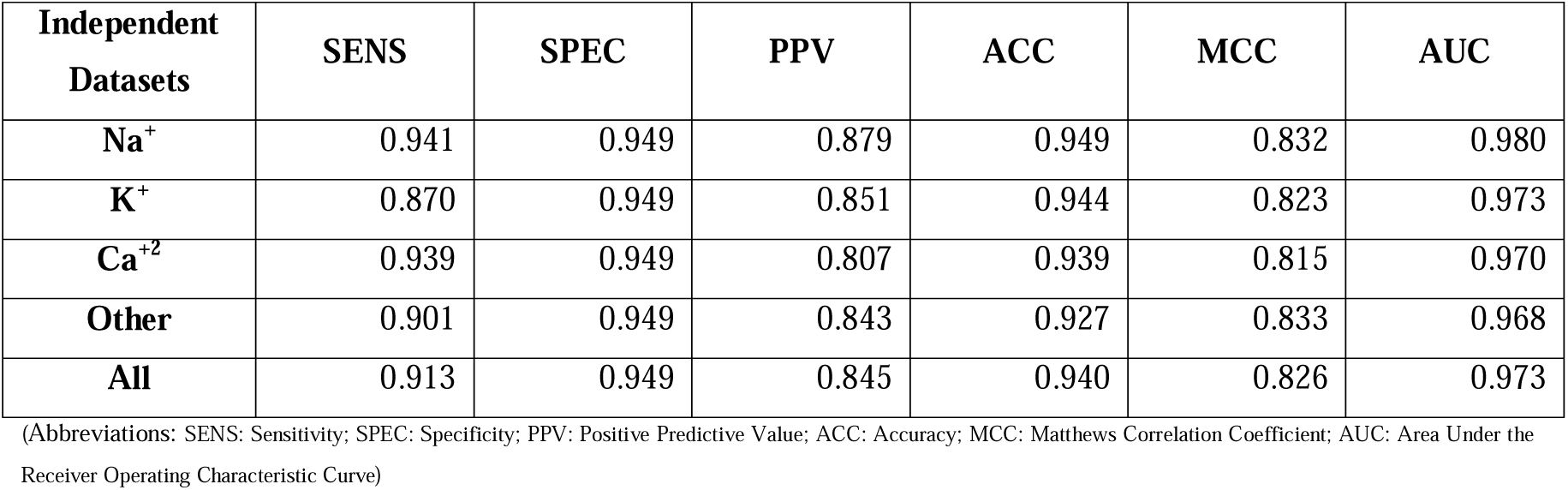
Performance of the unified ESM2-t33 model trained on combined datasets of all ion channels (Na. D**, K**D**, Ca²**D**, and Other) and evaluated on independent test sets.**

### 3.7 Benchmarking with Existing Methods

Several computational tools have been developed to predict ion channel modulating proteins, each with distinct dataset choices and scope. However, these approaches vary widely in their coverage, data curation, and applicability. For instance, MetaNaBP (2024) and PEP-PREDNa□ (2022) tools represent valuable contributions to the field, a key limitation lies in their reliance on datasets originally extracted from NTxPred, developed by Saha and Raghava (2007). Although NTxPred was an important pioneering effort, its training data are now over a decade old and do not reflect the substantial growth and refinement of protein sequence databases in recent years. By contrast, some recent tools have constructed newer datasets. For example, NallPred (2024) and PrIMP (2022) curated fresh collections of ion channel modulators, although their dataset sizes remain limited compared to present-day sequence repositories. Notably, the STACKION predictor directly reused PrIMP’s dataset, thereby inheriting the same data scope and constraints. Other methods have focused on specific families or channel types rather than providing broad coverage. For example, RBF Network (2013), iCNTxType (2014), and AVC-SVM (2017) targeted conotoxins only, while PPLK□C was designed exclusively for K□ channel toxins. These specialized predictors, though valuable for their respective domains, lack the generalizability needed for comprehensive ion channel modulator prediction. In contrast, IonNTxPred introduces the largest and most up-to-date dataset of ion channel modulators, systematically curated from contemporary protein repositories and filtered using CD-HIT at 40% sequence identity to remove redundancy and prevent data leakage. This not only ensures greater diversity and modern annotation accuracy but also improves the robustness and generalizability of the predictive models. Notably, most earlier predictors relied on 80-90% redundancy thresholds only. By benchmarking against preexisting tools, IonNTxPred establishes itself as a more comprehensive and reliable framework for predicting modulators of Na□, K□, Ca²□, and other ion channels.

Our benchmarking was performed using the IonNTxPred independent datasets across different ion channels (Na□, K□, Ca²□, and Others), enabling a fair and comprehensive evaluation. Unfortunately, several tools could not be compared due to practical limitations: STACKION did not provide access to their tool; MetaNaBP’s GitHub repository lacked a deployable model; PPLK□C only supports single-sequence prediction, which is infeasible for large-scale benchmarking; and the IonChannPred2.0 web server was not operational during our evaluation. Among the evaluated tools, our hybrid framework integrating ESM2-t33 with BLAST consistently demonstrated superior performance across key evaluation metrics. Specifically, IonNTxPred achieved an AUC of 0.997 and an MCC of 0.958 for Na□ modulators, significantly outperforming existing methods such as iCTX (AUC: 0.626, MCC: 0.107), NTXpred (AUC: 0.781, MCC: 0.431), and PEP-PREDNa□ (AUC: 0.696, MCC: 0.444). Similarly, IonNTxPred maintained high predictive performance for K□ and Ca²□ modulators, with AUC values exceeding 0.997 and 0.982, respectively (Table 12). Although the PrIMP model proposed by Lee et al. demonstrated competitive performance with AUCs of 0.965, 0.919, and 0.948 for Na□, K□, and Ca²□ modulators, respectively, it did not surpass the performance of IonNTxPred. This comparative analysis highlights the robustness and enhanced predictive capability of our proposed method for identifying ion channel modulators across diverse ion channel types.

**Table 12:**
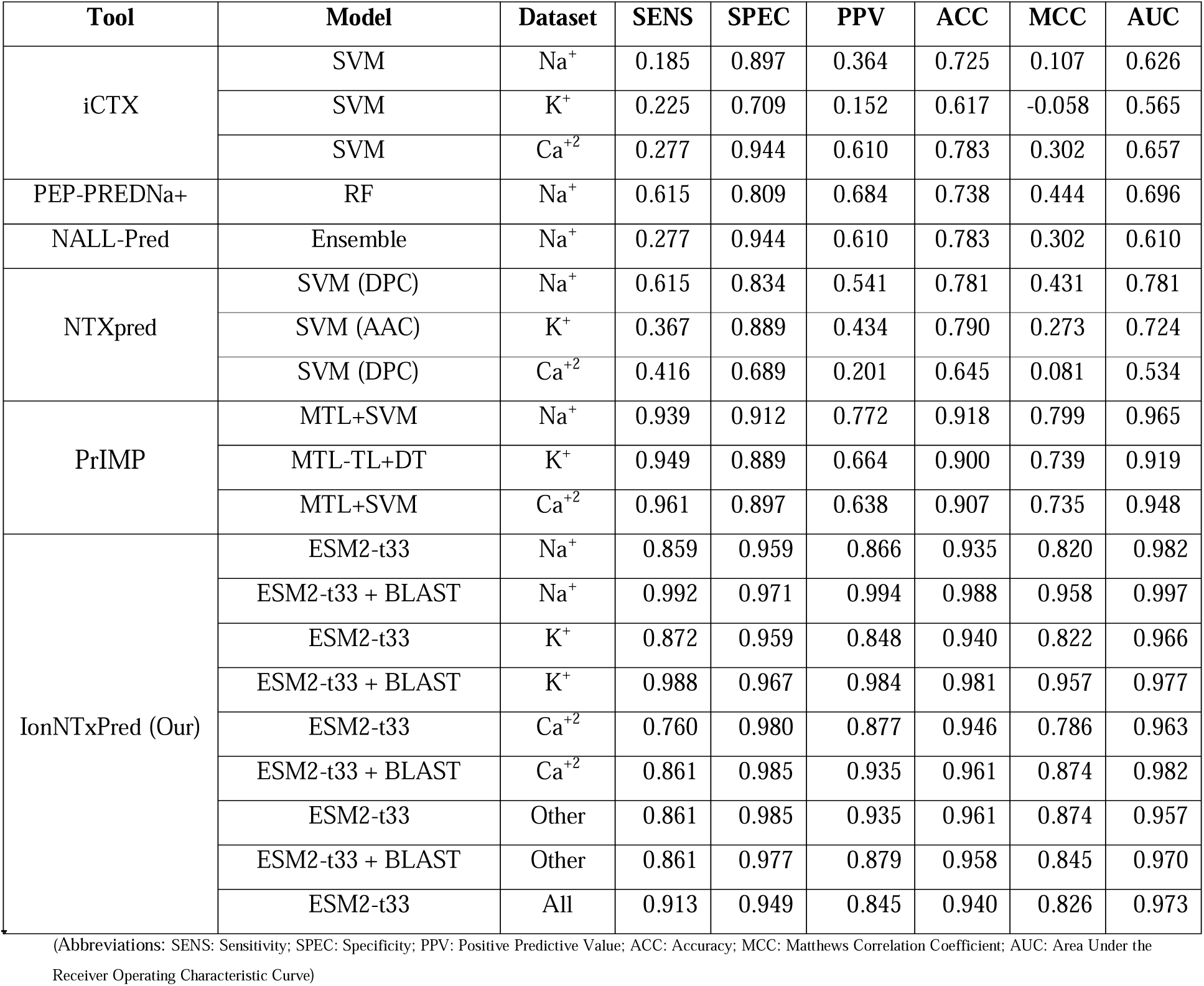
Performance of existing methods on independent datasets of IonNTxPred.

### 3.8 Case Study: Genome-Wide Prediction of Ion Channel Toxins

To demonstrate the utility of IonNTxPred beyond benchmark datasets, we applied it to proteomes of representative venomous and toxic organisms with sequenced genomes from NCBI (O’Leary et al., 2024), including cone snails (*Conus episcopatus*, *Conus munga*), snakes (*Naja naja*, *Ophiophagus hannah*), the horseshoe crab (*Limulus polyphemus*), and the pufferfish (*Takifugu rubripes*). The analysis revealed substantial numbers of ion channel-modulating proteins across all groups, with notable instances of moonlighting activity, where the same protein exhibited modulatory potential against multiple ion channel types. This highlights the rich diversity of naturally encoded toxins and their potential for translational applications in drug discovery and safety assessment (Figure 6A). These species are renowned sources of ion channel modulators, many of which have inspired or directly yielded therapeutic agents. For example, snake toxin cobratoxin b from Naja naja has been studied as a neuromuscular blocker [9498573, 35418867]. By systematically screening entire proteomes with IonNTxPred, we highlight the untapped reservoir of ion channel-modulating proteins that remain uncharacterized in these organisms. This capability is not only valuable for drug discovery, where novel toxins can serve as scaffolds for selective channel modulators, but also for biosafety and toxicology, ensuring early identification of harmful proteins in food sources (e.g., pufferfish) or environmental encounters (e.g., snake and snail venoms). Such genome-scale prediction exemplifies how computational pipelines can bridge natural toxin diversity with rational drug design, accelerating the translation of venom-derived peptides into safe and effective therapeutics.

**Figure 6.**
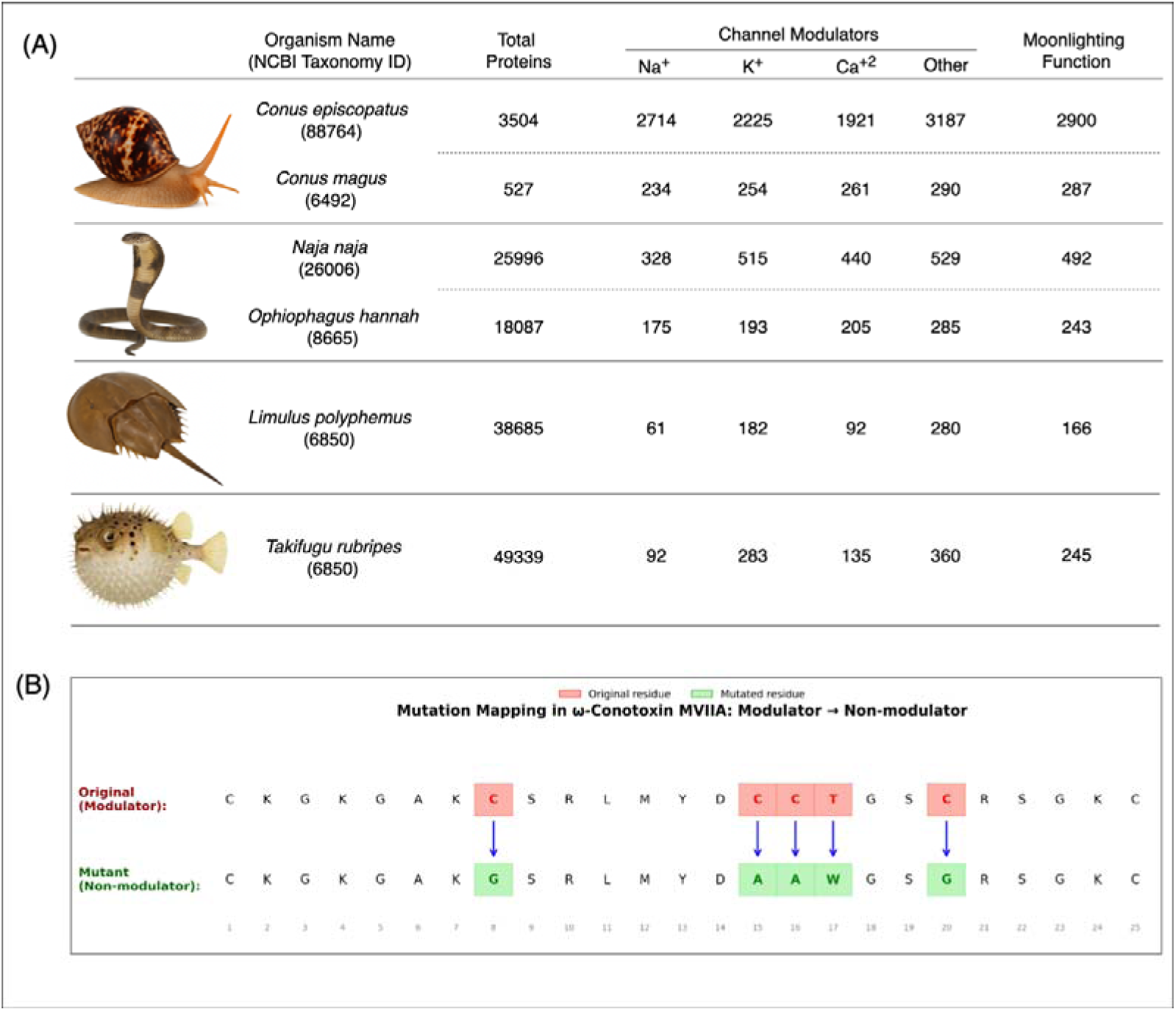
(A) Case study results showing predicted ion channel modulators (Na□, K□, Ca^2^□, and others) and moonlighting toxins identified from proteomes of representative venomous and toxic species. (B) Mutation study using the IonNTxPred design module showing conversion of ω-conotoxin MVIIA from Modulator to Non-modulator.

To further demonstrate the design utility of IonNTxPred, we generated a mutant peptide CKGKGAKGSRLMYDAAWGSGRSGKC through five site-specific substitutions in the canonical ω-conotoxin MVIIA scaffold, using the IonNTxPred computational platform. The parent peptide, MVIIA (ziconotide), is a disulfide-rich marine snail toxin and a clinically approved analgesic that acts as a potent N-type (CaV2.2) calcium channel blocker (McGivern, 2007). Mutations introduced by IonNTxPred disrupted four of the six cysteine residues that form the characteristic three-disulfide bond framework, along with a polar-to-aromatic substitution (Thr→Trp), leading to significant loss of structural integrity and channel-binding determinants (Figure 6B). As a result, the designed variant was predicted to function as a non-modulator, highlighting the sensitivity of ion channel modulation to cysteine patterning and side-chain orientation. This example underscores the utility of IonNTxPred in desining mutational landscapes, enabling researchers to evaluate how specific amino acid changes affect the modulatory activity of peptides, and to guide rational engineering of safer, non-toxic, or functionally selective peptide analogues.

### 3.9 Applications

IonNTxPred enables powerful applications across drug discovery, safety screening, and biosafety assessment. It can be used to rapidly identify ion channel-specific neurotoxins in venom proteomes or biological databases, accelerating the annotation of novel peptides. In drug design, IonNTxPred offers a means to redesign “undruggable” molecules by reducing their off-target ion channel interactions, akin to how the funnel-web peptide *Hi1a* is being explored as a cardioprotective agent in clinical trials for heart attack and stroke treatment (Duggan et al., 2021). Additionally, conotoxin-derived drugs such as *ziconotide (Prialt)* consists of 25 amino acid residues, an approved N-type calcium channel blocker by the FDA in 20004, and *Contulakin-G,* which entered Phase Ib clinical trials for chronic pain, exemplify the therapeutic potential of venom peptides (Gawade, 2007; Safavi-Hemami, Brogan, & Olivera, 2019), and IonNTxPred could streamline identification of similar leads. For safer peptide therapeutics, it aids in designing molecules with minimal ion channel toxicity, such as *rotigaptide (ZP123)*, a sodium channel modulator under investigation for atrial fibrillation (Phase II) (Guerra, Everett, Lee, Wilson, & Olgin, 2006).

Beyond therapeutic discovery, IonNTxPred is instrumental in the assessment of biosafety, particularly for genetically modified proteins intended for agricultural use, such as pesticidal proteins, by evaluating their potential unintended activity on mammalian ion channels, thereby supporting the safety of non-target organisms. The field of drug repurposing further benefits from IonNTxPred, where existing drugs are evaluated for new therapeutic roles based on their predicted ion channel modulation profiles. This cost-efficient strategy leverages pre-existing pharmacological and safety data, revealing novel electrophysiological effects and facilitating safer clinical implementation.

To demonstrate the translational reach of IonNTxPred, a systematic screening of approved and clinically studied therapeutic peptides from the ThPDB2 database (Jain, Gupta, Patiyal, & Raghava, 2024)[38830503] was conducted for ion channel–modulating activity. Several clinically utilized peptides emerged as strong predictors for Na□, K□, or Ca²□ channel modulation, illustrating new opportunities for drug repurposing and highlighting possible off-target electrophysiological risks that warrant consideration. For instance, CT-011, a 64-amino-acid immune checkpoint inhibitor developed for cancer immunotherapy through PD-1/PD-L1 axis targeting, was predicted to act as a multi-channel modulator affecting Na□, Ca²□, and other channels (See Table 13). This prediction underscores IonNTxPred’s capacity to flag mechanistic pathways that could influence cardiac or neurological safety profiles, informing subsequent electrophysiological validation and the rational design of next-generation peptide therapeutics with immunomodulatory or neuroactive functions. It is important to emphasize that these predictions are computational and require ultimate validation through rigorous wet lab experimentation, including electrophysiological assays and functional studies, to confirm the predicted ion channel modulatory effects and ensure translational relevance.

**Table 13:** Predicted ion channel-modulating activity of clinically approved therapeutic peptides/proteins from ThPDB2.

| Therapeutic ID | Therapeutic Name | Disease/Indication | Modulator | Moonlighting Activity |
| --- | --- | --- | --- | --- |
| Th1329 | CT-011 | Cancer | Na+, Ca+ , Other | Yes |
| Th1272 | Tigapotide | Hormonal Disorders | Na+, Ca+ , Other | Yes |
| Th1027 | Insulin Regular | Adjunct Therapy | Na+, K+, Ca+ , Other | Yes |
| Th1007 | Leuprolide | Cancer | Na+, K+, Ca+ , Other | Yes |
| Th1566 | Coagulation factor VII human | Cardiac Disorders | K+ | No |
| Th1398 | Thrombin alfa | Adjunct Therapy | K+ | No |

### 3.10 Implementation and Public Accessibility

To make IonNTxPred widely accessible, we have developed a user-friendly web server (https://webs.iiitd.edu.in/raghava/ionntxpred/) that integrates multiple modules designed to address different aspects of ion channel toxin prediction and analysis.

**1. Prediction Module** This core module allows users to submit protein sequences in FASTA format for classification as ion channel modulators or non-modulators. Users can select between ESM2-t33 protein language model and a hybrid approach that combines PLM predictions with BLAST outputs for improved performance. This provides flexibility in balancing speed, accuracy, and interpretability.
**2. Design Module** The design module supports protein engineering by mutation. Users can generate variants of their sequences, such as point mutants or exhaustive mutational scans and then predict their ion channel toxicity. This is particularly useful for optimizing therapeutic peptides by reducing toxicity while retaining efficacy.
**3. Channel Toxicity Scanning/Mapping Module** This module goes beyond binary classification by scanning a query protein to identify specific regions that contribute to ion channel modulation. By localizing toxic segments, the tool helps researchers understand which residues or motifs may be responsible for activity, guiding targeted mutagenesis and rational protein design.
**4. BLAST Search Module** The BLAST-based search option allows users to query their sequences against a curated database of known ion channel modulators and non-modulators. This homology-driven approach provides rapid annotation support, highlighting evolutionary conservation and enabling predictions when sequence similarity is available.
**5. Motif Scan Module** Using the MERCI motif extraction tool, this module enables the identification and mapping of biologically relevant motifs within protein sequences. Motif-level insights are critical for understanding conserved structural signatures of ion channel modulators and for designing novel peptides with tailored activity.

The web interface has been implemented with a responsive HTML and PHP design, ensuring compatibility across devices, platforms, and operating systems. For large-scale or offline use, we also provide a standalone version of IonNTxPred, which can be downloaded from the website. In addition, the source code is openly available on GitHub (https://github.com/raghavagps/ionntxpred/), and our pretrained PLM models are hosted on Hugging Face (https://huggingface.co/raghavagps-group/IonNTxPred). These models are not only ready-to-use for ion channel modulator prediction but can also be fine-tuned for other related tasks, such as predicting carcinogenicity. identifying therapeutic peptide scaffolds, or screening engineered proteins in synthetic biology. This modular and extensible design ensures that IonNTxPred can evolve with new datasets and applications, serving as a versatile resource for both computational biologists and experimental researchers.

## 4 Discussion and Conclusion

The development of therapeutic proteins and peptides has become a cornerstone of modern medicine, offering high specificity and efficacy for a wide range of diseases. The clinical success of these biomolecules is underscored by the growing number of approved drugs; by 2022, 109 peptide-based drugs had received FDA approval (K. Sharma, Sharma, Sharma, & Jain, 2023), with an additional 8 approved since then (Al Musaimi, AlShaer, de la Torre, & Albericio, 2025; Al Shaer, Al Musaimi, Albericio, & de la Torre, 2024). To accelerate the discovery and design of these therapeutics, various computational tools have been developed over the years. However, a significant challenge in the development of protein and peptide-based drugs is predicting their potential toxicity, which is a primary cause of failure in clinical trials. While several computational methods have been developed to address different types of toxicity, such as cytotoxicity (Beltrán et al., 2024; Cole & Brewer, 2019; Gacesa, Barlow, & Long, 2016; Gupta et al., 2013; Mall, Singh, Patel, Guirimand, & Castiglione, 2024; Rathore et al., 2024; Saha & Raghava, 2007a; N. Sharma, Naorem, Jain, & Raghava, 2022; Wei, Ye, Sakurai, Mu, & Wei, 2022; Wei, Ye, Xue, Sakurai, & Wei, 2021; Wong, Hardy, Wood, Bailey, & King, 2013), hemolytic toxicity (Chaudhary et al., 2016; Guntuboina, Das, Mollaei, Kim, & Barati Farimani, 2023; Hasan et al., 2020; Kumar, Kumar, Agrawal, Patiyal, & Raghava, 2020; Rathore, Kumar, Choudhury, Mehta, & Raghava, 2025; Salem, Keshavarzi Arshadi, & Yuan, 2022; Timmons & Hewage, 2020; Win et al., 2017), neurotoxicity (Chaohong, 2012; Lee et al., 2021; Mei & Zhao, 2018; Rathore, Jain, Choudhury, & Raghava, 2025; Saha & Raghava, 2007b; Tang et al., 2017), and allergenicity (Basith, Pham, Manavalan, & Lee, 2024; Dimitrov, Flower, & Doytchinova, 2013; Dimitrov, Naneva, Doytchinova, & Bangov, 2014; Garcia-Moreno & Gutiérrez-Naranjo, 2022; Maurer-Stroh et al., 2019; Muh, Tong, & Tammi, 2009; Nguyen et al., 2022; Saha & Raghava, 2006; N. Sharma et al., 2021; Wang, Yu, Zhao, Zhang, & Li, 2013). These methods enable initial safety assessments by predicting toxic or hemolytic peptides in silico (Figure 7). Parallel endeavors have also facilitated the creation of in silico prediction tools for other therapeutic classes of peptides, including immunomodulatory peptides capable of stimulating antigen-presenting cells (APCs) for vaccine development (Nagpal, Chaudhary, Agrawal, & Raghava, 2018), cell-penetrating peptides for improved intracellular drug delivery (Gautam, Chaudhary, Kumar, & Raghava, 2015), and anti-angiogenic peptides for cancer treatment (Ettayapuram Ramaprasad, Singh, Gajendra P S, & Venkatesan, 2015). These examples demonstrate growing dependability and utility of computer tools for peptide classification and safety assessment. However, there has been very limited work focused specifically on predicting ion channel toxins, which are still key determinants of peptide safety and therapeutic appropriateness. Here, we fill this void by describing IonNTxPred, a systematic and comprehensive prediction model for ion channel-modulating proteins. Our findings identified distinct molecular signatures that differentiate modulators from non-modulators and are critical to establishing safer biologic design and biosafety evaluation.

**Figure 7:**
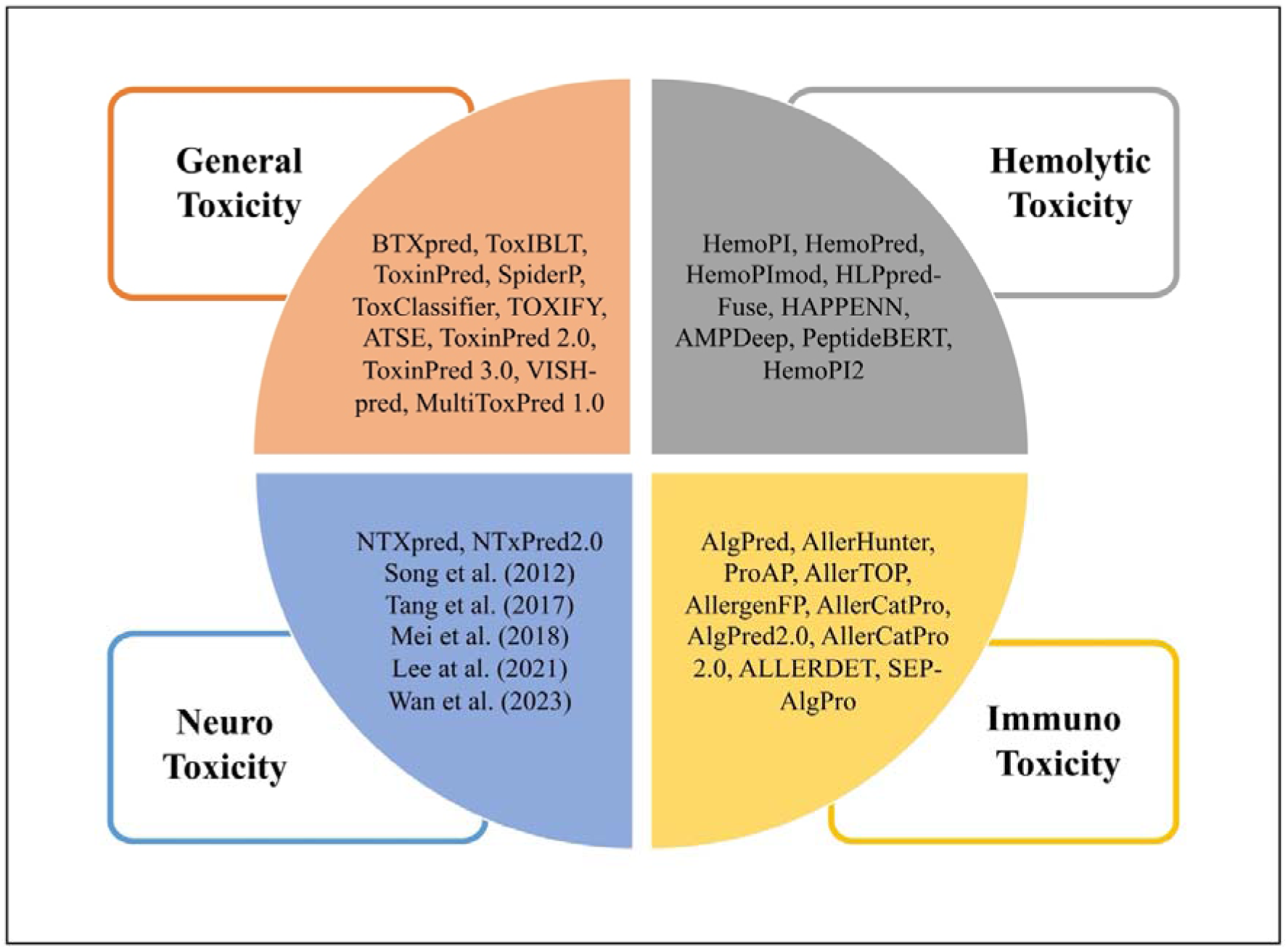
In silico tools for predicting various forms of peptide toxicities.

Predicting ion channel-modulating proteins is not merely an academic exercise but a practical safeguard across drug discovery, synthetic biology, and the broader bioeconomy. Ion channels (Na□, K□, Ca²□, etc.) regulate excitability in the heart, brain, and muscle, and unintended modulation can precipitate life-threatening outcomes such as arrhythmias, seizures, or respiratory failure. In therapeutics, early in silico triage of peptide and protein candidates helps flag hERG-like liabilities and other off-target channel effects before costly functional assays, thereby shortening development timelines, reducing reliance on animal testing, and directing experimental resources toward safer leads (Farm et al., 2023). Importantly, the clinical success of venom-derived ion channel modulators highlights the translational potential of this approach. For example, ω-conotoxin MVIIA (Rubiu et al., 2024), dalazatide (ShK-186), a sea anemone–derived Kv1.3 inhibitor, has advanced to human trials for autoimmune diseases (Cañas, Castaño-Valencia, & Castro-Herrera, 2022; Tyagi et al., 2022), while spider peptide toxins such as huwentoxin-IV (Liu et al., 2014) and protoxin-II (JBC) are being explored as non-opioid analgesics targeting NaV1.7 channels (Torres-Pérez et al., 2018) [29063143]. Beyond drug development, predictive screening is also critical for biosafety, where engineered proteins and genetically modified crops must be evaluated to avoid mammalian ion channel interference and minimize harm to non-target organisms such as pollinators and aquatic species.

In this study, we developed IonNTxPred, a robust and comprehensive framework for the prediction of ion channel-modulating proteins. Our analysis of ion channel-modulating proteins revealed clear molecular signatures that distinguish modulators from non-modulators. Compositional profiling showed that residues such as cysteine, glycine, lysine, and tyrosine were consistently enriched in toxic proteins, with cysteine emerging as the most discriminative feature, reflecting its role in forming disulfide bridges that stabilize active conformations. Positional analysis further highlighted that critical motifs were preferentially enriched at the C-terminal, suggesting structural and functional hotspots relevant for channel interaction. Importantly, feature importance analysis using the ET classifier independently reinforced these findings: cysteine-related descriptors and composition-based features consistently ranked among the top predictors across all ion channel datasets. The convergence of compositional, positional, and machine learning-driven feature importance underscores the pivotal role of cysteine-richness and sequence order in defining modulatory potential, providing both mechanistic insight and computational validation of our models.

Although the simple composition of cysteine showed strong predictive performance, it is not an absolute criterion. On average, a threshold of ∼7.3% cysteine content can discriminate between ion channel modulators and non-modulators: proteins below this threshold are more likely to be non-modulators, whereas those above it are likely modulators. However, this rule must be applied with caution. Certain toxins such as Glycyl-poneratoxin, a 26-residue peptide (FLPLLILGSLLMTPPVIQAIHDAQRG), contain no cysteine residues at all yet display potent modulatory activity by converting transient sodium currents into persistent ones, particularly on Nav1.6/SCN8A channels and, to a lesser extent, Nav1.7/SCN9A (Johnson, Rikli, Schmidt, & Evans, 2017; Robinson et al., 2023). Conversely, some proteins such as Metallothionein-3, a 68-residue astrocytic protein, contain nearly 30% cysteine yet do not function as ion channel modulators (Garrett, Sens, Todd, Somji, & Sens, 1999; You et al., 2002). These contrasting examples emphasize that while cysteine enrichment is a robust signal of modulatory potential, robust prediction requires the integration of additional sequence and structural descriptors, best achieved through ML and LLM-based approaches, rather than relying on cysteine composition alone.

This study introduces IonNTxPred, a comprehensive framework for predicting ion channel modulators by integrating both alignment-based and alignment-free computational approaches.

Our alignment-based methods, including BLAST and MERCI motif search, successfully identified true positive modulators by uncovering distinct sequence patterns that differentiate toxic from non-toxic proteins. Although BLAST reliably captures many true positives, its performance is hampered by low specificity and a high false-positive rate, resulting in limited overall coverage. The low coverage of BLAST and MERCI on the non-redundant independent set is expected, as the 40% identity threshold deliberately removes close homologs; this design choice limits the effectiveness of similarity-based methods and highlights the necessity of alignment-free approaches like Ml and PLMs. Similarly, the motif search revealed that while toxic motifs, particularly in sodium and potassium channel modulators, offer high sensitivity, their specificity is often compromised by overlaps with motifs in non-toxic sequences, whereas motifs consistently display high specificity.

In contrast, our alignment-free approach, encompassing machine learning models based on traditional compositional features (AAC, DPC, ALLCOMP) and state-of-the-art PLMs, achieved superior predictive performance. Notably, while the ALLCOMP feature set marginally improved accuracy and AUC compared to simpler feature sets, these gains came at the cost of a dramatic increase in feature dimensionality with only incremental improvements in sensitivity and MCC. PLM-based models, particularly those utilizing the ESM2-t33 architecture, demonstrated consistently high sensitivity, specificity, and near-perfect AUC values across all datasets, underscoring the efficacy of deep contextual embeddings in capturing subtle sequence and evolutionary signals without extensive feature engineering. Furthermore, our hybrid approach combining BLAST search and PLM predictions further enhanced the balance between true positive and negative rates, as evidenced by improved MCC scores. Despite these advances, challenges remain, including the need to further optimize MCC, reduce the computational burden of high-dimensional models, and enhance model interpretability. Our cross-dataset analysis revealed strong generalization across ion channel classes. This observation underscores that predictive models are capturing fundamental biophysical principles of ion channel interaction, rather than dataset-specific biases. Moreover, our findings highlight the importance of channel-specific modeling, given the observed variability in predictive performance among Na□, K□, Ca²□, and other channel modulators.

In summary, the integration of diverse computational methodologies in IonNTxPred significantly advances the prediction of ion channel modulators. While traditional ML models based on AAC offer a solid baseline, PLM-based approaches, particularly those leveraging ESM2-t33, consistently outperform them across multiple evaluation metrics. The hybrid model not only capitalizes on the strengths of similarity searches and motif-based predictions but also provides a more balanced and robust framework for protein classification tasks related to ion channel modulation. Future work should focus on expanding training datasets, incorporating structure-based analyses, and refining predictions to distinguish between different modes of channel modulation, which are crucial for therapeutic applications.

The high cross-dataset performance can be partly explained by the biological reality that many ion channel toxins exhibit moonlighting activity, modulating multiple ion channel types rather than acting in isolation. Such multifunctional peptides naturally introduce overlap between datasets, since the same sequence may appear in more than one ion channel class. Although we carefully split each dataset into independent training and testing sets, it remains possible that a peptide present in the training set of one channel class also appears in the testing set of another. This overlap, together with the conserved structural features shared by ion channel modulators, likely contributes to the strong cross-dataset generalization observed. Rather than being a limitation, this underscores the biological insight that ion channel modulators are not always strictly selective, and predictive models should therefore account for their multi-channel activity.

Tools like IonNTxPred make this actionable by coupling a language model with motif/similarity evidence to (i) classify modulators vs non-modulators with moonlighting function, (ii) localize toxic segments via protein scanning, and (iii) guide sequence redesign toward reduced channel engagement while preserving function. This granular view, where risk is tied to specific regions rather than entire proteins, enables rational edits and transparent safety dossiers. Our genome-wide predictions across diverse venomous organisms highlight the broad usability and scalability of IonNTxPred, demonstrating its ability to efficiently screen entire proteomes for ion channel modulators. This bulk application not only uncovers novel candidate toxins but also underscores the tool’s potential in large-scale drug discovery, biosafety evaluation, and comparative toxicology.

## 5 Limitations and Future Perspectives

One key limitation of our study is the relatively small and recent nature of the datasets used for training, which may affect the generalizability of the results. Furthermore, IonNTxPred is currently limited to natural amino acids, as it excludes sequences containing non-canonical amino acids and chemically modified sequences. Although our approach leverages sequence-based analysis and incorporates evolutionary features, it does not account for structural details that could provide deeper insights into the binding and functional properties of proteins. The three-dimensional conformation of a biomolecule is critical for its interaction with ion channels or receptors, and structure-based analysis could reveal how such interactions facilitate channel opening or closing. In practical drug development, proteins are often chemically modified to enhance stability, bioavailability, and precision, underscoring the need for predictive tools capable of assessing the channel-modulating activity of these modified sequences. Additionally, our current model only distinguishes between modulating and non-modulating proteins, whereas a more nuanced prediction, such as differentiating between proteins that close versus those that open ion channels, would be highly beneficial for therapeutic applications. Future research should aim to integrate structural analysis and expand the dataset to include modified protein sequences, ultimately refining the predictive capabilities for drug development.

## Authors’ contributions

ASR collected and processed the datasets. ASR and SJ implemented the algorithms and developed the prediction models. ASR designed and developed the front-end user interface, while SJ and NKM contributed to the back-end development of the web server and assisted in result analysis. ASR also developed the pip package and created the Hugging Face and GitHub repositories. ASR and GPSR co-wrote the manuscript. GPSR conceived and supervised the project. All authors read and approved the final manuscript.

## Acknowledgments

The authors express their gratitude to the University Grants Commission (UGC) and the Council of Scientific and Industrial Research (CSIR) for their generous fellowships and financial support. They also thank the Department of Computational Biology, IIIT-D, New Delhi, for the excellent infrastructure and facilities. The authors would like to acknowledge the Department of Biotechnology (DBT) for the infrastructure grant awarded to the institute. The current work has been supported by the Department of Biotechnology (DBT) grant BT/PR40158/BTIS/137/24/2021. Furthermore, they would like to acknowledge BioRender.com for creating the figures used in this work and for utilizing ChatGPT to enhance the language of the manuscript.

## Author’s Biography

1. Anand Singh Rathore is currently pursuing a Ph.D. in Computational Biology at the Department of Computational Biology, Indraprastha Institute of Information Technology, New Delhi, India.
2. Saloni Jain is currently pursuing a Ph.D. in Computational Biology at the Department of Computational Biology, Indraprastha Institute of Information Technology, New Delhi, India.
3. Naman Kumar Mehta is pursuing a Ph.D. in Computational Biology at the Department of Computational Biology, Indraprastha Institute of Information Technology, New Delhi, India.
4. Gajendra P. S. Raghava is currently working as a Professor and Head of the Department of Computational Biology, Indraprastha Institute of Information Technology, New Delhi, India.

## Data Availability

The datasets generated for this study can be accessed on the ’IonNTXPred’ web server at https://webs.iiitd.edu.in/raghava/ionntxpred/download.html, publicly available on GitHub https://github.com/raghavagps/IonNTXPred.

## Code availability

The source code for this study is publicly available on GitHub and can be found at https://github.com/raghavagps/IonNTXPred, the ’IonNTXPred’ web server at https://webs.iiitd.edu.in/raghava/ionntxpred/download.html and Hugging Face https://huggingface.co/raghavagps-group/IonNTxPred.

