## Supplementary Data for "A PLM-Based Method for Predicting Protein Ion Channel Modulators for Drug Discovery and Safety Evaluation"

### LLM-based Prediction of Ion Channel Toxins with Moonlighting Functions

**Mailing Address of Authors**

[*]**Corresponding Author**

Prof. Gajendra P. S. Raghava

Head and Professor

Department of Computational Biology

Indraprastha Institute of Information Technology, Delhi

Okhla Industrial Estate, Phase III, (Near Govind Puri Metro Station)

New Delhi, India – 110020

Office: A-302 (R&D Block)

Website: http://webs.iiitd.edu.in/raghava/

### Index Table


| Table | Content | Page No. |
|---|---|---|
| ***Supplementary Table S1*** | List of all the composition-based features computed from Pfeature standalone along with vector length. | 3 |
| ***Supplementary Figure S1*** | Average amino acid composition of all ion channel modulator and non-madulator protein in different datasets. | 4 |
| ***Supplementary Figure S2*** | Residue preferences at different positions in ion channel modulators and non-modulator proteins. | 4 |
| ***Supplementary Table S2*** | Amino Acid Composition-based Prediction. | 8 |
| ***Supplementary Table S3*** | BLAST similarity search results for training and independent dataset, where the blast database consists of sequences of the training dataset. | 11 |
| ***Supplementary Table S4*** | List of top motifs In different ion channel modulating protein dataset. | 16 |
| ***Supplementary Table S5*** | Performance of ML-based models on all four datasets using composition based features. | 19 |
| ***Supplementary Table S6*** | Performance of PLM-based models on all four datasets. | 33 |

*Supplementary Table S1: List of all the composition-based features computed from Pfeature standalone along with vector length.*

| Name of the Feature | Feature vector length |
|---|---|
| **Amino acid composition (AAC)** | 20 |
| **Dipeptide composition (DPC)** | 400 |
| **Tripeptide composition (TPC)** | 8000 |
| **Atom and Bond Type composition (ATC & BTC)** | 9 |
| **Physicochemical Properties Composition (PCP)** | 30 |
| **Residue Repeat Information (RRI)** | 20 |
| **Property Repeat Information (PRI)** | 25 |
| **Distance distribution of residue (DDR)** | 20 |
| **Shannon-Entropy of Protein (SEP)** | 1 |
| **Shannon Entropy of Physicochemical Property (SPC)** | 25 |
| **Shannon Entropy of Residues (SER)** | 20 |
| **Pseudo amino acid composition (PAAC)** | 21 |
| **Amphiphilic pseudo amino acid composition (APAAC)** | 23 |
| **Quasi-sequence order (QSO)** | 42 |
| **Sequence Order Coupling Number (SOCN)** | 2 |
| **Conjoint Triad Calculation (CTC)** | 343 |
| **Composition enhanced Transition Distribution (CeTD)** | 189 |
| **Total** | 9190 |

*Supplementary Figure S1: Average amino acid composition of all ion channel modulator and non-modulator proteins in different datasets.*

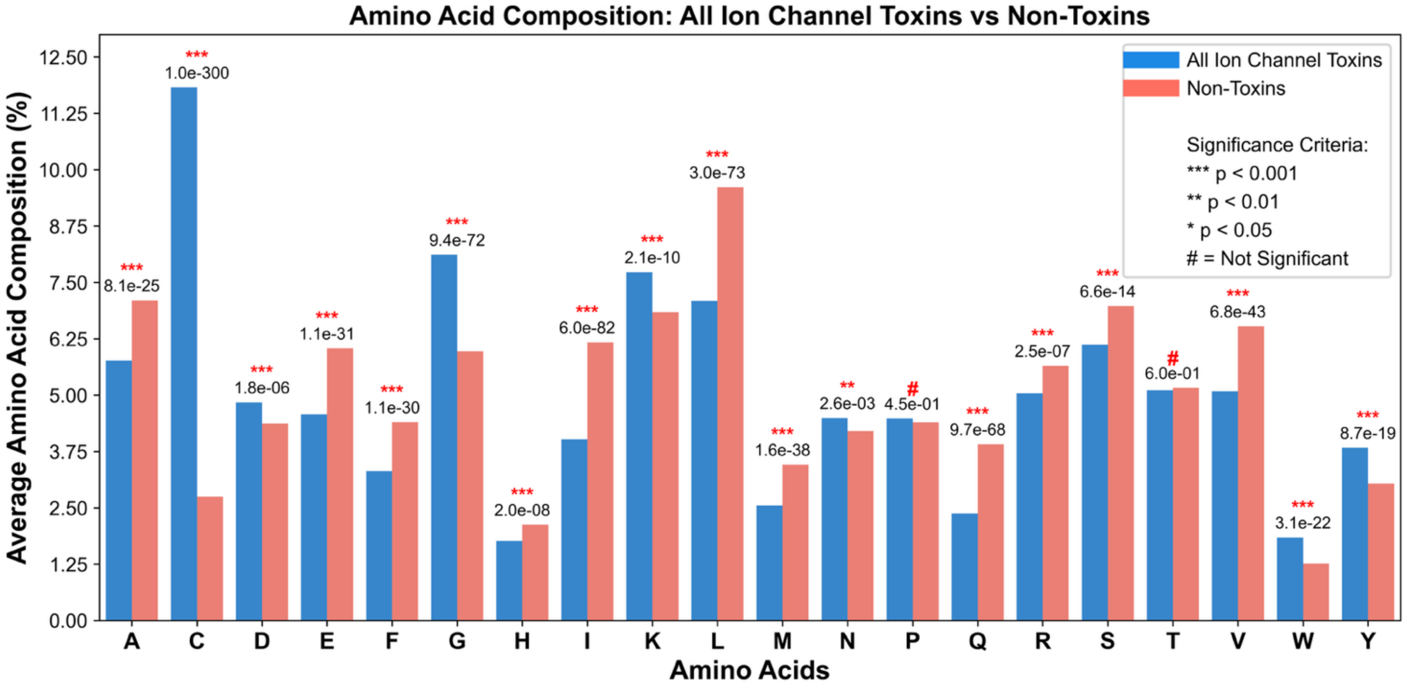


*Supplementary Figure S2: Residue preferences at different positions in ion channel modulators and non-modulator proteins.*

**Supplementary Figure S2.1: Amino acid preferences at specific positions in $Na^{+}$ ion channel modulator protein sequences, identified using the Two Sample Logo (TSL) method. (A) Displays the first 12 residues from the N-terminus; (B) displays the last 12 residues from the C-terminus.**

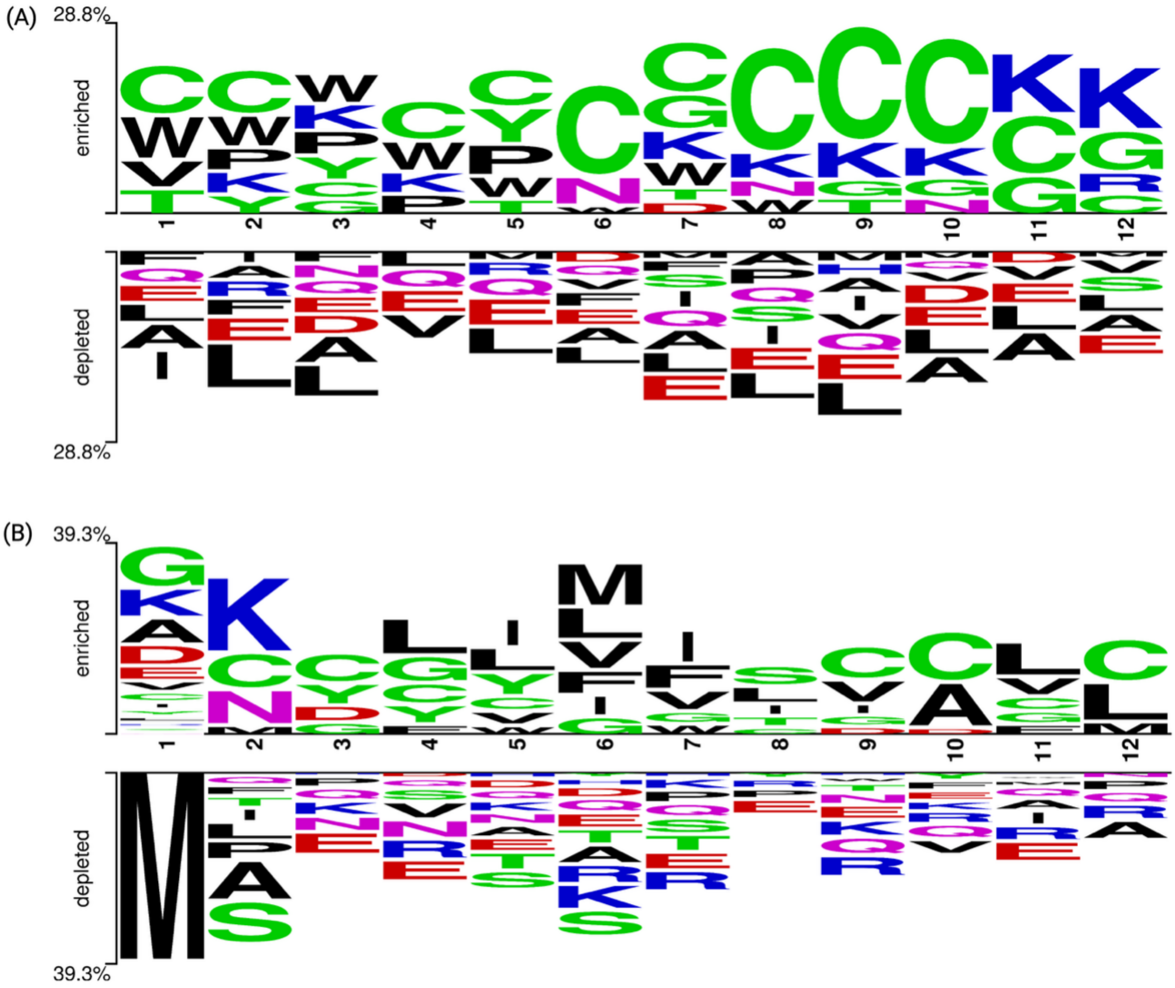


**Supplementary Figure S2.2: Amino acid preferences at specific positions in K⁺ ion channel modulator protein sequences, identified using the Two Sample Logo (TSL) method. (A) Displays the first 13 residues from the N-terminus; (B) displays the last 13 residues from the C-terminus.**

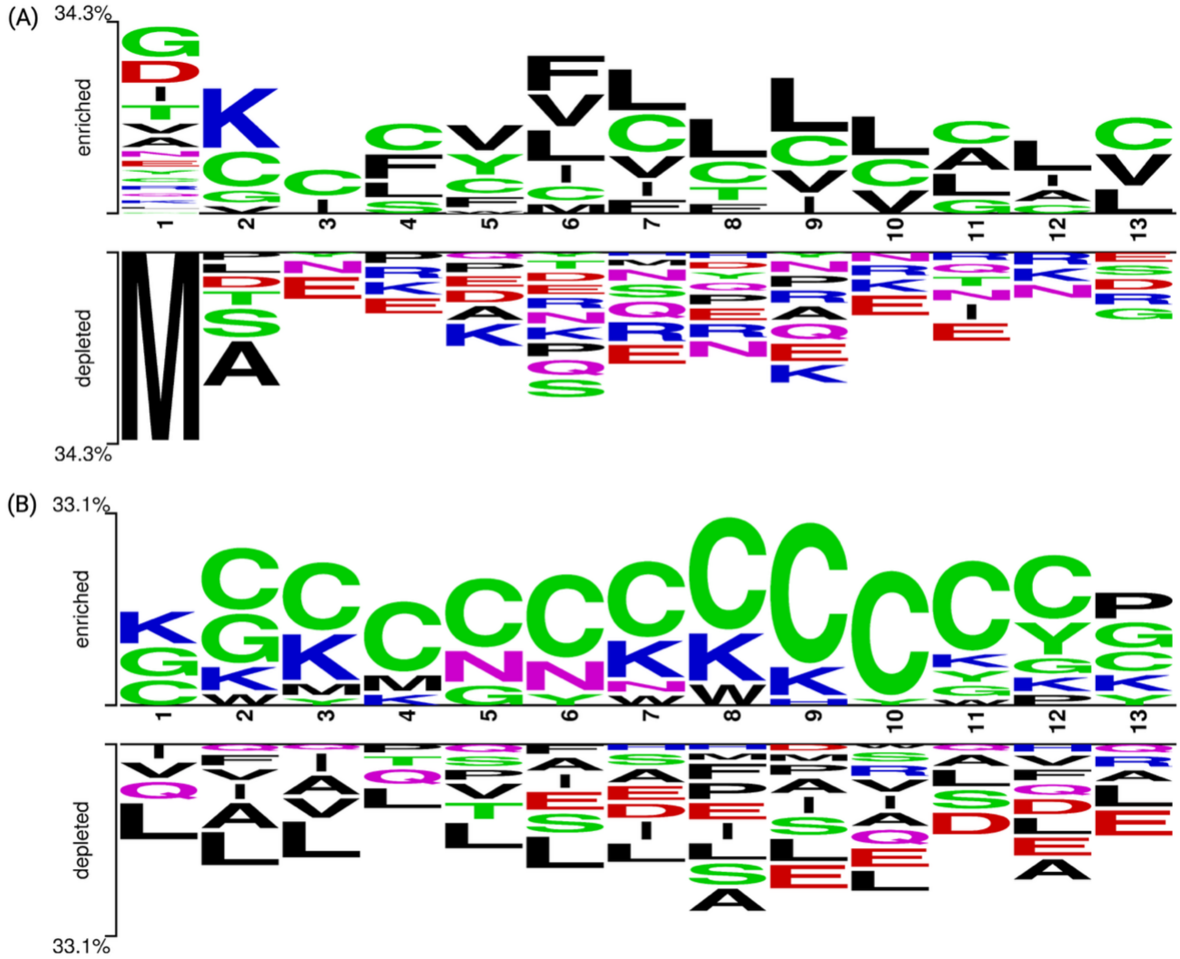


**Supplementary Figure S2.3: Amino acid preferences at specific positions in $Ca^{2+}$ ion channel modulator protein sequences, identified using the Two Sample Logo (TSL) method. (A) Displays the first 13 residues from the N-terminus; (B) displays the last 13 residues from the C-terminus.**

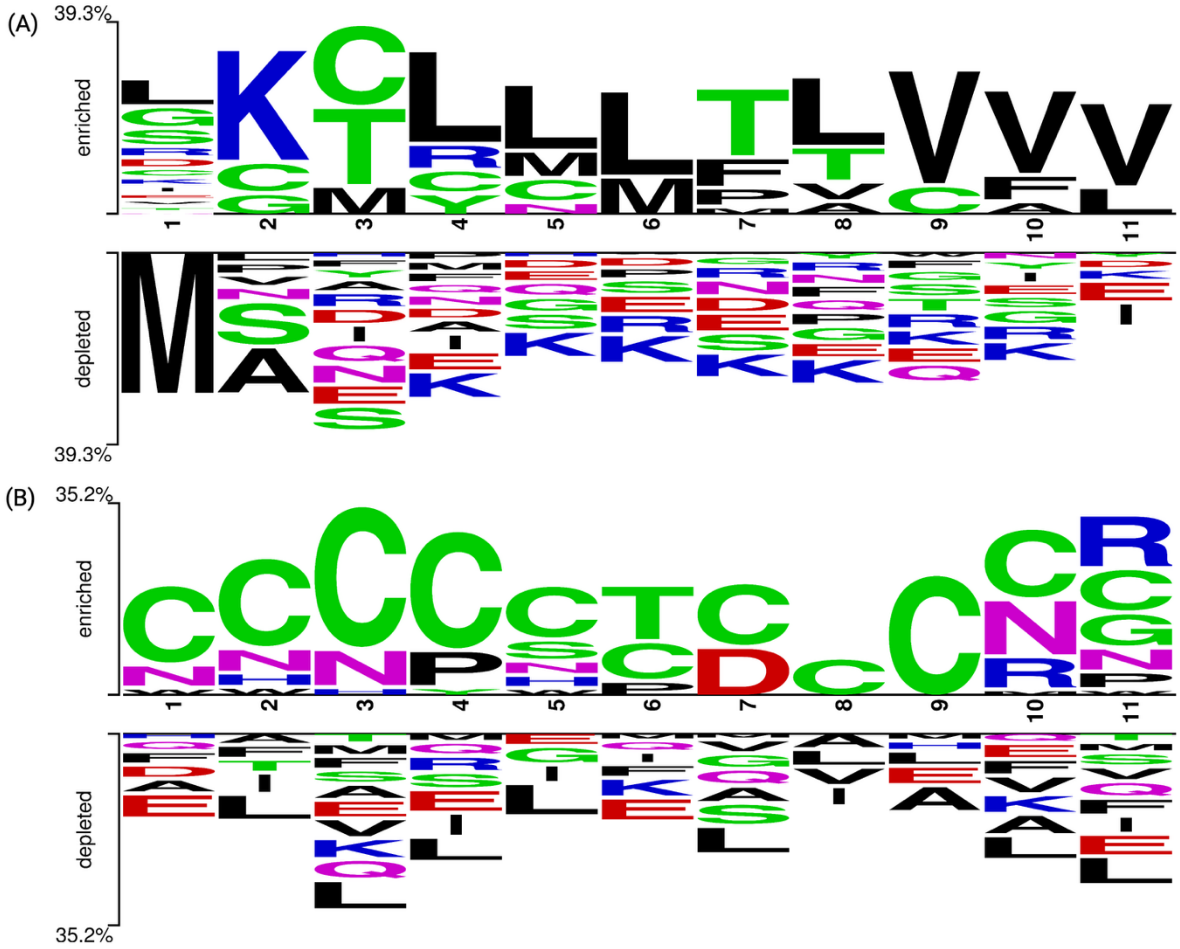


**Supplementary Figure S2.4: Amino acid preferences at specific positions in other ion channel modulator protein sequences, identified using the Two Sample Logo (TSL) method. (A) Displays the first 11 residues from the N-terminus; (B) displays the last 11 residues from the C-terminus.**

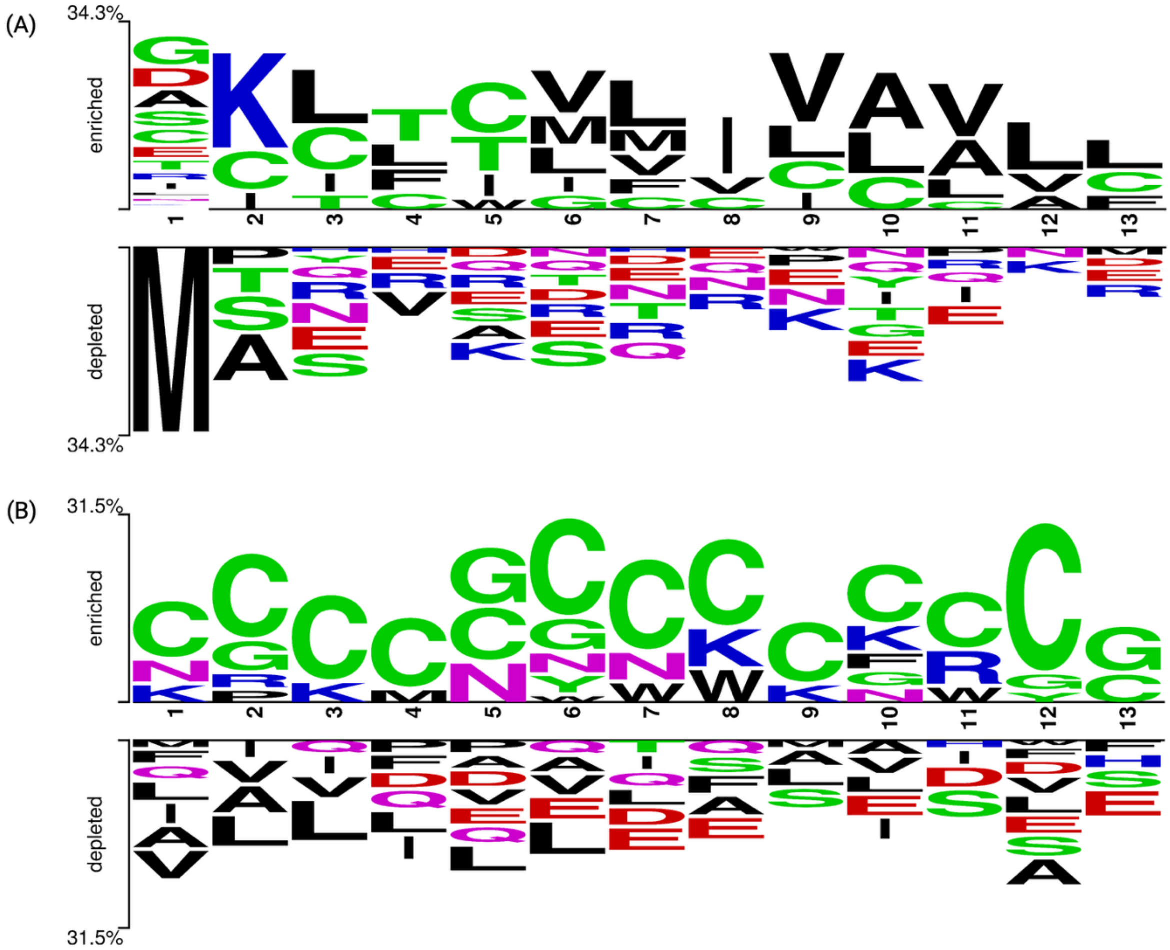


**Note: The number of residues analyzed at each terminus is based on the minimum sequence length.**

*Supplementary Table S2: Amino Acid Composition-based Prediction.*

**Supplementary Table S2.1: Amino acid composition-based prediction of Na$^+$ channel-modulating proteins and non-modulating sequences.**

| Amino Acid | Best Threshold | Mean (modulator) | Mean (Non-modulator) | Accuracy | AUC |
|---|---|---|---|---|---|
| A | 6.364 | 5.624 | 7.103 | 0.537 | 0.596 |
| C | 7.312 | 11.876 | 2.749 | 0.889 | 0.934 |

| | | | | | |
|---|---|---|---|---|---|
| **D** | 4.856 | 5.342 | 4.371 | 0.589 | 0.599 |
| **E** | 5.380 | 4.719 | 6.040 | 0.531 | 0.587 |
| **F** | 3.852 | 3.300 | 4.403 | 0.524 | 0.603 |
| **G** | 7.535 | 9.098 | 5.973 | 0.698 | 0.737 |
| **H** | 1.834 | 1.542 | 2.127 | 0.498 | 0.585 |
| **I** | 5.069 | 3.965 | 6.172 | 0.587 | 0.664 |
| **K** | 7.493 | 8.145 | 6.841 | 0.602 | 0.604 |
| **L** | 8.438 | 7.263 | 9.614 | 0.585 | 0.650 |
| **M** | 2.839 | 2.217 | 3.461 | 0.575 | 0.661 |
| **N** | 4.403 | 4.599 | 4.206 | 0.581 | 0.556 |
| **P** | 4.297 | 4.195 | 4.400 | 0.469 | 0.499 |
| **Q** | 2.836 | 1.760 | 3.912 | 0.612 | 0.730 |
| **R** | 4.912 | 4.170 | 5.653 | 0.552 | 0.607 |
| **S** | 6.429 | 5.880 | 6.978 | 0.536 | 0.579 |
| **T** | 4.645 | 4.126 | 5.164 | 0.554 | 0.601 |
| **V** | 5.628 | 4.727 | 6.530 | 0.571 | 0.642 |
| **W** | 1.935 | 2.607 | 1.263 | 0.725 | 0.700 |
| **Y** | 3.942 | 4.846 | 3.039 | 0.666 | 0.643 |

**Supplementary Table S2.2: Amino acid composition-based prediction of $K^+$ channel-modulating and non-modulating proteins.**

| Amino Acid | Best Threshold | Mean (modulator) | Mean (Non-modulator) | Accuracy | AUC |
|---|---|---|---|---|---|
| **A** | 6.298 | 5.492 | 7.103 | 0.531 | 0.597 |
| **C** | 7.312 | 11.875 | 2.749 | 0.879 | 0.932 |
| **D** | 4.337 | 4.303 | 4.371 | 0.464 | 0.505 |
| **E** | 5.218 | 4.395 | 6.040 | 0.548 | 0.616 |
| **F** | 4.081 | 3.759 | 4.403 | 0.478 | 0.545 |
| **G** | 7.034 | 8.094 | 5.973 | 0.648 | 0.669 |
| **H** | 1.977 | 1.827 | 2.127 | 0.455 | 0.535 |
| **I** | 5.320 | 4.467 | 6.172 | 0.547 | 0.625 |
| **K** | 7.839 | 8.837 | 6.841 | 0.620 | 0.629 |
| **L** | 8.055 | 6.496 | 9.614 | 0.619 | 0.693 |
| **M** | 3.135 | 2.809 | 3.461 | 0.532 | 0.579 |
| **N** | 4.326 | 4.446 | 4.206 | 0.572 | 0.540 |
| **P** | 4.332 | 4.264 | 4.400 | 0.456 | 0.500 |
| **Q** | 3.291 | 2.671 | 3.912 | 0.545 | 0.617 |
| **R** | 5.261 | 4.869 | 5.653 | 0.503 | 0.541 |
| **S** | 6.464 | 5.951 | 6.978 | 0.527 | 0.570 |
| **T** | 4.885 | 4.605 | 5.164 | 0.509 | 0.552 |

| | | | | | |
|---|---|---|---|---|---|
| **V** | 5.889 | 5.249 | 6.530 | 0.542 | 0.597 |
| **W** | 1.407 | 1.552 | 1.263 | 0.572 | 0.517 |
| **Y** | 3.537 | 4.036 | 3.039 | 0.612 | 0.599 |

**Supplementary Table S2.3: Amino acid composition-based prediction of $Ca^{2+}$ channel-modulating and non-modulating proteins.**

| Amino Acid | Best Threshold | Mean (modulator) | Mean (Non-modulator) | Accuracy | AUC |
|---|---|---|---|---|---|
| **A** | 6.692 | 6.280 | 7.103 | 0.486 | 0.529 |
| **C** | 7.124 | 11.498 | 2.749 | 0.869 | 0.910 |
| **D** | 4.627 | 4.883 | 4.371 | 0.566 | 0.552 |
| **E** | 5.370 | 4.700 | 6.040 | 0.522 | 0.595 |
| **F** | 3.854 | 3.304 | 4.403 | 0.504 | 0.591 |
| **G** | 6.884 | 7.795 | 5.973 | 0.628 | 0.652 |
| **H** | 1.865 | 1.603 | 2.127 | 0.468 | 0.567 |
| **I** | 5.139 | 4.106 | 6.172 | 0.562 | 0.654 |
| **K** | 7.306 | 7.771 | 6.841 | 0.590 | 0.564 |
| **L** | 8.366 | 7.118 | 9.614 | 0.583 | 0.654 |
| **M** | 3.079 | 2.698 | 3.461 | 0.541 | 0.595 |
| **N** | 4.368 | 4.530 | 4.206 | 0.582 | 0.550 |
| **P** | 4.280 | 4.160 | 4.400 | 0.453 | 0.497 |
| **Q** | 3.217 | 2.523 | 3.912 | 0.552 | 0.630 |
| **R** | 5.614 | 5.576 | 5.653 | 0.463 | 0.484 |
| **S** | 6.770 | 6.561 | 6.978 | 0.494 | 0.529 |
| **T** | 5.159 | 5.153 | 5.164 | 0.460 | 0.491 |
| **V** | 5.674 | 4.818 | 6.530 | 0.565 | 0.629 |
| **W** | 1.452 | 1.642 | 1.263 | 0.598 | 0.545 |
| **Y** | 3.158 | 3.278 | 3.039 | 0.549 | 0.529 |

**Supplementary Table S2.4: Amino acid composition-based prediction of other ion channel- modulating and non-modulating proteins.**

| Amino Acid | Best Threshold | Mean (modulator) | Mean (Non-modulator) | Accuracy | AUC |
|---|---|---|---|---|---|
| **A** | 6.484 | 5.865 | 7.103 | 0.520 | 0.580 |
| **C** | 7.354 | 11.959 | 2.749 | 0.877 | 0.932 |
| **D** | 4.559 | 4.748 | 4.371 | 0.558 | 0.554 |
| **E** | 5.269 | 4.498 | 6.040 | 0.536 | 0.614 |

| | | | | | |
|---|---|---|---|---|---|
| **F** | 3.632 | 2.860 | 4.403 | 0.546 | 0.644 |
| **G** | 6.553 | 7.133 | 5.973 | 0.593 | 0.616 |
| **H** | 2.116 | 2.105 | 2.127 | 0.435 | 0.505 |
| **I** | 4.873 | 3.573 | 6.172 | 0.600 | 0.700 |
| **K** | 6.407 | 5.972 | 6.841 | 0.491 | 0.544 |
| **L** | 8.552 | 7.490 | 9.614 | 0.566 | 0.622 |
| **M** | 3.028 | 2.595 | 3.461 | 0.550 | 0.620 |
| **N** | 4.292 | 4.379 | 4.206 | 0.561 | 0.525 |
| **P** | 4.859 | 5.319 | 4.400 | 0.604 | 0.582 |
| **Q** | 3.323 | 2.735 | 3.912 | 0.540 | 0.611 |
| **R** | 5.798 | 5.943 | 5.653 | 0.555 | 0.540 |
| **S** | 6.636 | 6.293 | 6.978 | 0.507 | 0.537 |
| **T** | 6.023 | 6.881 | 5.164 | 0.639 | 0.629 |
| **V** | 6.050 | 5.571 | 6.530 | 0.513 | 0.558 |
| **W** | 1.304 | 1.345 | 1.263 | 0.553 | 0.521 |
| **Y** | 2.887 | 2.736 | 3.039 | 0.495 | 0.531 |

*Supplementary Table S3: BLAST similarity search results for training and independent dataset, where the blast database consists of sequences of the training dataset.*

**Supplementary Table S3.1: BLAST similarity search results on $Na^+$ training dataset.**

| E-value | Coverage (%) | Total Hits | Modulators (Chits) | Modulators (Chits %) | Modulators (Whits) | Modulator (Whits %) | Non-modulators (Chits) | Non-modulators (Chits %) | Non-modulators (Whits) | Non-modulators (Whits %) |
|---|---|---|---|---|---|---|---|---|---|---|
| **1.00E-12** | 22.88% | 493 | 457 | 100.00% | 0 | 0.00% | 36 | 100.00% | 0 | 0.00% |
| **1.00E-11** | 24.22% | 522 | 461 | 100.00% | 0 | 0.00% | 61 | 100.00% | 0 | 0.00% |
| **1.00E-10** | 25.34% | 546 | 466 | 100.00% | 0 | 0.00% | 80 | 100.00% | 0 | 0.00% |
| **1.00E-09** | 27.38% | 590 | 472 | 100.00% | 0 | 0.00% | 118 | 100.00% | 0 | 0.00% |
| **1.00E-08** | 29.47% | 635 | 477 | 100.00% | 0 | 0.00% | 158 | 100.00% | 0 | 0.00% |
| **1.00E-07** | 31.28% | 674 | 483 | 100.00% | 0 | 0.00% | 191 | 100.00% | 0 | 0.00% |
| **1.00E-06** | 33.83% | 729 | 491 | 100.00% | 0 | 0.00% | 238 | 100.00% | 0 | 0.00% |
| **1.00E-05** | 36.19% | 780 | 495 | 100.00% | 0 | 0.00% | 285 | 100.00% | 0 | 0.00% |

| | | | | | | | | | | |
|---|---|---|---|---|---|---|---|---|---|---|
| **1.00E-04** | 38.89% | 838 | 496 | 100.00% | 0 | 0.00% | 342 | 100.00% | 0 | 0.00% |
| **1.00E-03** | 40.97% | 883 | 501 | 100.00% | 0 | 0.00% | 378 | 98.95% | 4 | 1.05% |
| **1.00E-02** | 43.11% | 929 | 503 | 100.00% | 0 | 0.00% | 416 | 97.65% | 10 | 2.35% |
| **1.00E-01** | 48.86% | 1053 | 506 | 100.00% | 0 | 0.00% | 515 | 94.15% | 32 | 5.85% |
| **1** | 72.53% | 1563 | 508 | 99.80% | 1 | 0.20% | 924 | 87.67% | 130 | 12.33% |
| **10** | 99.35% | 2141 | 509 | 99.41% | 3 | 0.59% | 1398 | 85.82% | 231 | 14.18% |
| **100** | 100.00% | 2155 | 509 | 99.41% | 3 | 0.59% | 1406 | 85.58% | 237 | 14.42% |

(Note: Chits = correct hits, Whits = wrong/incorrect hits.)

#### Supplementary Table S3.2: BLAST similarity search results on $Na^+$ independent dataset.

| E-value | Coverage (%) | Total Hits | Modulators (Chits) | Modulators (Chits %) | Modulators (Whits) | Modulator (Whits %) | Non-modulators (Chits) | Non-modulators (Chits %) | Non-modulators (Whits) | Non-modulators (Whits %) |
|---|---|---|---|---|---|---|---|---|---|---|
| **1.00E-12** | 9.46% | 51 | 31 | 96.88% | 1 | 3.12% | 19 | 100.00% | 0 | 0.00% |
| **1.00E-11** | 11.50% | 62 | 36 | 97.30% | 1 | 2.70% | 25 | 100.00% | 0 | 0.00% |
| **1.00E-10** | 14.47% | 78 | 40 | 95.24% | 2 | 4.76% | 35 | 97.22% | 1 | 2.78% |
| **1.00E-09** | 17.07% | 92 | 43 | 93.48% | 3 | 6.52% | 45 | 97.83% | 1 | 2.17% |
| **1.00E-08** | 19.48% | 105 | 49 | 92.45% | 4 | 7.55% | 51 | 98.08% | 1 | 1.92% |
| **1.00E-07** | 22.08% | 119 | 53 | 93.00% | 4 | 7.00% | 61 | 98.39% | 1 | 1.61% |
| **1.00E-06** | 25.23% | 136 | 59 | 93.65% | 4 | 6.35% | 72 | 98.63% | 1 | 1.37% |
| **1.00E-05** | 27.83% | 150 | 62 | 92.54% | 5 | 7.46% | 82 | 98.80% | 1 | 1.20% |
| **1.00E-04** | 29.50% | 159 | 64 | 92.75% | 5 | 7.25% | 89 | 98.89% | 1 | 1.11% |
| **1.00E-03** | **33.77%** | **182** | **74** | **93.67%** | **5** | **6.33%** | **102** | **99.03%** | **1** | **0.97%** |
| **1.00E-02** | 37.66% | 203 | 79 | 90.80% | 8 | 9.20% | 112 | 96.55% | 4 | 3.45% |
| **1.00E-01** | 43.78% | 236 | 82 | 91.11% | 8 | 8.89% | 137 | 93.85% | 9 | 6.16% |
| **1** | 72.73% | 392 | 98 | 85.22% | 17 | 14.78% | 245 | 88.45% | 32 | 11.55% |
| **10** | 99.44% | 536 | 104 | 81.25% | 24 | 18.75% | 346 | 84.79% | 62 | 15.21% |
| **100** | 100.00% | 539 | 102 | 79.07% | 26 | 20.93% | 351 | 85.41% | 60 | 14.59% |

(Note: Chits = correct hits, Whits = wrong/incorrect hits.)

#### Supplementary Table S3.3: BLAST similarity search results on $K^+$ training dataset.

| E-value | Coverage (%) | Total Hits | Modulators (Chits) | Modulators (Chits %) | Modulators (Whits) | Modulator (Whits %) | Non-modulators (Chits) | Non-modulators (Chits %) | Non-modulators (Whits) | Non-modulators (Whits %) |
|---|---|---|---|---|---|---|---|---|---|---|
| **1.00E-12** | 19.70% | 409 | 371 | 100.00% | 0 | 0.00% | 37 | 97.37% | 1 | 2.63% |
| **1.00E-11** | 21.24% | 441 | 379 | 100.00% | 0 | 0.00% | 61 | 98.39% | 1 | 1.61% |
| **1.00E-10** | 22.45% | 466 | 385 | 100.00% | 0 | 0.00% | 80 | 98.77% | 1 | 1.23% |
| **1.00E-09** | 24.61% | 511 | 391 | 100.00% | 0 | 0.00% | 119 | 99.17% | 1 | 0.83% |
| **1.00E-08** | 26.73% | 555 | 396 | 100.00% | 0 | 0.00% | 158 | 99.37% | 1 | 0.63% |
| **1.00E-07** | 28.65% | 595 | 402 | 100.00% | 0 | 0.00% | 192 | 99.48% | 1 | 0.52% |
| **1.00E-06** | 30.97% | 643 | 402 | 100.00% | 0 | 0.00% | 239 | 99.17% | 2 | 0.83% |
| **1.00E-05** | 33.52% | 696 | 406 | 100.00% | 0 | 0.00% | 287 | 98.97% | 3 | 1.03% |
| **1.00E-04** | 36.46% | 757 | 411 | 100.00% | 0 | 0.00% | 342 | 98.84% | 4 | 1.16% |
| **1.00E-03** | 38.52% | 800 | 416 | 100.00% | 0 | 0.00% | 378 | 98.44% | 6 | 1.56% |
| **1.00E-02** | 40.85% | 848 | 418 | 100.00% | 0 | 0.00% | 417 | 96.99% | 13 | 3.02% |
| **1.00E-01** | 46.79% | 971 | 423 | 100.00% | 0 | 0.00% | 518 | 94.53% | 30 | 5.48% |
| **1** | 72.42% | 1503 | 426 | 99.77% | 1 | 0.23% | 946 | 87.91% | 130 | 12.09% |
| **10** | 99.57% | 2067 | 426 | 98.38% | 7 | 1.62% | 1408 | 85.03% | 226 | 14.97% |
| **100** | 100.00% | 2076 | 426 | 98.38% | 7 | 1.62% | 1410 | 85.94% | 233 | 14.06% |

(Note: Chits = correct hits, Whits = wrong/incorrect hits.)

#### Supplementary Table S3.4: BLAST similarity search results on $K^+$ independent dataset.

| E-value | Coverage (%) | Total Hits | Modulators (Chits) | Modulators (Chits %) | Modulators (Whits) | Modulator (Whits %) | Non-modulators (Chits) | Non-modulators (Chits %) | Non-modulators (Whits) | Non-modulators (Whits %) |
|---|---|---|---|---|---|---|---|---|---|---|
| **1.00E-12** | 8.27% | 43 | 24 | 100.00% | 0 | 0.00% | 19 | 100.00% | 0 | 0.00% |
| **1.00E-11** | 10.00% | 52 | 27 | 100.00% | 0 | 0.00% | 25 | 100.00% | 0 | 0.00% |
| **1.00E-10** | 12.69% | 66 | 31 | 100.00% | 0 | 0.00% | 35 | 100.00% | 0 | 0.00% |
| **1.00E-09** | 14.81% | 77 | 32 | 100.00% | 0 | 0.00% | 45 | 100.00% | 0 | 0.00% |
| **1.00E-08** | 17.12% | 89 | 38 | 100.00% | 0 | 0.00% | 51 | 100.00% | 0 | 0.00% |
| **1.00E-07** | 19.81% | 103 | 40 | 95.24% | 2 | 4.76% | 61 | 100.00% | 0 | 0.00% |
| **1.00E-06** | 22.50% | 117 | 43 | 95.56% | 2 | 4.44% | 72 | 100.00% | 0 | 0.00% |
| **1.00E-05** | 25.19% | 131 | 47 | 95.92% | 2 | 4.08% | 82 | 100.00% | 0 | 0.00% |

| | | | | | | | | | | |
|---|---|---|---|---|---|---|---|---|---|---|
| **1.00E-04** | 27.50% | 143 | 52 | 96.30% | 2 | 3.70% | 89 | 100.00% | 0 | 0.00% |
| **1.00E-03** | **31.92%** | **166** | **61** | **95.31%** | **3** | **4.69%** | **102** | **100.00%** | **0** | **0.00%** |
| **1.00E-02** | 35.58% | 185 | 66 | 92.96% | 5 | 7.04% | 112 | 98.25% | 2 | 1.75% |
| **1.00E-01** | 44.23% | 230 | 73 | 91.25% | 7 | 8.75% | 137 | 91.34% | 13 | 8.66% |
| **1** | 71.73% | 373 | 80 | 86.96% | 12 | 13.04% | 245 | 87.18% | 36 | 12.82% |
| **10** | 99.23% | 516 | 86 | 78.90% | 23 | 21.10% | 349 | 85.75% | 58 | 14.25% |
| **100** | 100.00% | 520 | 84 | 77.06% | 25 | 22.94% | 357 | 86.87% | 54 | 13.13% |

#### Supplementary Table S3.5: BLAST similarity search results on $Ca^{+}$ training dataset.

| E-value | Coverage (%) | Total Hits | Modulators (Chits) | Modulators (Chits %) | Modulators (Whits) | Modulator (Whits %) | Non-modulators (Chits) | Non-modulators (Chits %) | Non-modulators (Whits) | Non-modulators (Whits %) |
|---|---|---|---|---|---|---|---|---|---|---|
| **1.00E-12** | 14.30% | 277 | 240 | 100.00% | 0 | 0.00% | 37 | 100.00% | 0 | 0.00% |
| **1.00E-11** | 16.15% | 313 | 252 | 100.00% | 0 | 0.00% | 61 | 100.00% | 0 | 0.00% |
| **1.00E-10** | 17.29% | 335 | 254 | 100.00% | 0 | 0.00% | 81 | 100.00% | 0 | 0.00% |
| **1.00E-09** | 19.57% | 379 | 258 | 100.00% | 0 | 0.00% | 120 | 99.17% | 1 | 0.83% |
| **1.00E-08** | 21.99% | 426 | 265 | 100.00% | 0 | 0.00% | 160 | 99.38% | 1 | 0.62% |
| **1.00E-07** | 24.05% | 466 | 270 | 100.00% | 0 | 0.00% | 195 | 99.49% | 1 | 0.51% |
| **1.00E-06** | 26.58% | 515 | 273 | 100.00% | 0 | 0.00% | 241 | 99.59% | 1 | 0.41% |
| **1.00E-05** | 29.18% | 565 | 276 | 100.00% | 0 | 0.00% | 288 | 99.65% | 1 | 0.35% |
| **1.00E-04** | 32.26% | 625 | 281 | 100.00% | 0 | 0.00% | 341 | 99.13% | 3 | 0.87% |
| **1.00E-03** | **34.33%** | **665** | **284** | **100.00%** | **0** | **0.00%** | **377** | **98.95%** | **4** | **1.05%** |
| **1.00E-02** | 36.76% | 712 | 284 | 100.00% | 0 | 0.00% | 417 | 97.43% | 11 | 2.57% |
| **1.00E-01** | 43.31% | 839 | 287 | 100.00% | 0 | 0.00% | 523 | 94.75% | 29 | 5.25% |
| **1** | 70.26% | 1361 | 290 | 100.00% | 0 | 0.00% | 964 | 90.01% | 107 | 9.99% |
| **10** | 99.59% | 1929 | 289 | 98.29% | 5 | 1.70% | 1446 | 88.45% | 189 | 11.55% |
| **100** | 100.00% | 1937 | 289 | 98.29% | 5 | 1.70% | 1451 | 88.31% | 192 | 11.69% |

(Note: Chits = correct hits, Whits = wrong/incorrect hits.)

#### Supplementary Table S3.6: BLAST similarity search results on $Ca^{+}$ independent dataset.

| E-value | Coverage (%) | Total Hits | Modulators (Chits) | Modulators (Chits %) | Modulators (Whits) | Modulator (Whits %) | Non-modulators (Chits) | Non-modulators (Chits %) | Non-modulators (Whits) | Non-modulators (Whits %) |
|---|---|---|---|---|---|---|---|---|---|---|

| | | | | | | | | | | |
|---|---|---|---|---|---|---|---|---|---|---|
| **1.00E -12** | 5.77% | 28 | 8 | 88.89% | 1 | 11.11% | 19 | 100.00% | 0 | 0.00% |
| **1.00E -11** | 7.22% | 35 | 9 | 90.00% | 1 | 10.00% | 25 | 100.00% | 0 | 0.00% |
| **1.00E -10** | 9.69% | 47 | 11 | 91.67% | 1 | 8.33% | 35 | 100.00% | 0 | 0.00% |
| **1.00E -09** | 11.75% | 57 | 11 | 91.67% | 1 | 8.33% | 45 | 100.00% | 0 | 0.00% |
| **1.00E -08** | 13.20% | 64 | 12 | 92.31% | 1 | 7.69% | 51 | 100.00% | 0 | 0.00% |
| **1.00E -07** | 15.46% | 75 | 13 | 92.86% | 1 | 7.14% | 61 | 100.00% | 0 | 0.00% |
| **1.00E -06** | 18.76% | 91 | 17 | 94.44% | 1 | 5.56% | 73 | 100.00% | 0 | 0.00% |
| **1.00E -05** | 21.65% | 105 | 22 | 95.65% | 1 | 4.35% | 82 | 100.00% | 0 | 0.00% |
| **1.00E -04** | 23.71% | 115 | 25 | 96.15% | 1 | 3.85% | 89 | 100.00% | 0 | 0.00% |
| **1.00E -03** | **27.63%** | **134** | **30** | **96.77%** | **1** | **3.23%** | **102** | **99.03%** | **1** | **0.97%** |
| **1.00E -02** | 32.16% | 156 | 38 | 92.68% | 3 | 7.32% | 112 | 97.39% | 3 | 2.61% |
| **1.00E -01** | 41.44% | 201 | 47 | 90.38% | 5 | 9.62% | 139 | 93.29% | 10 | 6.71% |
| **1** | 70.93% | 344 | 53 | 81.54% | 12 | 18.46% | 249 | 89.25% | 30 | 10.75% |
| **10** | 99.59% | 483 | 56 | 75.68% | 18 | 24.32% | 357 | 86.01% | 52 | 13.99% |
| **100** | 100.00% | 485 | 55 | 74.32% | 19 | 25.68% | 364 | 88.57% | 47 | 11.43% |

(Note: Chits = correct hits, Whits = wrong/incorrect hits.)

#### Supplementary Table S3.7: BLAST similarity search results on other training dataset.

| E-value | Coverage (%) | Total Hits | Modulators (Chits) | Modulators (Chits %) | Modulators (Whits) | Modulator (Whits %) | Non-modulators (Chits) | Non-modulators (Chits %) | Non-modulators (Whits) | Non-modulators (Whits %) |
|---|---|---|---|---|---|---|---|---|---|---|
| **1.00E -12** | 18.67% | 382 | 344 | 100.00% | 0 | 0.00% | 37 | 97.37% | 1 | 2.63% |
| **1.00E -11** | 20.04% | 410 | 348 | 100.00% | 0 | 0.00% | 61 | 98.39% | 1 | 1.61% |
| **1.00E -10** | 20.96% | 429 | 348 | 100.00% | 0 | 0.00% | 80 | 98.77% | 1 | 1.23% |
| **1.00E -09** | 23.02% | 471 | 351 | 100.00% | 0 | 0.00% | 119 | 99.17% | 1 | 0.83% |
| **1.00E -08** | 25.17% | 515 | 354 | 100.00% | 0 | 0.00% | 160 | 99.38% | 1 | 0.62% |
| **1.00E -07** | 26.98% | 552 | 358 | 100.00% | 0 | 0.00% | 193 | 99.48% | 1 | 0.52% |
| **1.00E -06** | 29.57% | 605 | 364 | 100.00% | 0 | 0.00% | 240 | 99.59% | 1 | 0.41% |
| **1.00E -05** | 32.06% | 656 | 367 | 100.00% | 0 | 0.00% | 287 | 99.31% | 2 | 0.69% |
| **1.00E -04** | 35.24% | 721 | 377 | 100.00% | 0 | 0.00% | 342 | 99.42% | 2 | 0.58% |
| **1.00E -03** | 37.15% | 760 | 380 | 100.00% | 0 | 0.00% | 378 | 99.47% | 2 | 0.53% |

| | | | | | | | | | | |
|---|---|---|---|---|---|---|---|---|---|---|
| **1.00E-02** | 39.59% | 810 | 388 | 100.00% | 0 | 0.00% | 418 | 99.05% | 4 | 0.95% |
| **1.00E-01** | 45.71% | 935 | 394 | 100.00% | 0 | 0.00% | 523 | 96.67% | 18 | 3.33% |
| **1** | 71.01% | 1453 | 395 | 99.50% | 2 | 0.50% | 950 | 89.96% | 106 | 10.04% |
| **10** | 99.27% | 2031 | 395 | 98.50% | 6 | 1.50% | 1435 | 88.03% | 195 | 11.97% |
| **100** | 100.00% | 2046 | 396 | 98.26% | 7 | 1.74% | 1446 | 88.01% | 197 | 11.99% |

**Supplementary Table S3.8: BLAST similarity search results on other independent dataset.**

| E-value | Coverage (%) | Total Hits | Modulators (Chits) | Modulators (Chits %) | Modulators (Whits) | Modulator (Whits %) | Non-modulators (Chits) | Non-modulators (Chits %) | Non-modulators (Whits) | Non-modulators (Whits %) |
|---|---|---|---|---|---|---|---|---|---|---|
| **1.00E-12** | 6.64% | 34 | 14 | 100.00% | 0 | 0.00% | 19 | 95.00% | 1 | 5.00% |
| **1.00E-11** | 8.40% | 43 | 17 | 100.00% | 0 | 0.00% | 25 | 96.15% | 1 | 3.85% |
| **1.00E-10** | 10.35% | 53 | 17 | 100.00% | 0 | 0.00% | 35 | 97.22% | 1 | 2.78% |
| **1.00E-09** | 12.70% | 65 | 18 | 94.74% | 1 | 5.26% | 45 | 97.83% | 1 | 2.17% |
| **1.00E-08** | 14.26% | 73 | 20 | 95.24% | 1 | 4.76% | 51 | 98.08% | 1 | 1.92% |
| **1.00E-07** | 16.41% | 84 | 21 | 95.45% | 1 | 4.55% | 61 | 98.39% | 1 | 1.61% |
| **1.00E-06** | 19.14% | 98 | 24 | 96.00% | 1 | 4.00% | 72 | 98.63% | 1 | 1.37% |
| **1.00E-05** | 21.48% | 110 | 26 | 96.30% | 1 | 3.70% | 82 | 98.80% | 1 | 1.20% |
| **1.00E-04** | 24.22% | 124 | 33 | 97.06% | 1 | 2.94% | 89 | 98.89% | 1 | 1.11% |
| **1.00E-03** | 28.13% | 144 | 40 | 97.56% | 1 | 2.44% | 102 | 99.03% | 1 | 0.97% |
| **1.00E-02** | 32.03% | 164 | 47 | 95.92% | 2 | 4.08% | 112 | 97.39% | 3 | 2.61% |
| **1.00E-01** | 41.60% | 213 | 62 | 95.38% | 3 | 4.62% | 139 | 93.92% | 9 | 6.08% |
| **1** | 71.09% | 364 | 77 | 92.77% | 6 | 7.23% | 256 | 91.10% | 25 | 8.90% |
| **10** | 99.61% | 510 | 80 | 80.00% | 20 | 20.00% | 370 | 90.24% | 40 | 9.76% |
| **100** | 100.00% | 512 | 79 | 78.22% | 22 | 21.78% | 367 | 89.30% | 44 | 10.70% |

(Note: Chits = correct hits, Whits = wrong/incorrect hits.)

#### *Supplementary Table S4: MERCI result on different ion channel modulating protein dataset.*

#### 1. $Na^+$ independent dataset results

**fp 20 and top 100 motifs**

| **MERCI Options (c=None, fp=20, K=100, fn =0)** | **Collected Motifs** | **Total hits in test data** |
|---|---|---|
| **Positive Motifs** | 74 | 22 |
| **Negative Motifs** | 111 | 397 |

**Test Confusion Matrix:**

| | **Predicted Positive** | **Predicted Negative** |
|---|---|---|
| **Actual Positive** | 15 (TP) | 55 (FN) |
| **Actual Negative** | 7 (FP) | 342 (TN) |

**fp 5 and top 50 motifs Koolman-Rohm**

| **MERCI Options (c=Koolman-Rohm, fp=5, K=50, fn =0)** | Collected Motifs | Total hits in test data |
|---|---|---|
| **Positive Motifs** | 51 | 21 |
| **Negative Motifs** | 51 | 362 |

**Test Confusion Matrix:**

| | Predicted Positive | Predicted Negative |
|---|---|---|
| Actual Positive | 16 (TP) | 44 (FN) |
| Actual Negative | 5 (FP) | 318 (TN) |

#### 2. $K^+$ independent dataset results

**fp 5 and top 50 motifs**

| **MERCI Options (c=None, fp=5, K=50, fn =0)** | Collected Motifs | Total hits in test data |
|---|---|---|
| **Positive Motifs** | 63 | 8 |
| **Negative Motifs** | 58 | 312 |

**Test Confusion Matrix:**

| | Predicted Positive | Predicted Negative |
|---|---|---|
| Actual Positive | 8 (TP) | 44 (FN) |
| Actual Negative | 0 (FP) | 268 (TN) |

```
fp 5 and top 50 motifs Koolman-Rohm
```

| **MERCI Options (c= Koolman-Rohm, fp=5, K=50, fn =0)** | **Collected Motifs** | **Total hits in test data** |
|---|---|---|
| **Positive Motifs** | 65 | 11 |
| **Negative Motifs** | 51 | 376 |

**Test Confusion Matrix:**

| | **Predicted Positive** | **Predicted Negative** |
|---|---|---|
| **Actual Positive** | 8 (TP) | 56 (FN) |
| **Actual Negative** | 3 (FP) | 320 (TN) |

#### 3. $Ca^+$ independent dataset results

```
fp 20 and top 50 motifs
```

| **MERCI Options (c=None, fp=5, K=50, fn =0)** | **Collected Motifs** | **Total hits in test data** |
|---|---|---|
| **Positive Motifs** | 41 | 3 |
| **Negative Motifs** | 62 | 306 |

**Test Confusion Matrix:**

| | Predicted Positive | Predicted Negative |
|---|---|---|
| **Actual Positive** | 2 (TP) | 35 (FN) |
| **Actual Negative** | 1 (FP) | 271 (TN) |

```
fp 5 and top 50 motifs Koolman-Rohm
```

| **MERCI Options (c= Koolman-Rohm, fp=5, K=50, fn =0)** | **Collected Motifs** | **Total hits in test data** |
|---|---|---|
| **Positive Motifs** | 71 | 5 |
| **Negative Motifs** | 52 | 350 |

**Test Confusion Matrix:**

| | Predicted Positive | Predicted Negative |
|---|---|---|
| **Actual Positive** | 1 (TP) | 33 (FN) |

| | | |
|---|---|---|
| **Actual Negative** | 4 (FP) | 317 (TN) |

#### 4. Other independent dataset results

`fp 20 and top 100 motifs`

| **MERCI Options (c=NONE, fp=20, K=100, fn =0)** | Collected Motifs | Total hits in test data |
|---|---|---|
| **Positive Motifs** | 102 | 5 |
| **Negative Motifs** | 108 | 391 |

**Test Confusion Matrix:**

| | Predicted Positive | Predicted Negative |
|---|---|---|
| **Actual Positive** | 2 (TP) | 52 (FN) |
| **Actual Negative** | 3 (FP) | 339 (TN) |

`fp 15 and top 100 motifs Koolman-Rohm`

| **MERCI Options (c= Koolman-Rohm, fp=15, K=500, fn =0)** | Collected Motifs | Total hits in test data |
|---|---|---|
| **Positive Motifs** | 741 | 17 |
| **Negative Motifs** | 551 | 494 |

**Test Confusion Matrix:**

| | Predicted Positive | Predicted Negative |
|---|---|---|
| **Actual Positive** | 5 (TP) | 84 (FN) |
| **Actual Negative** | 12 (FP) | 410 (TN) |

*Supplementary Table S5: Performance of ML-based Classifiers on all four datasets using an array of compositional features extracted using pfeature.*

**Na+ Dataset**

#### Feature Name: AAC

| Classifier | Dataset | SENS | SPEC | PPV | ACC | MCC | AUC |
|---|---|---|---|---|---|---|---|
| **RF** | **Cross-validation** | 0.705 | 0.970 | 0.865 | 0.905 | 0.740 | 0.956 |
| **RF** | **Independent** | 0.570 | 0.966 | 0.839 | 0.872 | 0.620 | 0.925 |
| **GB** | **Cross-validation** | 0.720 | 0.960 | 0.835 | 0.900 | 0.710 | 0.950 |
| **GB** | **Independent** | 0.609 | 0.949 | 0.788 | 0.868 | 0.614 | 0.935 |
| **ElasticNet** | **Cross-validation** | 0.640 | 0.950 | 0.805 | 0.880 | 0.640 | 0.935 |
| **ElasticNet** | **Independent** | 0.586 | 0.956 | 0.806 | 0.868 | 0.611 | 0.925 |
| **Lasso regularization** | **Cross-validation** | 0.630 | 0.951 | 0.800 | 0.875 | 0.630 | 0.934 |
| **Lasso regularization** | **Independent** | 0.578 | 0.956 | 0.804 | 0.866 | 0.604 | 0.925 |
| **Ridge regularization** | **Cross-validation** | 0.650 | 0.955 | 0.820 | 0.890 | 0.655 | 0.938 |
| **Ridge regularization** | **Independent** | 0.594 | 0.964 | 0.835 | 0.876 | 0.633 | 0.926 |
| **LR** | **Cross-validation** | 0.650 | 0.955 | 0.820 | 0.890 | 0.655 | 0.938 |
| **LR** | **Independent** | 0.594 | 0.964 | 0.835 | 0.876 | 0.633 | 0.926 |
| **SVC (rbf)** | **Cross-validation** | 0.710 | 0.965 | 0.860 | 0.900 | 0.715 | 0.960 |
| **SVC (rbf)** | **Independent** | 0.602 | 0.966 | 0.846 | 0.879 | 0.645 | 0.941 |
| **DT** | **Cross-validation** | 0.700 | 0.925 | 0.760 | 0.880 | 0.670 | 0.930 |
| **DT** | **Independent** | 0.672 | 0.915 | 0.711 | 0.857 | 0.598 | 0.899 |
| **MLP** | **Cross-validation** | 0.690 | 0.935 | 0.770 | 0.880 | 0.660 | 0.850 |
| **MLP** | **Independent** | 0.617 | 0.937 | 0.752 | 0.861 | 0.595 | 0.777 |
| **AdB** | **Cross-validation** | 0.690 | 0.960 | 0.850 | 0.890 | 0.660 | 0.950 |
| **AdB** | **Independent** | 0.578 | 0.961 | 0.822 | 0.870 | 0.615 | 0.929 |
| **GNB** | **Cross-validation** | 0.660 | 0.940 | 0.800 | 0.870 | 0.620 | 0.930 |
| **GNB** | **Independent** | 0.570 | 0.934 | 0.730 | 0.848 | 0.552 | 0.886 |
| **XGB** | **Cross-validation** | 0.700 | 0.955 | 0.830 | 0.890 | 0.655 | 0.950 |
| **XGB** | **Independent** | 0.617 | 0.954 | 0.806 | 0.874 | 0.630 | 0.929 |
| **ET** | **Cross-validation** | 0.640 | 0.970 | 0.870 | 0.880 | 0.650 | 0.960 |
| **ET** | **Independent** | 0.547 | 0.978 | 0.886 | 0.876 | 0.632 | 0.949 |

#### Na+ Dataset

#### Feature Name: DPC

| Classifier | Dataset | SENS | SPEC | PPV | ACC | MCC | AUC |
|---|---|---|---|---|---|---|---|
| **RF** | **Cross-validation** | 0.500 | 0.975 | 0.870 | 0.860 | 0.590 | 0.930 |
| **RF** | **Independent** | 0.422 | 0.983 | 0.885 | 0.850 | 0.544 | 0.948 |
| **GB** | **Cross-validation** | 0.540 | 0.960 | 0.790 | 0.860 | 0.580 | 0.920 |
| **GB** | **Independent** | 0.484 | 0.961 | 0.795 | 0.848 | 0.539 | 0.920 |
| **ElasticNet** | **Cross-validation** | 0.540 | 0.970 | 0.860 | 0.870 | 0.610 | 0.930 |
| **ElasticNet** | **Independent** | 0.500 | 0.983 | 0.901 | 0.868 | 0.608 | 0.907 |

| | | | | | | | |
|---|---|---|---|---|---|---|---|
| **Lasso regularization** | **Cross-validation** | 0.530 | 0.970 | 0.860 | 0.870 | 0.600 | 0.930 |
| **Lasso regularization** | **Independent** | 0.477 | 0.983 | 0.897 | 0.863 | 0.589 | 0.896 |
| **Ridge regularization** | **Cross-validation** | 0.550 | 0.975 | 0.870 | 0.875 | 0.620 | 0.935 |
| **Ridge regularization** | **Independent** | 0.508 | 0.981 | 0.890 | 0.868 | 0.607 | 0.906 |
| **LR** | **Cross-validation** | 0.550 | 0.975 | 0.870 | 0.875 | 0.620 | 0.935 |
| **LR** | **Independent** | 0.508 | 0.981 | 0.890 | 0.868 | 0.607 | 0.906 |
| **SVC (rbf)** | **Cross-validation** | 0.520 | 0.975 | 0.870 | 0.860 | 0.590 | 0.935 |
| **SVC (rbf)** | **Independent** | 0.438 | 0.981 | 0.875 | 0.852 | 0.550 | 0.947 |
| **DT** | **Cross-validation** | 0.550 | 0.920 | 0.680 | 0.820 | 0.490 | 0.810 |
| **DT** | **Independent** | 0.508 | 0.908 | 0.631 | 0.813 | 0.450 | 0.708 |
| **MLP** | **Cross-validation** | 0.580 | 0.970 | 0.870 | 0.870 | 0.620 | 0.930 |
| **MLP** | **Independent** | 0.516 | 0.983 | 0.904 | 0.872 | 0.620 | 0.927 |
| **AdB** | **Cross-validation** | 0.530 | 0.960 | 0.820 | 0.850 | 0.560 | 0.920 |
| **AdB** | **Independent** | 0.430 | 0.966 | 0.797 | 0.839 | 0.504 | 0.857 |
| **GNB** | **Cross-validation** | 0.530 | 0.960 | 0.810 | 0.850 | 0.550 | 0.920 |
| **GNB** | **Independent** | 0.430 | 0.961 | 0.775 | 0.835 | 0.492 | 0.920 |
| **XGB** | **Cross-validation** | 0.460 | 0.985 | 0.910 | 0.860 | 0.580 | 0.940 |
| **XGB** | **Independent** | 0.375 | 0.993 | 0.941 | 0.846 | 0.535 | 0.952 |
| **ET** | **Cross-validation** | 0.500 | 0.975 | 0.870 | 0.860 | 0.590 | 0.930 |
| **ET** | **Independent** | 0.422 | 0.983 | 0.885 | 0.850 | 0.544 | 0.948 |

(Abbreviations: SENS: Sensitivity; SPEC: Specificity; PPV: Positive Predictive Value; ACC: Accuracy; MCC: Matthews Correlation Coefficient; AUC: Area Under the Receiver Operating Characteristic Curve)

#### Na+ Dataset

#### Feature Name: AAC+DPC

| Classifier | Dataset | SENS | SPEC | PPV | ACC | MCC | AUC |
|---|---|---|---|---|---|---|---|
| **RF** | **Cross-validation** | 0.660 | 0.975 | 0.880 | 0.870 | 0.600 | 0.940 |
| **RF** | **Independent** | 0.577 | 0.978 | 0.871 | 0.859 | 0.576 | 0.955 |
| **GB** | **Cross-validation** | 0.620 | 0.960 | 0.830 | 0.875 | 0.610 | 0.950 |
| **GB** | **Independent** | 0.570 | 0.964 | 0.830 | 0.870 | 0.615 | 0.948 |
| **ElasticNet** | **Cross-validation** | 0.580 | 0.970 | 0.850 | 0.875 | 0.600 | 0.910 |
| **ElasticNet** | **Independent** | 0.500 | 0.976 | 0.865 | 0.863 | 0.588 | 0.910 |
| **Lasso regularization** | **Cross-validation** | 0.570 | 0.970 | 0.850 | 0.875 | 0.590 | 0.910 |
| **Lasso regularization** | **Independent** | 0.484 | 0.978 | 0.873 | 0.861 | 0.582 | 0.912 |
| **Ridge regularization** | **Cross-validation** | 0.590 | 0.970 | 0.860 | 0.880 | 0.610 | 0.915 |
| **Ridge regularization** | **Independent** | 0.508 | 0.976 | 0.867 | 0.865 | 0.594 | 0.914 |
| **LR** | **Cross-validation** | 0.590 | 0.970 | 0.860 | 0.880 | 0.610 | 0.915 |
| **LR** | **Independent** | 0.508 | 0.976 | 0.867 | 0.865 | 0.594 | 0.915 |
| **SVC (rbf)** | **Cross-validation** | 0.630 | 0.970 | 0.850 | 0.885 | 0.620 | 0.945 |
| **SVC (rbf)** | **Independent** | 0.578 | 0.966 | 0.841 | 0.874 | 0.626 | 0.944 |

| | | | | | | | |
|---|---|---|---|---|---|---|---|
| **DT** | **Cross-validation** | 0.650 | 0.935 | 0.740 | 0.855 | 0.580 | 0.860 |
| **DT** | **Independent** | 0.617 | 0.922 | 0.712 | 0.850 | 0.568 | 0.865 |
| **MLP** | **Cross-validation** | 0.640 | 0.925 | 0.720 | 0.850 | 0.570 | 0.830 |
| **MLP** | **Independent** | 0.602 | 0.915 | 0.688 | 0.840 | 0.542 | 0.758 |
| **AdB** | **Cross-validation** | 0.610 | 0.975 | 0.900 | 0.880 | 0.610 | 0.930 |
| **AdB** | **Independent** | 0.516 | 0.988 | 0.930 | 0.876 | 0.633 | 0.927 |
| **GNB** | **Cross-validation** | 0.580 | 0.960 | 0.850 | 0.870 | 0.600 | 0.900 |
| **GNB** | **Independent** | 0.500 | 0.971 | 0.842 | 0.859 | 0.576 | 0.898 |
| **XGB** | **Cross-validation** | 0.630 | 0.970 | 0.850 | 0.885 | 0.620 | 0.945 |
| **XGB** | **Independent** | 0.578 | 0.961 | 0.822 | 0.870 | 0.615 | 0.941 |
| **ET** | **Cross-validation** | 0.490 | 0.985 | 0.930 | 0.875 | 0.580 | 0.950 |
| **ET** | **Independent** | 0.414 | 0.990 | 0.930 | 0.853 | 0.560 | 0.947 |

(Abbreviations: SENS: Sensitivity; SPEC: Specificity; PPV: Positive Predictive Value; ACC: Accuracy; MCC: Matthews Correlation Coefficient; AUC: Area Under the Receiver Operating Characteristic Curve)

#### Na+ Dataset

#### Feature Name: ALLCOMP

| Classifier | Dataset | SENS | SPEC | PPV | ACC | MCC | AUC |
|---|---|---|---|---|---|---|---|
| **RF** | **Cross-validation** | 0.723 | 0.973 | 0.892 | 0.921 | 0.702 | 0.914 |
| **RF** | **Independent** | 0.516 | 0.973 | 0.857 | 0.865 | 0.594 | 0.959 |
| **GB** | **Cross-validation** | 0.746 | 0.974 | 0.901 | 0.927 | 0.724 | 0.917 |
| **GB** | **Independent** | 0.570 | 0.976 | 0.880 | 0.879 | 0.644 | 0.957 |
| **ElasticNet** | **Cross-validation** | 0.762 | 0.966 | 0.876 | 0.922 | 0.734 | 0.911 |
| **ElasticNet** | **Independent** | 0.602 | 0.966 | 0.846 | 0.879 | 0.645 | 0.957 |
| **Lasso regularization** | **Cross-validation** | 0.664 | 0.960 | 0.840 | 0.885 | 0.665 | 0.905 |
| **Lasso regularization** | **Independent** | 0.516 | 0.961 | 0.805 | 0.855 | 0.565 | 0.895 |
| **Ridge regularization** | **Cross-validation** | 0.658 | 0.961 | 0.840 | 0.883 | 0.659 | 0.900 |
| **Ridge regularization** | **Independent** | 0.508 | 0.961 | 0.802 | 0.853 | 0.558 | 0.894 |
| **LR** | **Cross-validation** | 0.674 | 0.960 | 0.839 | 0.888 | 0.674 | 0.902 |
| **LR** | **Independent** | 0.531 | 0.959 | 0.800 | 0.857 | 0.572 | 0.896 |
| **SVC (rbf)** | **Cross-validation** | 0.674 | 0.960 | 0.839 | 0.888 | 0.674 | 0.902 |
| **SVC (rbf)** | **Independent** | 0.531 | 0.959 | 0.800 | 0.857 | 0.572 | 0.896 |
| **DT** | **Cross-validation** | 0.693 | 0.951 | 0.816 | 0.884 | 0.678 | 0.892 |
| **DT** | **Independent** | 0.563 | 0.934 | 0.727 | 0.846 | 0.546 | 0.882 |
| **MLP** | **Cross-validation** | 0.820 | 0.960 | 0.866 | 0.912 | 0.761 | 0.900 |
| **MLP** | **Independent** | 0.672 | 0.951 | 0.811 | 0.885 | 0.667 | 0.907 |
| **AdB** | **Cross-validation** | 0.898 | 0.948 | 0.844 | 0.926 | 0.812 | 0.902 |
| **AdB** | **Independent** | 0.570 | 0.961 | 0.820 | 0.868 | 0.609 | 0.766 |

| | | | | | | | |
|---|---|---|---|---|---|---|---|
| GNB | Cross-validation | 0.742 | 0.978 | 0.914 | 0.919 | 0.725 | 0.925 |
| GNB | Independent | 0.516 | 0.978 | 0.880 | 0.868 | 0.607 | 0.933 |
| XGB | Cross-validation | 0.791 | 0.972 | 0.900 | 0.911 | 0.739 | 0.919 |
| XGB | Independent | 0.625 | 0.968 | 0.860 | 0.887 | 0.668 | 0.933 |
| ET | Cross-validation | 0.801 | 0.979 | 0.921 | 0.922 | 0.747 | 0.931 |
| ET | Independent | 0.594 | 0.976 | 0.884 | 0.885 | 0.662 | 0.959 |

(Abbreviations: SENS: Sensitivity; SPEC: Specificity; PPV: Positive Predictive Value; ACC: Accuracy; MCC: Matthews Correlation Coefficient; AUC: Area Under the Receiver Operating Characteristic Curve)

#### K+ Dataset

#### Feature Name: AAC

| Classifier | Dataset | SENS | SPEC | PPV | ACC | MCC | AUC |
|---|---|---|---|---|---|---|---|
| RF | Cross-validation | 0.640 | 0.960 | 0.800 | 0.890 | 0.660 | 0.940 |
| RF | Independent | 0.550 | 0.959 | 0.779 | 0.873 | 0.583 | 0.907 |
| GB | Cross-validation | 0.670 | 0.940 | 0.750 | 0.880 | 0.640 | 0.940 |
| GB | Independent | 0.578 | 0.942 | 0.724 | 0.865 | 0.567 | 0.903 |
| ElasticNet | Cross-validation | 0.520 | 0.950 | 0.740 | 0.860 | 0.520 | 0.910 |
| ElasticNet | Independent | 0.440 | 0.951 | 0.706 | 0.844 | 0.473 | 0.891 |
| Lasso regularization | Cross-validation | 0.520 | 0.950 | 0.740 | 0.860 | 0.520 | 0.910 |
| Lasso regularization | Independent | 0.440 | 0.951 | 0.706 | 0.844 | 0.473 | 0.891 |
| Ridge regularization | Cross-validation | 0.510 | 0.945 | 0.720 | 0.850 | 0.500 | 0.900 |
| Ridge regularization | Independent | 0.431 | 0.946 | 0.681 | 0.838 | 0.453 | 0.887 |
| LR | Cross-validation | 0.510 | 0.945 | 0.720 | 0.850 | 0.500 | 0.900 |
| LR | Independent | 0.431 | 0.946 | 0.681 | 0.838 | 0.453 | 0.887 |
| SVC (rbf) | Cross-validation | 0.640 | 0.940 | 0.740 | 0.870 | 0.620 | 0.930 |
| SVC (rbf) | Independent | 0.560 | 0.934 | 0.693 | 0.856 | 0.536 | 0.902 |
| DT | Cross-validation | 0.610 | 0.920 | 0.670 | 0.860 | 0.530 | 0.910 |
| DT | Independent | 0.578 | 0.908 | 0.624 | 0.838 | 0.500 | 0.876 |
| MLP | Cross-validation | 0.600 | 0.910 | 0.630 | 0.850 | 0.500 | 0.770 |
| MLP | Independent | 0.495 | 0.900 | 0.568 | 0.815 | 0.417 | 0.698 |
| AdB | Cross-validation | 0.660 | 0.940 | 0.760 | 0.880 | 0.640 | 0.930 |
| AdB | Independent | 0.569 | 0.954 | 0.765 | 0.873 | 0.587 | 0.900 |
| GNB | Cross-validation | 0.660 | 0.930 | 0.720 | 0.870 | 0.590 | 0.910 |
| GNB | Independent | 0.606 | 0.934 | 0.710 | 0.865 | 0.573 | 0.888 |
| XGB | Cross-validation | 0.670 | 0.940 | 0.740 | 0.880 | 0.630 | 0.930 |
| XGB | Independent | 0.596 | 0.946 | 0.747 | 0.873 | 0.592 | 0.913 |
| ET | Cross-validation | 0.530 | 0.960 | 0.770 | 0.850 | 0.500 | 0.910 |
| ET | Independent | 0.431 | 0.966 | 0.770 | 0.854 | 0.502 | 0.920 |

(Abbreviations: SENS: Sensitivity; SPEC: Specificity; PPV: Positive Predictive Value; ACC: Accuracy; MCC: Matthews Correlation Coefficient; AUC: Area Under the Receiver Operating Characteristic Curve)

### K+ Dataset

#### Feature Name: DPC

| Classifier | Dataset | SENS | SPEC | PPV | ACC | MCC | AUC |
|---|---|---|---|---|---|---|---|
| **RF** | **Cross-validation** | 0.470 | 0.970 | 0.840 | 0.870 | 0.540 | 0.910 |
| **RF** | **Independent** | 0.367 | 0.976 | 0.800 | 0.848 | 0.473 | 0.901 |
| **GB** | **Cross-validation** | 0.530 | 0.960 | 0.790 | 0.870 | 0.570 | 0.920 |
| **GB** | **Independent** | 0.495 | 0.954 | 0.740 | 0.858 | 0.526 | 0.899 |
| **ElasticNet** | **Cross-validation** | 0.520 | 0.960 | 0.800 | 0.870 | 0.560 | 0.910 |
| **ElasticNet** | **Independent** | 0.468 | 0.961 | 0.761 | 0.858 | 0.521 | 0.896 |
| **Lasso regularization** | **Cross-validation** | 0.510 | 0.960 | 0.790 | 0.860 | 0.550 | 0.910 |
| **Lasso regularization** | **Independent** | 0.486 | 0.964 | 0.779 | 0.863 | 0.543 | 0.892 |
| **Ridge regularization** | **Cross-validation** | 0.520 | 0.960 | 0.800 | 0.870 | 0.560 | 0.910 |
| **Ridge regularization** | **Independent** | 0.468 | 0.959 | 0.750 | 0.856 | 0.515 | 0.894 |
| **LR** | **Cross-validation** | 0.520 | 0.960 | 0.800 | 0.870 | 0.560 | 0.910 |
| **LR** | **Independent** | 0.468 | 0.959 | 0.750 | 0.856 | 0.515 | 0.894 |
| **SVC (rbf)** | **Cross-validation** | 0.510 | 0.955 | 0.780 | 0.860 | 0.530 | 0.910 |
| **SVC (rbf)** | **Independent** | 0.468 | 0.956 | 0.739 | 0.854 | 0.509 | 0.900 |
| **DT** | **Cross-validation** | 0.540 | 0.910 | 0.670 | 0.830 | 0.470 | 0.810 |
| **DT** | **Independent** | 0.505 | 0.895 | 0.561 | 0.813 | 0.416 | 0.700 |
| **MLP** | **Cross-validation** | 0.520 | 0.950 | 0.750 | 0.860 | 0.530 | 0.900 |
| **MLP** | **Independent** | 0.505 | 0.951 | 0.733 | 0.858 | 0.528 | 0.900 |
| **AdB** | **Cross-validation** | 0.510 | 0.950 | 0.750 | 0.850 | 0.520 | 0.900 |
| **AdB** | **Independent** | 0.495 | 0.946 | 0.711 | 0.852 | 0.509 | 0.848 |
| **GNB** | **Cross-validation** | 0.510 | 0.960 | 0.780 | 0.860 | 0.530 | 0.910 |
| **GNB** | **Independent** | 0.450 | 0.956 | 0.731 | 0.850 | 0.493 | 0.897 |
| **XGB** | **Cross-validation** | 0.400 | 0.990 | 0.880 | 0.850 | 0.520 | 0.930 |
| **XGB** | **Independent** | 0.294 | 0.983 | 0.821 | 0.838 | 0.427 | 0.911 |
| **ET** | **Cross-validation** | 0.470 | 0.970 | 0.840 | 0.870 | 0.540 | 0.910 |
| **ET** | **Independent** | 0.367 | 0.976 | 0.800 | 0.848 | 0.473 | 0.901 |

(Abbreviations: SENS: Sensitivity; SPEC: Specificity; PPV: Positive Predictive Value; ACC: Accuracy; MCC: Matthews Correlation Coefficient; AUC: Area Under the Receiver Operating Characteristic Curve)

### K+ Dataset

##### Feature Name: AAC+DPC

| Classifier | Dataset | SENS | SPEC | PPV | ACC | MCC | AUC |
|---|---|---|---|---|---|---|---|
| **RF** | **Cross-validation** | 0.600 | 0.970 | 0.850 | 0.900 | 0.670 | 0.940 |
| **RF** | **Independent** | 0.450 | 0.966 | 0.778 | 0.858 | 0.518 | 0.915 |
| **GB** | **Cross-validation** | 0.660 | 0.950 | 0.800 | 0.890 | 0.690 | 0.940 |
| **GB** | **Independent** | 0.578 | 0.949 | 0.750 | 0.871 | 0.583 | 0.916 |
| **ElasticNet** | **Cross-validation** | 0.630 | 0.950 | 0.800 | 0.880 | 0.650 | 0.910 |
| **ElasticNet** | **Independent** | 0.495 | 0.959 | 0.761 | 0.862 | 0.538 | 0.889 |
| **Lasso regularization** | **Cross-validation** | 0.630 | 0.955 | 0.810 | 0.885 | 0.655 | 0.910 |
| **Lasso regularization** | **Independent** | 0.532 | 0.961 | 0.784 | 0.871 | 0.575 | 0.891 |
| **Ridge regularization** | **Cross-validation** | 0.620 | 0.960 | 0.820 | 0.885 | 0.660 | 0.920 |
| **Ridge regularization** | **Independent** | 0.477 | 0.956 | 0.743 | 0.856 | 0.517 | 0.892 |
| **LR** | **Cross-validation** | 0.620 | 0.960 | 0.820 | 0.885 | 0.660 | 0.920 |
| **LR** | **Independent** | 0.477 | 0.956 | 0.743 | 0.856 | 0.517 | 0.892 |
| **SVC (rbf)** | **Cross-validation** | 0.700 | 0.955 | 0.810 | 0.890 | 0.690 | 0.940 |
| **SVC (rbf)** | **Independent** | 0.596 | 0.944 | 0.739 | 0.871 | 0.587 | 0.907 |
| **DT** | **Cross-validation** | 0.670 | 0.920 | 0.720 | 0.865 | 0.610 | 0.900 |
| **DT** | **Independent** | 0.624 | 0.908 | 0.642 | 0.848 | 0.537 | 0.865 |
| **MLP** | **Cross-validation** | 0.630 | 0.925 | 0.700 | 0.860 | 0.590 | 0.800 |
| **MLP** | **Independent** | 0.486 | 0.929 | 0.646 | 0.837 | 0.464 | 0.708 |
| **AdB** | **Cross-validation** | 0.680 | 0.955 | 0.800 | 0.890 | 0.670 | 0.930 |
| **AdB** | **Independent** | 0.514 | 0.961 | 0.778 | 0.867 | 0.560 | 0.902 |
| **GNB** | **Cross-validation** | 0.650 | 0.945 | 0.790 | 0.880 | 0.640 | 0.910 |
| **GNB** | **Independent** | 0.560 | 0.951 | 0.753 | 0.869 | 0.574 | 0.890 |
| **XGB** | **Cross-validation** | 0.670 | 0.950 | 0.800 | 0.885 | 0.670 | 0.930 |
| **XGB** | **Independent** | 0.550 | 0.951 | 0.750 | 0.867 | 0.566 | 0.909 |
| **ET** | **Cross-validation** | 0.500 | 0.985 | 0.910 | 0.860 | 0.580 | 0.940 |
| **ET** | **Independent** | 0.303 | 0.985 | 0.846 | 0.842 | 0.445 | 0.917 |

(Abbreviations: SENS: Sensitivity; SPEC: Specificity; PPV: Positive Predictive Value; ACC: Accuracy; MCC: Matthews Correlation Coefficient; AUC: Area Under the Receiver Operating Characteristic Curve)

#### K+ Dataset

##### Feature Name: ALLCOMP

| Classifier | Dataset | SENS | SPEC | PPV | ACC | MCC | AUC |
|---|---|---|---|---|---|---|---|
| **RF** | **Cross-validation** | 0.660 | 0.975 | 0.880 | 0.890 | 0.700 | 0.935 |
| **RF** | **Independent** | 0.486 | 0.981 | 0.869 | 0.877 | 0.590 | 0.940 |

| | | | | | | | |
|---|---|---|---|---|---|---|---|
| **GB** | **Cross-validation** | 0.730 | 0.955 | 0.830 | 0.890 | 0.700 | 0.930 |
| **GB** | **Independent** | 0.587 | 0.961 | 0.800 | 0.883 | 0.618 | 0.923 |
| **ElasticNet** | **Cross-validation** | 0.380 | 0.970 | 0.800 | 0.840 | 0.490 | 0.880 |
| **ElasticNet** | **Independent** | 0.303 | 0.968 | 0.717 | 0.829 | 0.389 | 0.860 |
| **Lasso regularization** | **Cross-validation** | 0.390 | 0.970 | 0.800 | 0.842 | 0.495 | 0.885 |
| **Lasso regularization** | **Independent** | 0.303 | 0.968 | 0.717 | 0.829 | 0.389 | 0.859 |
| **Ridge regularization** | **Cross-validation** | 0.400 | 0.968 | 0.790 | 0.840 | 0.490 | 0.880 |
| **Ridge regularization** | **Independent** | 0.303 | 0.966 | 0.702 | 0.827 | 0.381 | 0.864 |
| **LR** | **Cross-validation** | 0.400 | 0.968 | 0.790 | 0.840 | 0.490 | 0.880 |
| **LR** | **Independent** | 0.303 | 0.966 | 0.702 | 0.827 | 0.381 | 0.864 |
| **SVC (rbf)** | **Cross-validation** | 0.710 | 0.950 | 0.800 | 0.880 | 0.700 | 0.920 |
| **SVC (rbf)** | **Independent** | 0.477 | 0.959 | 0.754 | 0.858 | 0.523 | 0.908 |
| **DT** | **Cross-validation** | 0.430 | 0.960 | 0.740 | 0.850 | 0.510 | 0.870 |
| **DT** | **Independent** | 0.303 | 0.961 | 0.673 | 0.823 | 0.368 | 0.855 |
| **MLP** | **Cross-validation** | 0.760 | 0.925 | 0.750 | 0.880 | 0.680 | 0.890 |
| **MLP** | **Independent** | 0.569 | 0.939 | 0.713 | 0.862 | 0.554 | 0.878 |
| **AdB** | **Cross-validation** | 0.670 | 0.930 | 0.750 | 0.870 | 0.630 | 0.820 |
| **AdB** | **Independent** | 0.560 | 0.932 | 0.685 | 0.854 | 0.531 | 0.746 |
| **GNB** | **Cross-validation** | 0.720 | 0.960 | 0.800 | 0.880 | 0.690 | 0.920 |
| **GNB** | **Independent** | 0.422 | 0.968 | 0.780 | 0.854 | 0.501 | 0.927 |
| **XGB** | **Cross-validation** | 0.720 | 0.960 | 0.800 | 0.880 | 0.690 | 0.920 |
| **XGB** | **Independent** | 0.606 | 0.959 | 0.795 | 0.885 | 0.627 | 0.924 |
| **ET** | **Cross-validation** | 0.760 | 0.960 | 0.810 | 0.890 | 0.720 | 0.930 |
| **ET** | **Independent** | 0.615 | 0.954 | 0.779 | 0.883 | 0.623 | 0.942 |

(Abbreviations: SENS: Sensitivity; SPEC: Specificity; PPV: Positive Predictive Value; ACC: Accuracy; MCC: Matthews Correlation Coefficient; AUC: Area Under the Receiver Operating Characteristic Curve)

#### Ca+ Dataset

#### Feature Name: AAC

| Classifier | Dataset | SENS | SPEC | PPV | ACC | MCC | AUC |
|---|---|---|---|---|---|---|---|
| **RF** | **Cross-validation** | 0.600 | 0.975 | 0.820 | 0.910 | 0.640 | 0.950 |
| **RF** | **Independent** | 0.560 | 0.971 | 0.778 | 0.907 | 0.610 | 0.939 |
| **GB** | **Cross-validation** | 0.640 | 0.960 | 0.760 | 0.900 | 0.640 | 0.940 |
| **GB** | **Independent** | 0.600 | 0.956 | 0.714 | 0.901 | 0.598 | 0.910 |
| **ElasticNet** | **Cross-validation** | 0.460 | 0.965 | 0.730 | 0.880 | 0.500 | 0.910 |
| **ElasticNet** | **Independent** | 0.467 | 0.963 | 0.700 | 0.887 | 0.511 | 0.895 |

| | | | | | | | |
|---|---|---|---|---|---|---|---|
| **Lasso regularization** | **Cross-validation** | 0.460 | 0.965 | 0.730 | 0.880 | 0.500 | 0.910 |
| **Lasso regularization** | **Independent** | 0.467 | 0.963 | 0.700 | 0.887 | 0.511 | 0.894 |
| **Ridge regularization** | **Cross-validation** | 0.490 | 0.965 | 0.720 | 0.880 | 0.520 | 0.910 |
| **Ridge regularization** | **Independent** | 0.507 | 0.959 | 0.691 | 0.889 | 0.530 | 0.896 |
| **LR** | **Cross-validation** | 0.490 | 0.965 | 0.720 | 0.880 | 0.520 | 0.910 |
| **LR** | **Independent** | 0.507 | 0.959 | 0.691 | 0.889 | 0.530 | 0.896 |
| **SVC (rbf)** | **Cross-validation** | 0.630 | 0.960 | 0.770 | 0.900 | 0.620 | 0.940 |
| **SVC (rbf)** | **Independent** | 0.507 | 0.966 | 0.731 | 0.895 | 0.552 | 0.911 |
| **DT** | **Cross-validation** | 0.610 | 0.920 | 0.630 | 0.870 | 0.550 | 0.900 |
| **DT** | **Independent** | 0.613 | 0.915 | 0.568 | 0.868 | 0.512 | 0.868 |
| **MLP** | **Cross-validation** | 0.600 | 0.930 | 0.650 | 0.870 | 0.560 | 0.780 |
| **MLP** | **Independent** | 0.560 | 0.944 | 0.646 | 0.885 | 0.535 | 0.752 |
| **AdB** | **Cross-validation** | 0.600 | 0.970 | 0.800 | 0.900 | 0.610 | 0.910 |
| **AdB** | **Independent** | 0.467 | 0.983 | 0.833 | 0.903 | 0.578 | 0.880 |
| **GNB** | **Cross-validation** | 0.630 | 0.950 | 0.700 | 0.880 | 0.600 | 0.920 |
| **GNB** | **Independent** | 0.573 | 0.959 | 0.717 | 0.899 | 0.584 | 0.914 |
| **XGB** | **Cross-validation** | 0.660 | 0.960 | 0.750 | 0.900 | 0.640 | 0.940 |
| **XGB** | **Independent** | 0.627 | 0.961 | 0.746 | 0.909 | 0.632 | 0.927 |
| **ET** | **Cross-validation** | 0.550 | 0.980 | 0.850 | 0.890 | 0.600 | 0.950 |
| **ET** | **Independent** | 0.453 | 0.985 | 0.850 | 0.903 | 0.577 | 0.934 |

(Abbreviations: SENS: Sensitivity; SPEC: Specificity; PPV: Positive Predictive Value; ACC: Accuracy; MCC: Matthews Correlation Coefficient; AUC: Area Under the Receiver Operating Characteristic Curve)

#### Ca+ Dataset

#### Feature Name: DPC

| Classifier | Dataset | SENS | SPEC | PPV | ACC | MCC | AUC |
|---|---|---|---|---|---|---|---|
| **RF** | **Cross-validation** | 0.420 | 0.980 | 0.860 | 0.880 | 0.540 | 0.920 |
| **RF** | **Independent** | 0.267 | 0.990 | 0.833 | 0.878 | 0.428 | 0.919 |
| **GB** | **Cross-validation** | 0.470 | 0.970 | 0.800 | 0.880 | 0.530 | 0.910 |
| **GB** | **Independent** | 0.360 | 0.973 | 0.711 | 0.878 | 0.448 | 0.888 |
| **ElasticNet** | **Cross-validation** | 0.480 | 0.960 | 0.710 | 0.870 | 0.520 | 0.880 |
| **ElasticNet** | **Independent** | 0.373 | 0.966 | 0.667 | 0.874 | 0.436 | 0.842 |
| **Lasso regularization** | **Cross-validation** | 0.480 | 0.950 | 0.700 | 0.860 | 0.510 | 0.860 |
| **Lasso regularization** | **Independent** | 0.373 | 0.966 | 0.667 | 0.874 | 0.436 | 0.833 |
| **Ridge regularization** | **Cross-validation** | 0.460 | 0.960 | 0.710 | 0.860 | 0.500 | 0.880 |
| **Ridge regularization** | **Independent** | 0.347 | 0.971 | 0.684 | 0.874 | 0.427 | 0.867 |
| **LR** | **Cross-validation** | 0.460 | 0.960 | 0.710 | 0.860 | 0.500 | 0.880 |
| **LR** | **Independent** | 0.347 | 0.971 | 0.684 | 0.874 | 0.427 | 0.867 |

| | | | | | | | |
|---|---|---|---|---|---|---|---|
| **SVC (rbf)** | **Cross-validation** | 0.410 | 0.980 | 0.790 | 0.870 | 0.500 | 0.900 |
| **SVC (rbf)** | **Independent** | 0.267 | 0.985 | 0.769 | 0.874 | 0.405 | 0.908 |
| **DT** | **Cross-validation** | 0.520 | 0.930 | 0.620 | 0.860 | 0.470 | 0.780 |
| **DT** | **Independent** | 0.520 | 0.929 | 0.574 | 0.866 | 0.468 | 0.725 |
| **MLP** | **Cross-validation** | 0.500 | 0.970 | 0.740 | 0.870 | 0.510 | 0.880 |
| **MLP** | **Independent** | 0.400 | 0.968 | 0.698 | 0.880 | 0.468 | 0.875 |
| **AdB** | **Cross-validation** | 0.480 | 0.960 | 0.720 | 0.860 | 0.480 | 0.880 |
| **AdB** | **Independent** | 0.427 | 0.961 | 0.667 | 0.878 | 0.469 | 0.833 |
| **GNB** | **Cross-validation** | 0.500 | 0.970 | 0.770 | 0.880 | 0.520 | 0.890 |
| **GNB** | **Independent** | 0.400 | 0.978 | 0.769 | 0.889 | 0.503 | 0.890 |
| **XGB** | **Cross-validation** | 0.390 | 0.990 | 0.860 | 0.870 | 0.480 | 0.920 |
| **XGB** | **Independent** | 0.240 | 0.993 | 0.857 | 0.876 | 0.413 | 0.914 |
| **ET** | **Cross-validation** | 0.420 | 0.980 | 0.860 | 0.880 | 0.540 | 0.920 |
| **ET** | **Independent** | 0.267 | 0.990 | 0.833 | 0.878 | 0.428 | 0.919 |

(Abbreviations: SENS: Sensitivity; SPEC: Specificity; PPV: Positive Predictive Value; ACC: Accuracy; MCC: Matthews Correlation Coefficient; AUC: Area Under the Receiver Operating Characteristic Curve)

#### Ca+ Dataset

#### Feature Name: AAC+DPC

| Classifier | Dataset | SENS | SPEC | PPV | ACC | MCC | AUC |
|---|---|---|---|---|---|---|---|
| **RF** | **Cross-validation** | 0.580 | 0.970 | 0.850 | 0.900 | 0.660 | 0.940 |
| **RF** | **Independent** | 0.280 | 0.990 | 0.840 | 0.880 | 0.442 | 0.936 |
| **GB** | **Cross-validation** | 0.650 | 0.960 | 0.800 | 0.900 | 0.680 | 0.930 |
| **GB** | **Independent** | 0.480 | 0.971 | 0.750 | 0.895 | 0.546 | 0.908 |
| **ElasticNet** | **Cross-validation** | 0.610 | 0.950 | 0.760 | 0.880 | 0.630 | 0.900 |
| **ElasticNet** | **Independent** | 0.440 | 0.966 | 0.702 | 0.885 | 0.496 | 0.887 |
| **Lasso regularization** | **Cross-validation** | 0.610 | 0.945 | 0.750 | 0.875 | 0.615 | 0.890 |
| **Lasso regularization** | **Independent** | 0.427 | 0.966 | 0.696 | 0.882 | 0.484 | 0.865 |
| **Ridge regularization** | **Cross-validation** | 0.600 | 0.955 | 0.770 | 0.880 | 0.620 | 0.900 |
| **Ridge regularization** | **Independent** | 0.413 | 0.963 | 0.674 | 0.878 | 0.465 | 0.880 |
| **LR** | **Cross-validation** | 0.600 | 0.955 | 0.770 | 0.880 | 0.620 | 0.900 |
| **LR** | **Independent** | 0.413 | 0.963 | 0.674 | 0.878 | 0.465 | 0.879 |
| **SVC (rbf)** | **Cross-validation** | 0.630 | 0.960 | 0.790 | 0.890 | 0.640 | 0.910 |
| **SVC (rbf)** | **Independent** | 0.413 | 0.968 | 0.705 | 0.882 | 0.480 | 0.901 |
| **DT** | **Cross-validation** | 0.670 | 0.920 | 0.700 | 0.860 | 0.590 | 0.870 |
| **DT** | **Independent** | 0.667 | 0.939 | 0.667 | 0.897 | 0.606 | 0.802 |
| **MLP** | **Cross-validation** | 0.640 | 0.920 | 0.690 | 0.860 | 0.590 | 0.800 |
| **MLP** | **Independent** | 0.413 | 0.951 | 0.608 | 0.868 | 0.430 | 0.682 |
| **AdB** | **Cross-validation** | 0.690 | 0.960 | 0.800 | 0.900 | 0.670 | 0.920 |
| **AdB** | **Independent** | 0.320 | 0.971 | 0.667 | 0.870 | 0.401 | 0.876 |

| | | | | | | | |
|---|---|---|---|---|---|---|---|
| **GNB** | **Cross-validation** | 0.620 | 0.960 | 0.770 | 0.880 | 0.610 | 0.900 |
| **GNB** | **Independent** | 0.507 | 0.939 | 0.603 | 0.872 | 0.479 | 0.890 |
| **XGB** | **Cross-validation** | 0.650 | 0.965 | 0.810 | 0.890 | 0.670 | 0.920 |
| **XGB** | **Independent** | 0.493 | 0.966 | 0.725 | 0.893 | 0.541 | 0.917 |
| **ET** | **Cross-validation** | 0.480 | 0.990 | 0.920 | 0.860 | 0.560 | 0.930 |
| **ET** | **Independent** | 0.253 | 0.990 | 0.826 | 0.876 | 0.414 | 0.919 |

(Abbreviations: SENS: Sensitivity; SPEC: Specificity; PPV: Positive Predictive Value; ACC: Accuracy; MCC: Matthews Correlation Coefficient; AUC: Area Under the Receiver Operating Characteristic Curve)

#### Ca+ Dataset

#### Feature Name: ALLCOMP

| Classifier | Dataset | SENS | SPEC | PPV | ACC | MCC | AUC |
|---|---|---|---|---|---|---|---|
| **RF** | **Cross-validation** | 0.590 | 0.985 | 0.910 | 0.900 | 0.680 | 0.940 |
| **RF** | **Independent** | 0.473 | 0.988 | 0.848 | 0.893 | 0.519 | 0.943 |
| **GB** | **Cross-validation** | 0.670 | 0.975 | 0.890 | 0.910 | 0.720 | 0.930 |
| **GB** | **Independent** | 0.427 | 0.976 | 0.762 | 0.891 | 0.517 | 0.935 |
| **ElasticNet** | **Cross-validation** | 0.250 | 0.985 | 0.840 | 0.870 | 0.430 | 0.860 |
| **ElasticNet** | **Independent** | 0.280 | 0.980 | 0.724 | 0.872 | 0.397 | 0.886 |
| **Lasso regularization** | **Cross-validation** | 0.250 | 0.985 | 0.835 | 0.870 | 0.420 | 0.860 |
| **Lasso regularization** | **Independent** | 0.280 | 0.983 | 0.750 | 0.874 | 0.408 | 0.886 |
| **Ridge regularization** | **Cross-validation** | 0.270 | 0.983 | 0.830 | 0.870 | 0.440 | 0.860 |
| **Ridge regularization** | **Independent** | 0.280 | 0.980 | 0.724 | 0.872 | 0.397 | 0.887 |
| **LR** | **Cross-validation** | 0.270 | 0.983 | 0.830 | 0.870 | 0.440 | 0.860 |
| **LR** | **Independent** | 0.280 | 0.980 | 0.724 | 0.872 | 0.397 | 0.887 |
| **SVC (rbf)** | **Cross-validation** | 0.680 | 0.960 | 0.830 | 0.910 | 0.700 | 0.910 |
| **SVC (rbf)** | **Independent** | 0.413 | 0.973 | 0.738 | 0.887 | 0.497 | 0.885 |
| **GNB** | **Cross-validation** | 0.230 | 0.980 | 0.800 | 0.860 | 0.390 | 0.830 |
| **GNB** | **Independent** | 0.213 | 0.990 | 0.800 | 0.870 | 0.370 | 0.855 |
| **DT** | **Cross-validation** | 0.700 | 0.940 | 0.720 | 0.910 | 0.660 | 0.880 |
| **DT** | **Independent** | 0.347 | 0.956 | 0.591 | 0.862 | 0.381 | 0.742 |
| **MLP** | **Cross-validation** | 0.670 | 0.940 | 0.720 | 0.900 | 0.660 | 0.820 |
| **MLP** | **Independent** | 0.533 | 0.944 | 0.635 | 0.880 | 0.513 | 0.739 |
| **AdB** | **Cross-validation** | 0.720 | 0.960 | 0.820 | 0.910 | 0.690 | 0.920 |
| **AdB** | **Independent** | 0.307 | 0.978 | 0.719 | 0.874 | 0.415 | 0.906 |
| **XGB** | **Cross-validation** | 0.730 | 0.965 | 0.830 | 0.920 | 0.710 | 0.920 |
| **XGB** | **Independent** | 0.573 | 0.961 | 0.729 | 0.901 | 0.591 | 0.917 |
| **ET** | **Cross-validation** | 0.750 | 0.975 | 0.860 | 0.930 | 0.740 | 0.940 |
| **ET** | **Independent** | 0.480 | 0.980 | 0.818 | 0.903 | 0.580 | 0.949 |

(Abbreviations: SENS: Sensitivity; SPEC: Specificity; PPV: Positive Predictive Value; ACC: Accuracy; MCC: Matthews Correlation Coefficient; AUC: Area Under the Receiver Operating Characteristic Curve)

#### Other Dataset

##### Feature Name: AAC

| Classifier | Dataset | SENS | SPEC | PPV | ACC | MCC | AUC |
|---|---|---|---|---|---|---|---|
| **RF** | **Cross-validation** | 0.640 | 0.960 | 0.800 | 0.890 | 0.660 | 0.940 |
| **RF** | **Independent** | 0.594 | 0.966 | 0.811 | 0.893 | 0.634 | 0.919 |
| **GB** | **Cross-validation** | 0.700 | 0.950 | 0.780 | 0.890 | 0.690 | 0.940 |
| **GB** | **Independent** | 0.683 | 0.949 | 0.767 | 0.896 | 0.661 | 0.925 |
| **ElasticNet** | **Cross-validation** | 0.590 | 0.945 | 0.730 | 0.880 | 0.580 | 0.920 |
| **ElasticNet** | **Independent** | 0.554 | 0.949 | 0.727 | 0.871 | 0.560 | 0.904 |
| **Lasso regularization** | **Cross-validation** | 0.600 | 0.945 | 0.740 | 0.880 | 0.590 | 0.920 |
| **Lasso regularization** | **Independent** | 0.574 | 0.944 | 0.716 | 0.871 | 0.565 | 0.903 |
| **Ridge regularization** | **Cross-validation** | 0.610 | 0.950 | 0.750 | 0.880 | 0.590 | 0.920 |
| **Ridge regularization** | **Independent** | 0.574 | 0.946 | 0.725 | 0.873 | 0.571 | 0.906 |
| **LR** | **Cross-validation** | 0.610 | 0.950 | 0.750 | 0.880 | 0.590 | 0.920 |
| **LR** | **Independent** | 0.574 | 0.946 | 0.725 | 0.873 | 0.571 | 0.906 |
| **SVC (rbf)** | **Cross-validation** | 0.690 | 0.950 | 0.770 | 0.890 | 0.690 | 0.930 |
| **SVC (rbf)** | **Independent** | 0.673 | 0.956 | 0.791 | 0.900 | 0.670 | 0.916 |
| **DT** | **Cross-validation** | 0.660 | 0.920 | 0.670 | 0.860 | 0.580 | 0.900 |
| **DT** | **Independent** | 0.663 | 0.910 | 0.644 | 0.861 | 0.567 | 0.889 |
| **MLP** | **Cross-validation** | 0.630 | 0.930 | 0.680 | 0.870 | 0.570 | 0.800 |
| **MLP** | **Independent** | 0.554 | 0.929 | 0.659 | 0.855 | 0.518 | 0.742 |
| **AdB** | **Cross-validation** | 0.660 | 0.950 | 0.770 | 0.880 | 0.610 | 0.920 |
| **AdB** | **Independent** | 0.545 | 0.961 | 0.775 | 0.879 | 0.582 | 0.914 |
| **GNB** | **Cross-validation** | 0.740 | 0.940 | 0.740 | 0.890 | 0.640 | 0.920 |
| **GNB** | **Independent** | 0.713 | 0.942 | 0.750 | 0.896 | 0.667 | 0.925 |
| **XGB** | **Cross-validation** | 0.710 | 0.950 | 0.770 | 0.890 | 0.670 | 0.930 |
| **XGB** | **Independent** | 0.644 | 0.954 | 0.774 | 0.893 | 0.642 | 0.929 |
| **ET** | **Cross-validation** | 0.560 | 0.970 | 0.810 | 0.880 | 0.580 | 0.930 |
| **ET** | **Independent** | 0.505 | 0.971 | 0.797 | 0.871 | 0.544 | 0.933 |

(Abbreviations: SENS: Sensitivity; SPEC: Specificity; PPV: Positive Predictive Value; ACC: Accuracy; MCC: Matthews Correlation Coefficient; AUC: Area Under the Receiver Operating Characteristic Curve)

#### Other Dataset

##### Feature Name: DPC

| Classifier | Dataset | SENS | SPEC | PPV | ACC | MCC | AUC |
|---|---|---|---|---|---|---|---|
| **RF** | **Cross-validation** | 0.400 | 0.985 | 0.870 | 0.880 | 0.490 | 0.930 |
| **RF** | **Independent** | 0.377 | 0.990 | 0.889 | 0.857 | 0.478 | 0.926 |
| **GB** | **Cross-validation** | 0.530 | 0.940 | 0.670 | 0.860 | 0.500 | 0.910 |
| **GB** | **Independent** | 0.515 | 0.927 | 0.634 | 0.846 | 0.479 | 0.899 |
| **ElasticNet** | **Cross-validation** | 0.540 | 0.950 | 0.700 | 0.870 | 0.500 | 0.890 |
| **ElasticNet** | **Independent** | 0.455 | 0.946 | 0.676 | 0.850 | 0.471 | 0.871 |
| **Lasso regularization** | **Cross-validation** | 0.540 | 0.950 | 0.700 | 0.870 | 0.500 | 0.890 |
| **Lasso regularization** | **Independent** | 0.455 | 0.946 | 0.676 | 0.850 | 0.471 | 0.870 |
| **Ridge regularization** | **Cross-validation** | 0.550 | 0.955 | 0.710 | 0.875 | 0.510 | 0.900 |
| **Ridge regularization** | **Independent** | 0.465 | 0.951 | 0.701 | 0.855 | 0.492 | 0.878 |
| **LR** | **Cross-validation** | 0.550 | 0.955 | 0.710 | 0.875 | 0.510 | 0.900 |
| **LR** | **Independent** | 0.465 | 0.951 | 0.701 | 0.855 | 0.492 | 0.878 |
| **SVC (rbf)** | **Cross-validation** | 0.430 | 0.970 | 0.790 | 0.860 | 0.460 | 0.910 |
| **SVC (rbf)** | **Independent** | 0.307 | 0.978 | 0.775 | 0.846 | 0.423 | 0.904 |
| **GNB** | **Cross-validation** | 0.510 | 0.920 | 0.620 | 0.840 | 0.450 | 0.760 |
| **GNB** | **Independent** | 0.465 | 0.925 | 0.603 | 0.834 | 0.432 | 0.695 |
| **DT** | **Cross-validation** | 0.540 | 0.950 | 0.740 | 0.870 | 0.510 | 0.900 |
| **DT** | **Independent** | 0.475 | 0.959 | 0.738 | 0.863 | 0.519 | 0.899 |
| **MLP** | **Cross-validation** | 0.520 | 0.940 | 0.680 | 0.860 | 0.470 | 0.880 |
| **MLP** | **Independent** | 0.485 | 0.937 | 0.653 | 0.848 | 0.475 | 0.876 |
| **AdB** | **Cross-validation** | 0.550 | 0.945 | 0.700 | 0.870 | 0.490 | 0.895 |
| **AdB** | **Independent** | 0.495 | 0.944 | 0.685 | 0.855 | 0.500 | 0.910 |
| **XGB** | **Cross-validation** | 0.350 | 0.990 | 0.860 | 0.860 | 0.420 | 0.920 |
| **XGB** | **Independent** | 0.218 | 0.993 | 0.880 | 0.840 | 0.389 | 0.925 |
| **ET** | **Cross-validation** | 0.400 | 0.985 | 0.870 | 0.880 | 0.490 | 0.930 |
| **ET** | **Independent** | 0.377 | 0.990 | 0.889 | 0.857 | 0.478 | 0.926 |

(Abbreviations: SENS: Sensitivity; SPEC: Specificity; PPV: Positive Predictive Value; ACC: Accuracy; MCC: Matthews Correlation Coefficient; AUC: Area Under the Receiver Operating Characteristic Curve)

#### Other Dataset

##### Feature Name: AAC+DPC

| Classifier | Dataset | SENS | SPEC | PPV | ACC | MCC | AUC |
|---|---|---|---|---|---|---|---|
| **RF** | **Cross-validation** | 0.590 | 0.975 | 0.910 | 0.910 | 0.720 | 0.945 |
| **RF** | **Independent** | 0.476 | 0.978 | 0.809 | 0.859 | 0.488 | 0.937 |
| **GB** | **Cross-validation** | 0.720 | 0.950 | 0.820 | 0.930 | 0.740 | 0.930 |
| **GB** | **Independent** | 0.663 | 0.939 | 0.728 | 0.885 | 0.625 | 0.922 |
| **ElasticNet** | **Cross-validation** | 0.690 | 0.950 | 0.830 | 0.920 | 0.740 | 0.920 |
| **ElasticNet** | **Independent** | 0.545 | 0.961 | 0.775 | 0.879 | 0.582 | 0.890 |
| **Lasso regularization** | **Cross-validation** | 0.700 | 0.950 | 0.820 | 0.920 | 0.740 | 0.920 |

| | | | | | | | |
|---|---|---|---|---|---|---|---|
| **Lasso regularization** | **Independent** | 0.545 | 0.966 | 0.797 | 0.883 | 0.595 | 0.891 |
| **Ridge regularization** | **Cross-validation** | 0.690 | 0.950 | 0.830 | 0.920 | 0.740 | 0.920 |
| **Ridge regularization** | **Independent** | 0.505 | 0.956 | 0.739 | 0.867 | 0.537 | 0.886 |
| **LR** | **Cross-validation** | 0.690 | 0.950 | 0.830 | 0.920 | 0.740 | 0.920 |
| **LR** | **Independent** | 0.505 | 0.956 | 0.739 | 0.867 | 0.537 | 0.886 |
| **SVC (rbf)** | **Cross-validation** | 0.670 | 0.950 | 0.810 | 0.920 | 0.730 | 0.920 |
| **SVC (rbf)** | **Independent** | 0.584 | 0.956 | 0.766 | 0.883 | 0.602 | 0.924 |
| **GNB** | **Cross-validation** | 0.640 | 0.920 | 0.730 | 0.900 | 0.670 | 0.880 |
| **GNB** | **Independent** | 0.515 | 0.915 | 0.598 | 0.836 | 0.455 | 0.833 |
| **DT** | **Cross-validation** | 0.650 | 0.925 | 0.740 | 0.900 | 0.670 | 0.850 |
| **DT** | **Independent** | 0.584 | 0.922 | 0.648 | 0.855 | 0.527 | 0.753 |
| **MLP** | **Cross-validation** | 0.720 | 0.950 | 0.860 | 0.930 | 0.750 | 0.910 |
| **MLP** | **Independent** | 0.525 | 0.959 | 0.757 | 0.873 | 0.560 | 0.910 |
| **AdB** | **Cross-validation** | 0.720 | 0.950 | 0.850 | 0.930 | 0.750 | 0.920 |
| **AdB** | **Independent** | 0.673 | 0.956 | 0.791 | 0.900 | 0.670 | 0.934 |
| **XGB** | **Cross-validation** | 0.730 | 0.955 | 0.830 | 0.935 | 0.760 | 0.940 |
| **XGB** | **Independent** | 0.683 | 0.942 | 0.742 | 0.891 | 0.645 | 0.925 |
| **ET** | **Cross-validation** | 0.570 | 0.985 | 0.930 | 0.910 | 0.660 | 0.950 |
| **ET** | **Independent** | 0.267 | 0.988 | 0.844 | 0.846 | 0.419 | 0.933 |

(Abbreviations: SENS: Sensitivity; SPEC: Specificity; PPV: Positive Predictive Value; ACC: Accuracy; MCC: Matthews Correlation Coefficient; AUC: Area Under the Receiver Operating Characteristic Curve)

#### Other Dataset

#### Feature Name: ALLCOMP

| Classifier | Dataset | SENS | SPEC | PPV | ACC | MCC | AUC |
|---|---|---|---|---|---|---|---|
| **RF** | **Cross-validation** | 0.630 | 0.975 | 0.910 | 0.900 | 0.720 | 0.945 |
| **RF** | **Independent** | 0.545 | 0.978 | 0.859 | 0.893 | 0.629 | 0.946 |
| **GB** | **Cross-validation** | 0.700 | 0.965 | 0.890 | 0.930 | 0.740 | 0.940 |
| **GB** | **Independent** | 0.653 | 0.961 | 0.805 | 0.900 | 0.667 | 0.925 |
| **ElasticNet** | **Cross-validation** | 0.220 | 0.985 | 0.800 | 0.840 | 0.380 | 0.710 |
| **ElasticNet** | **Independent** | 0.277 | 0.995 | 0.933 | 0.854 | 0.461 | 0.681 |
| **Lasso regularization** | **Cross-validation** | 0.210 | 0.990 | 0.830 | 0.835 | 0.370 | 0.690 |
| **Lasso regularization** | **Independent** | 0.287 | 0.995 | 0.935 | 0.855 | 0.471 | 0.674 |
| **Ridge regularization** | **Cross-validation** | 0.310 | 0.970 | 0.770 | 0.840 | 0.430 | 0.830 |
| **Ridge regularization** | **Independent** | 0.347 | 0.971 | 0.745 | 0.848 | 0.437 | 0.849 |
| **LR** | **Cross-validation** | 0.310 | 0.970 | 0.770 | 0.840 | 0.430 | 0.830 |
| **LR** | **Independent** | 0.347 | 0.971 | 0.745 | 0.848 | 0.437 | 0.849 |
| **SVC (rbf)** | **Cross-validation** | 0.590 | 0.995 | 0.975 | 0.910 | 0.690 | 0.960 |

| | | | | | | | |
|---|---|---|---|---|---|---|---|
| **SVC (rbf)** | **Independent** | 0.356 | 0.993 | 0.923 | 0.867 | 0.524 | 0.932 |
| **GNB** | **Cross-validation** | 0.190 | 0.975 | 0.750 | 0.830 | 0.320 | 0.770 |
| **GNB** | **Independent** | 0.218 | 0.985 | 0.786 | 0.834 | 0.356 | 0.816 |
| **DT** | **Cross-validation** | 0.710 | 0.960 | 0.860 | 0.930 | 0.740 | 0.900 |
| **DT** | **Independent** | 0.446 | 0.966 | 0.763 | 0.863 | 0.513 | 0.841 |
| **MLP** | **Cross-validation** | 0.690 | 0.940 | 0.760 | 0.910 | 0.710 | 0.830 |
| **MLP** | **Independent** | 0.634 | 0.937 | 0.711 | 0.877 | 0.596 | 0.785 |
| **AdB** | **Cross-validation** | 0.700 | 0.970 | 0.880 | 0.930 | 0.740 | 0.910 |
| **AdB** | **Independent** | 0.436 | 0.976 | 0.815 | 0.869 | 0.533 | 0.915 |
| **XGB** | **Cross-validation** | 0.730 | 0.965 | 0.890 | 0.930 | 0.750 | 0.930 |
| **XGB** | **Independent** | 0.743 | 0.946 | 0.773 | 0.906 | 0.700 | 0.940 |
| **ET** | **Cross-validation** | 0.710 | 0.965 | 0.870 | 0.930 | 0.740 | 0.940 |
| **ET** | **Independent** | 0.663 | 0.959 | 0.798 | 0.900 | 0.668 | 0.941 |

(Abbreviations: SENS: Sensitivity; SPEC: Specificity; PPV: Positive Predictive Value; ACC: Accuracy; MCC: Matthews Correlation Coefficient; AUC: Area Under the Receiver Operating Characteristic Curve)

#### *Supplementary Table S6: Performance of PLM-based models on all four datasets.*

| **Dataset** | **PLM Classifier** | **Coss-validation Data** | | | | | | **Independent Data** | | | | | |
|---|---|---|---|---|---|---|---|---|---|---|---|---|---|
| | | **SENS** | **SPEC** | **PPV** | **ACC** | **MCC** | **AUC** | **SENS** | **SPEC** | **PPV** | **ACC** | **MCC** | **AUC** |
| **Na+** | ESM2-t6 | 0.617 | 0.986 | 0.926 | 0.890 | 0.672 | 0.955 | 0.594 | 0.978 | 0.894 | 0.887 | 0.668 | 0.951 |
| | ESM2-t12 | 0.655 | 0.981 | 0.976 | 0.917 | 0.719 | 0.966 | 0.617 | 0.988 | 0.941 | 0.900 | 0.712 | 0.969 |
| | ESM2-t30 | 0.709 | 0.996 | 0.964 | 0.959 | 0.729 | 0.960 | 0.633 | 0.985 | 0.931 | 0.902 | 0.715 | 0.959 |
| | ESM2-t33 | **0.803** | **0.994** | **0.980** | **0.960** | **0.826** | **0.988** | **0.788** | **0.983** | **0.933** | **0.930** | **0.799** | **0.982** |
| | ESM2-t36 | 0.775 | 0.988 | 0.960 | 0.966 | 0.835 | 0.975 | 0.742 | 0.988 | 0.950 | 0.930 | 0.799 | 0.966 |
| | bert-base-uncased | 0.792 | 0.966 | 0.906 | 0.799 | 0.809 | 0.941 | 0.789 | 0.917 | 0.887 | 0.694 | 0.768 | 0.934 |
| | ProtBert | 0.981 | 0.970 | 0.909 | 0.972 | 0.926 | 0.963 | 0.875 | 0.956 | 0.862 | 0.937 | 0.827 | 0.955 |
| | distilbert-base-uncased | 0.737 | 0.927 | 0.760 | 0.893 | 0.681 | 0.832 | 0.708 | 0.853 | 0.605 | 0.817 | 0.528 | 0.847 |
| **K+** | ESM2-t6 | 0.765 | 0.988 | 0.954 | 0.953 | 0.786 | 0.977 | 0.716 | 0.981 | 0.907 | 0.925 | 0.763 | 0.959 |
| | ESM2-t12 | 0.780 | 0.973 | 0.885 | 0.930 | 0.790 | 0.958 | 0.752 | 0.978 | 0.901 | 0.930 | 0.780 | 0.960 |
| | ESM2-t30 | 0.935 | 0.989 | 0.957 | 0.955 | 0.785 | 0.969 | 0.734 | 0.968 | 0.860 | 0.919 | 0.746 | 0.963 |
| | ESM2-t33 | **0.852** | **0.985** | **0.932** | **0.952** | **0.838** | **0.973** | **0.829** | **0.976** | **0.896** | **0.947** | **0.812** | **0.971** |
| | ESM2-t36 | 0.801 | 0.991 | 0.945 | 0.953 | 0.832 | 0.971 | 0.771 | 0.981 | 0.913 | 0.937 | 0.801 | 0.966 |

| | | | | | | | | | | | | | |
|---|---|---|---|---|---|---|---|---|---|---|---|---|---|
| | bert-base-uncased | 0.494 | 0.998 | 0.876 | 0.526 | 0.539 | 0.915 | 0.449 | 0.983 | 0.859 | 0.489 | 0.494 | 0.904 |
| | ProtBert | 0.649 | 0.999 | 0.925 | 0.918 | 0.648 | 0.980 | 0.587 | 0.985 | 0.914 | 0.902 | 0.683 | 0.962 |
| | distilbert-base-uncased | 0.698 | 0.933 | 0.738 | 0.881 | 0.631 | 0.873 | 0.561 | 0.915 | 0.604 | 0.845 | 0.499 | 0.881 |
| **Ca+** | ESM2-t6 | 0.983 | 0.993 | 0.836 | 0.939 | 0.687 | 0.940 | 0.627 | 0.971 | 0.797 | 0.918 | 0.661 | 0.934 |
| | ESM2-t12 | 0.709 | 0.994 | 0.866 | 0.979 | 0.773 | 0.973 | 0.680 | 0.985 | 0.895 | 0.938 | 0.747 | 0.945 |
| | ESM2-t30 | 0.563 | 0.994 | 0.916 | 0.940 | 0.645 | 0.938 | 0.493 | 0.990 | 0.902 | 0.913 | 0.629 | 0.927 |
| | ESM2-t33 | **0.753** | **0.976** | **0.859** | **0.942** | **0.771** | **0.950** | **0.733** | **0.981** | **0.873** | **0.942** | **0.768** | **0.955** |
| | ESM2-t36 | 0.706 | 0.991 | 0.919 | 0.957 | 0.766 | 0.960 | 0.667 | 0.985 | 0.893 | 0.936 | 0.738 | 0.951 |
| | bert-base-uncased | 0.690 | 0.993 | 0.918 | 0.606 | 0.663 | 0.912 | 0.653 | 0.937 | 0.893 | 0.590 | 0.653 | 0.900 |
| | ProtBert | 0.873 | 0.931 | 0.727 | 0.921 | 0.747 | 0.942 | 0.840 | 0.929 | 0.685 | 0.916 | 0.709 | 0.925 |
| | distilbert-base-uncased | 0.722 | 0.950 | 0.754 | 0.874 | 0.621 | 0.871 | 0.653 | 0.910 | 0.628 | 0.831 | 0.541 | 0.881 |
| **Other** | ESM2-t6 | 0.963 | 0.994 | 0.975 | 0.928 | 0.752 | 0.965 | 0.644 | 0.981 | 0.890 | 0.914 | 0.710 | 0.953 |
| | ESM2-t12 | 0.947 | 0.994 | 0.975 | 0.983 | 0.857 | 0.972 | 0.772 | 0.985 | 0.929 | 0.943 | 0.814 | 0.951 |
| | ESM2-t30 | 0.643 | 0.993 | 0.929 | 0.933 | 0.757 | 0.968 | 0.614 | 0.985 | 0.912 | 0.912 | 0.703 | 0.945 |
| | ESM2-t33 | **0.785** | **0.993** | **0.930** | **0.950** | **0.836** | **0.975** | **0.831** | **0.985** | **0.949** | **0.955** | **0.854** | **0.957** |
| | ESM2-t36 | 0.786 | 0.991 | 0.978 | 0.961 | 0.831 | 0.966 | 0.753 | 0.990 | 0.950 | 0.943 | 0.814 | 0.947 |
| | bert-base-uncased | 0.631 | 0.995 | 0.817 | 0.919 | 0.682 | 0.926 | 0.624 | 0.964 | 0.808 | 0.897 | 0.650 | 0.918 |
| | ProtBert | 0.743 | 0.997 | 0.938 | 0.941 | 0.798 | 0.827 | 0.733 | 0.985 | 0.925 | 0.936 | 0.787 | 0.816 |
| | distilbert-base-uncased | 0.693 | 0.915 | 0.826 | 0.844 | 0.639 | 0.832 | 0.627 | 0.918 | 0.598 | 0.825 | 0.513 | 0.827 |

(Abbreviations: SENS: Sensitivity; SPEC: Specificity; PPV: Positive Predictive Value; ACC: Accuracy; MCC: Matthews Correlation Coefficient; AUC: Area Under the Receiver Operating Characteristic Curve; Note: The bold values indicate the best-performing models for each dataset)
